# Leveraging a machine learning derived surrogate phenotype to improve power for genome-wide association studies of partially missing phenotypes in population biobanks

**DOI:** 10.1101/2022.12.12.520180

**Authors:** Zachary R. McCaw, Jianhui Gao, Xihong Lin, Jessica Gronsbell

## Abstract

Within population biobanks, genetic discovery for specialized phenotypes is often limited by incomplete ascertainment. Machine learning (ML) is increasingly used to impute missing phenotypes from surrogate information. However, imputing missing phenotypes can invalidate statistical inference when the imputation model is misspecified, and proxy analysis of the ML-phenotype can introduce spurious associations. To overcome these limitations, we introduce SynSurr, an approach that jointly analyzes a partially missing target phenotype with a “synthetic surrogate”, its predicted value from an ML-model. SynSurr estimates the same genetic effect as standard genome-wide association studies (GWAS) of the target phenotype, but improves power provided the synthetic surrogate is correlated with the target. Unlike imputation or proxy analysis, SynSurr does not require that the synthetic surrogate is obtained from a correctly specified generative model. We perform extensive simulations and an ablation analysis to compare SynSurr with existing methods. We also apply SynSurr to empower GWAS of dual-energy x-ray absorptiometry traits within the UK Biobank, leveraging a synthetic surrogate composed of bioelectrical impedance and anthropometric traits.

## Introduction

Over the last decade, there has been immense interest in conducting genomic research in population biobanks such as FinnGenn (*>* 200,000 individuals), the Million Veteran Program (*>* 500,000 veterans), and the UK Biobank (*>* 400,000 individuals)^1–3^. While these deeply phenotyped resources have created an unprecedented opportunity to increase both the scope and precision of genome-wide association studies (GWAS), they have simultaneously introduced new statistical challenges^4,5^. Among these challenges, incompletely measured or partially missing phenotypes are a key issue^6^. Missingness often arises due to the difficulty, expense, or invasiveness of ascertaining the target phenotype^7^. Examples of partially missing phenotypes from the UK Biobank (UKBB)^8^ include body composition phenotypes obtained from dual-energy x-ray absorptiometry (DEXA) scans^9^, neurological^10^ and cardiac^11^ structural features extracted from functional magnetic resonance imaging, optic morphology parameters extracted from retinal fundus images^12^, and sleep wake patterns extracted from wearable accelerometry trackers^13^. Each of these phenotypes was ascertained, at least initially, in only a subset of the full cohort. Restricting the analysis to only those individuals with observed target phenotypes would substantially diminish statistical power. To address this practical and statistical issue, we propose a novel method for conducting large-scale GWAS of a partially missing target phenotype that improves power by leveraging a machine learning (ML) derived surrogate phenotype.

As biobanks typically contain extensive demographic and clinical information from baseline assessments and electronic health records, an ML-model can often be developed to reliably predict the partially missing phenotype, potentially increasing power for subsequent GWAS^14–16^. In contrast to the target phenotype, which is only observed for a subset of the cohort, the ML-derived phenotype is available for the entire cohort, enabling GWAS with the full data set. There are now several approaches that can leverage ML-based predictions in performing GWAS^17,18^. The simplest approach is to perform proxy GWAS, using the predicted phenotype as the outcome of interest for all subjects. Proxy GWAS performs well if the prediction model is highly accurate. However, if the predictions are imperfect, then genetic variants associated with the ML-phenotype may not be relevant to the target phenotype. Additionally, the estimated genetic effects will not reflect the effect of genotype on the target phenotype. Moreover, proxy GWAS may forego power by disregarding the fact that the target phenotype was actually observed for a subset of the cohort^19,20^.

Another simple approach is to retain the observed values of the target phenotype, impute the missing values from an ML-model, then perform standard GWAS analysis using the pooled observed and imputed target phenotypes. Although simple to implement, inference based on single-imputation is generally invalid because it ignores imputation uncertainty by treating imputed values as if they were the actual values^18,21–23^. Specifically, single-imputation regards the observed and imputed values as originating from the same distribution, when in fact the imputed values originate from a different distribution and are less variable than their observed counterparts. In the context of linear regression, it is well known that failing to account for the reduced variability introduced by imputation will lead to systematic under-estimation of the residual variance and consequent inflation of the type I error^24^. Multiple imputation (MI)^25–27^ has been proposed to overcome the limitations of single imputation. MI generates several imputations of the missing values from a probabilistic model, performs an analysis with each set of imputed values, then combines the resulting estimates, via Rubin’s Rules, to correctly account for imputation uncertainty. The major limitation of MI is that the imputations must come from a correctly specified model^27–30^. When the imputation model is misspecified, as is likely in practice, the estimates resulting from MI are biased.

To overcome the limitations of imputation and proxy analysis, we propose synthetic surrogate analysis, or “SynSurr” for short, which improves power for GWAS of a partially missing target phenotype. Within a model-building data set, we first train an ML-model for predicting the target phenotype on the basis of readily available surrogate data. The model is then transferred to the GWAS data set, where it is applied to generate a “synthetic surrogate” phenotype for all subjects. The partially observed target phenotype and fully observed synthetic surrogate are then *jointly* analyzed within a bivariate outcome framework. SynSurr falls within the previously described surrogate phenotype regression framework^31^, but the estimation and inference procedures described here have been specialized and accelerated for application at biobank scale. Unlike proxy GWAS, SynSurr always estimates the same genetic effect as a standard GWAS. SynSurr improves power to discover variants specifically associated with the target phenotype by leveraging the correlation between the target phenotype and its synthetic surrogate to enhance precision. Finally, unlike imputation-based inference, SynSurr does not require the strong and often untenable assumption that the ML-model which generates the synthetic surrogate is correctly specified. In fact, SynSurr remains valid and comparably powerful to standard GWAS even when the synthetic surrogate is completely uncorrelated with the target phenotype.

We illustrate the methodological advantages and practical implementation of SynSurr through extensive analyses of simulated and real data. Our simulation studies illustrate three key properties of SynSurr, which are backed by theoretical analyses provided in the Supplementary Methods. First, SynSurr is *robust* in the sense that it is no less powerful than standard GWAS even when the synthetic surrogate is uncorrelated with the target phenotype. Second, SynSurr is more *powerful* than standard GWAS when the synthetic surrogate is correlated with the target phenotype. The power advantage increases with target missingness and the target-surrogate correlation. Third, SynSurr always provides *valid inference*, meaning that the estimated genetic effects are unbiased and the type I error is properly controlled, even when the ML-model that generates the synthetic surrogate is misspecified (i.e. differs from the generative model for the target phenotype).

Our real data analyses of phenotypes from the UK Biobank (UKBB) include an ablation analysis of two phenotypes with minimal missingness, height and forced expiratory volume in 1 second (FEV1), and an application of SynSurr to 6 body composition phenotypes, measured by DEXA, with substantial missingness. The ablation analysis demonstrates that SynSurr and standard GWAS identify all the same variants in the absence of missingness, but that SynSurr is uniformly more powerful in the presence of missingness. SynSurr achieves this power advantage not by inflating the false discovery rate or distorting the estimated genetic effect, but by leveraging the surrogate outcome to obtain more precise estimation (i.e. smaller standard errors). The application to DEXA phenotypes demonstrates the substantial opportunity for improved power with SynSurr. Compared to standard GWAS using only the partially observed phenotypes, SynSurr identified on average 21.5 times as many genome-wide significant variants, and did so at 3.3 times the level of significance. Moreover, the variants identified by SynSurr are relevant to the phenotypes under study, being significantly enriched for gene sets related to body composition, and overlapping substantially with GWAS Catalog^32^ variants previously associated with body composition traits.

## Results

### Overview of Method

**Figure 1** provides a graphical overview of a SynSurr GWAS. The overarching goal is to utilize surrogate information to improve power for GWAS of a partially missing target phenotype. The complete data set is first split into model-building and inference data sets. The model-building data set includes the target phenotype and a set of surrogates, but does not require genetics. For exposition, we suppose the model that generates the synthetic surrogate is trained via supervised learning, however this is not prescriptive, and other approaches (e.g. semi-supervised learning) may be employed. In the supervised case, the model-building data set is fully labeled, meaning the target outcome is available for all subjects. The inference data set, in which GWAS is performed, requires genetics and surrogates, and is assumed to be only partially labeled, meaning the target outcome is partially missing.

**Figure 1:**
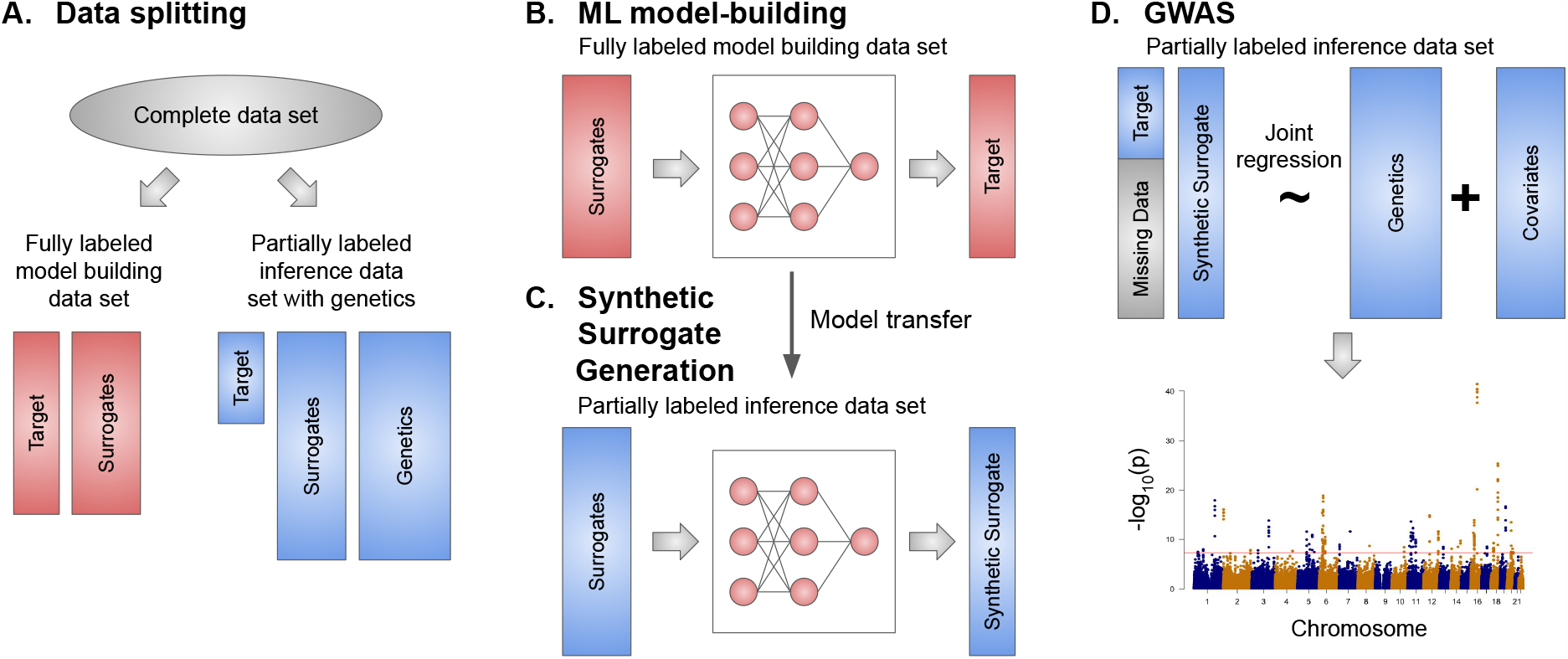
Graphical overview of a SynSurr GWAS. **A**. The complete data set is first split into a fully labeled model-building data set, including the target phenotype and surrogates, and a partially labeled inference data set, which also includes genetics. **B**. Within the fully labeled model-building data set, an ML-model is trained to predicted the target phenotype on the basis of surrogates. **C**. The ML-model is transferred to the partially labeled inference data set and applied to predict the target outcome for all subjects. The predicted value of the target outcome is referred to as the “synthetic surrogate”. Importantly, the synthetic surrogate is maintained as a separate and distinct outcome from the partially missing target phenotype. **D**. Finally, within the inference data set, the partially missing target phenotype and the fully observed synthetic surrogate are jointly regressed on genotype and covariates to identify genetic variants associated with the target outcome.

Within the model-building data set, an ML-model is trained to predict the target phenotype on the basis of surrogate information. The trained model is then transferred to the inference data set, where it is used to predict the target phenotype for *all* subjects (not only those with missing phenotypes). Because the ML-model synthesizes multiple surrogates to predict the target phenotype, we refer to its output as a *synthetic surrogate*. In contrast to imputation, the synthetic surrogate is maintained as a separate and distinct outcome from the partially missing target phenotype. Finally, within the inference data set, the partially missing target phenotype and the fully observed synthetic surrogate are jointly regressed on genotype and covariates to identify genetic variants associated with the target outcome.

In contrast to standard GWAS, which can only incorporate subjects whose target phenotype is observed, SynSurr improves power by allowing subjects whose target phenotypes are missing to enter the analysis. Critically, unlike imputation-based inference, SynSurr does not require that the synthetic surrogate is generated from a correctly specified imputation model, as the synthetic surrogate is never used as a replacement for the target outcome. SynSurr is robust, remaining valid and not foregoing power even if the synthetic surrogate is uncorrelated with the target phenotype. Moreover, SynSurr improves power to the extent that the target and synthetic surrogate outcomes are correlated. As we will show, SynSurr provides an unbiased estimate for the effect of genotype on the target outcome, increases estimation precision, and improves power for identifying variants specifically associated with the target phenotype.

The Methods section provides a mathematical description of the SynSurr model and an overview of the estimation procedure. The Supplementary Methods provide a detailed derivation of maximum likelihood estimates for all model parameters, their standard errors, and a Wald test for evaluating the association between genotype and the target phenotype.

### Generation of synthetic surrogates

SynSurr depends on the availability of a synthetic surrogate, *Ŷ*, that is constructed for all subjects and ideally correlated with the phenotype of interest, *Y* . We focus on the setting where the surrogate is a prediction of *Y* from an ML-model. This setting is particularly relevant in population biobanks where certain phenotypes are too invasive, expensive, or time-consuming to measure for the entire cohort, but can be reliably predicted from a model trained on the available surrogate data (e.g. information from electronic health records, administrative data, or baseline assessments). The inputs to the model that generates the synthetic surrogate (“the surrogate model”) should not include genetic data *G*, and may or may not include covariates *X* that are also adjusted for during the GWAS. In training a prediction model to generate the synthetic surrogate, the goal is to obtain a prediction *Ŷ*that is highly correlated with the target phenotype *Y* . More specifically, *Y* and *Ŷ* should be highly correlated after adjusting for the covariates *X* included in the GWAS. As detailed in the Supplementary Methods section, a stronger residual correlation leads to increased power. To capture the potentially complex relationship between a set of covariates and the phenotype of interest, we recommend that the synthetic surrogate be generated from a nonlinear ML-model. For structured data settings with well-defined covariates, tree-based models such as random forest^33^ or Extreme Gradient Boosting (XGBoost)^34^ generally perform well and are straightforward to train with standard software. Neural network models can also be used and are particularly advantageous for deriving synthetic surrogates from unstructured data such as images or free-text.

### SynSurr overcomes the pitfalls of imputation-based inference

Recall that our goal is to perform inference on the association between *Y* and *G* when the outcome *Y* is only partially observed, but a synthetic surrogate *Ŷ*is available for all subjects. The simplest approach that incorporates all subjects in the analysis is to perform a proxy GWAS for *Ŷ* in place of *Y* . This approach, however, changes the research question from performing inference on the association between *Y* and *G* (i.e. *β*_*G*_ in Equation 5) to performing inference on the association between *Ŷ* and *G* (i.e. *α*_*G*_ in Equation 5). Moreover, analyzing *Ŷ* instead of *Y* can reduce power or lead to spurious associations if *Ŷ* is an imperfect proxy for *Y* . To preserve the original research question, an alternative approach is to generate a new outcome *Y* ^*\**^ by imputing missing values of *Y* with *Ŷ* :

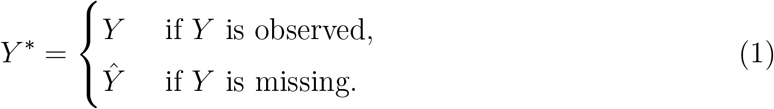

Single imputation is generally invalid, as the analytical standard errors will be underestimated^24,27^. To perform valid inference after imputation, multiple imputations of the missing data can be generated, and the resulting point estimates and standard errors combined using Rubin’s rules^24,27^. A major limitation of multiple imputation, as demonstrated below, is that its validity requires a correctly specified imputation model. A key contribution of the the present work, with ramifications beyond GWAS, is to provide a framework for utilizing *Ŷ* to improve inference on the association between *Y* and *G without* requiring that *Ŷ* be generated from a correctly specified imputation model.

As an example of the pitfalls that can arise when performing imputation-based inference, we generated phenotypes from the following model:

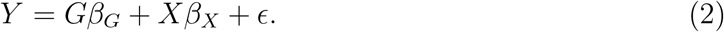

Here *Y* represents the target phenotype, *G* the genotype, *X* a correlated covariate, and *ϵ* a residual. The true genetic effect is *β*_*G*_ = 0.1, corresponding to a variant with *h*^2^ = 1%. The total sample size is *n* = 10^4^ and 25% of the values of *Y* are missing. We compare the performance of an *oracle estimator*, which has access to *Y* before the introduction of missingness, the *standard estimator*, which only has access to the observed values of *Y*, single imputation (SI), multiple imputation (MI), and the proposed SynSurr estimator. The imputation models are as follows:

1. A correctly specified model, which imputes *Y* based on *G* and *X*.
2. A misspecified model, which imputes *Y* using *G* only.
3. A misspecified model, which imputes *Y* using *X* only.

Each imputation model was fit on an independent model-building data set of size 10^3^ then applied to generate *Ŷ* for all subjects in the inference data set. The SI and MI estimators used *Y* ^*\**^ as their outcome, constructed via Equation (1), while the corresponding SynSurr estimator regarded *Ŷ* as the synthetic surrogate. All estimators utilized Equation (2) as the association model, allowing for direct comparison of the resulting estimates of *β*_*G*_.

**Figure 2** and **Supplementary Table 1** present the point estimates and analytic and empirical standard errors (SEs) of the imputation-based estimators compared to SynSurr. The oracle estimator (green) is unbiased for *β*_*G*_, and as a correctly specified maximum likelihood estimator, its SE represents the best that is attainable in the absence of missing data^35^. The standard estimator (orange) is also unbiased, and because observations with missing outcomes do not contribute to this estimator, its SE is larger than that of oracle. The SI (red) and MI (blue) estimators based on the correctly specified imputation model are unbiased. While the 95% confidence interval (CI) for the MI estimator has proper coverage, as indicated by the agreement between the analytical CI (dotted) and the empirical CI (solid), the analytical CI for the SI estimator falls short of the empirical CI. This occurs because the SE of the SI estimator is underestimated. Consequently, inference based on the SI estimator would lead to an overstatement of significance (i.e. inflated type I error)^21^. Estimates based on an incorrectly specified imputation model are biased, whether the the misspecification was due to omission of the variable of interest (i.e. *G*) or simply a correlated covariate (i.e. *X*)^26^. In contrast, the synthetic surrogate estimator (purple) is unbiased regardless of what collection of covariates is used to generate *Ŷ* . Because *Ŷ* was generated from a linear model, and the covariates used to generate *Ŷ* are conditioned on the GWAS model, the SynSurr estimator is not appreciably more efficient than the standard estimator in this simulation.

**Figure 2:**
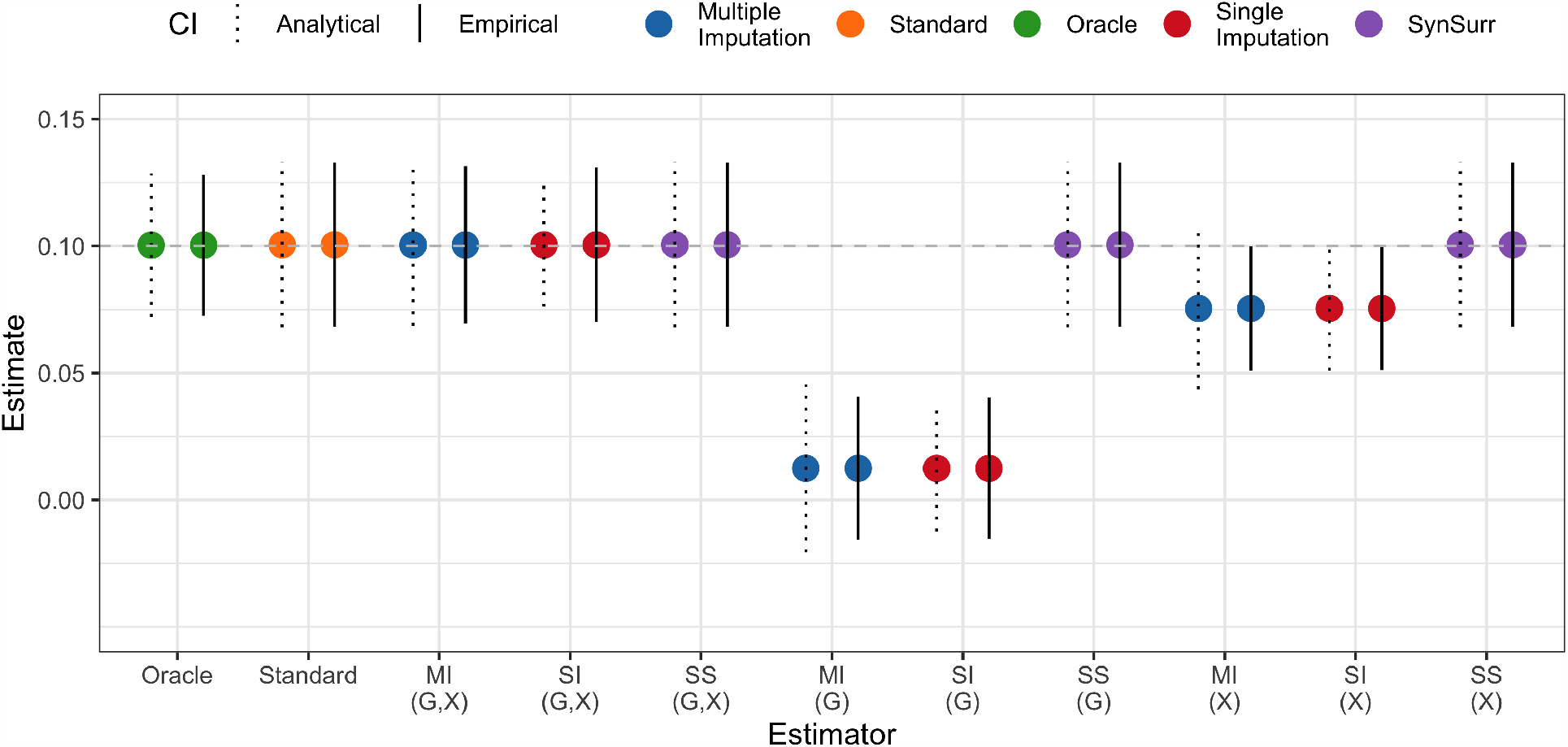
Unlike imputation-based estimators, SynSurr is robust to misspecification of the imputation model. The true value for the parameter of interest is *β*_*G*_ = 0.1, corresponding to a variant with *h*^2^ = 1%. For each estimator, the mean value across 10^3^ simulations is shown by the point, and two 95% confidence intervals (CIs) are presented: the dotted CI is based on the root-mean-square analytical standard error (SE) while the solid CI is based on the empirical SE. The oracle estimator has access to the complete version of *Y*, before 25% of values were set to missing. The standard estimator has access to the observed values of *Y* only. The imputation-based estimators (SI: single imputation, MI: multiple imputation) impute the missing values of *Y* using an imputation-model fit on an independent data set. The set of covariates used to fit the imputation model are shown as a tuple: the imputation model based on *G* and *X* is correctly specified, whereas that based on *G* alone or *X* alone is misspecified. The SynSurr estimator (SS) jointly analyzes the partially missing *Y* with the synthetic surrogate *Ŷ*, where *Ŷ* is generated for all subjects from the imput tion model. The key observation is that SS does not require a correctly specified generative model to yield unbiased estimation and valid inference.

In practice, to improve the surrogacy of *Ŷ* and allow the SynSurr estimator to realize a power improvement, *Ŷ* should be generated from a non-linear model and should incorporate predictors other than those adjusted for in the GWAS model.

### SynSurr is robust to uninformative surrogates and yields more precise inference with informative surrogates

Unlike imputation-based inference, which is sensitive to correct specification of the imputation model, SynSurr is robust to the choice of synthetic surrogate in that it (i) consistently estimates the effect of the genotype on the target phenotype regardless of how well the surrogate predicts the target phenotype and (ii) provides improved power over standard GWAS when the synthetic surrogate is correlated with the target phenotype. To demonstrate these points, we again simulated phenotypes from Equation (2) and varied the proportion of subjects with missing phenotypes and the correlation between *Y* and *Ŷ* (see Supplementary Section 3.1). **Supplementary Figure 1** presents box plots of 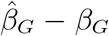for the standard GWAS estimator and the SynSurr estimator under different levels of missingness in *Y* . In panel A, the synthetic surrogate *Ŷ* is completely uncorrelated with, and in fact independent of, the phenotype of interest *Y* . Nevertheless, the SynSurr estimator is unbiased and no less efficient than the standard GWAS estimator, as indicated by consistent widths of the box plots. This demonstrates that SynSurr is robust to the use of an uninformative surrogate. However, as demonstrated in panel B, when *Ŷ* is correlated with *Y*, the precision of the SynSurr estimator increases with the number of subjects with missing phenotypes (also see **Supplementary Figure 2**). These qualitative observations are supported by a quantitative comparison of standard errors in **Supplementary Table 2** as well as a theoretical analysis (see Supplementary Section 1.4).

### SynSurr controls type I error and increases power relative to standard GWAS

Building on the findings of the previous section, we again simulated phenotypes from Equation (2) to evaluate the type I error and power of SynSurr across various missing rates, choices of synthetic surrogate, and levels of SNP heritability. Here type I error refers to the probability of incorrectly rejecting *H*_0_ : *β*_*G*_ = 0 when *β*_*G*_ = 0 and power refers to the probability of correctly rejecting *H*_0_ when *β*_*G*_ ≠ 0. **Figure 3** demonstrates that, across missingness rates and target-surrogate correlations, the SynSurr p-values are uniformly distributed under the null hypothesis of no association. **Supplementary Table 3** presents the type I error and power at a SNP heritability of *h*^2^ = 0.5% (*β*_*G*_ *≈* 0.07), together with the average *χ*^2^ statistics. The type I error is consistently controlled across missing rates and choices of the synthetic surrogate. Consonant with the previous section and with the efficiency analysis in the Supplementary Methods, **Supplementary Table 3** demonstrates that the power of SynSurr increases with the target-synthetic surrogate correlation and level of target missingness. For instance, when the missingness rate is 90% and the correlation between the synthetic surrogate and the target phenotype is 0.75, there is a 27% increase in power relative to the standard analysis at the same missing rate. Moreover, **Figure 4** illustrates the benefit of SynSurr with respect to power across SNP heritabilities ranging from 0.1% to 1.0%. As the proportion of subjects with missing target phenotypes increases, the benefit of SynSurr with a well-correlated surrogate phenotype is increasingly apparent. Interestingly, the relative efficiency of SynSurr does not depend on the SNP heritability (**Supplementary Figure 3**).

**Figure 3:**
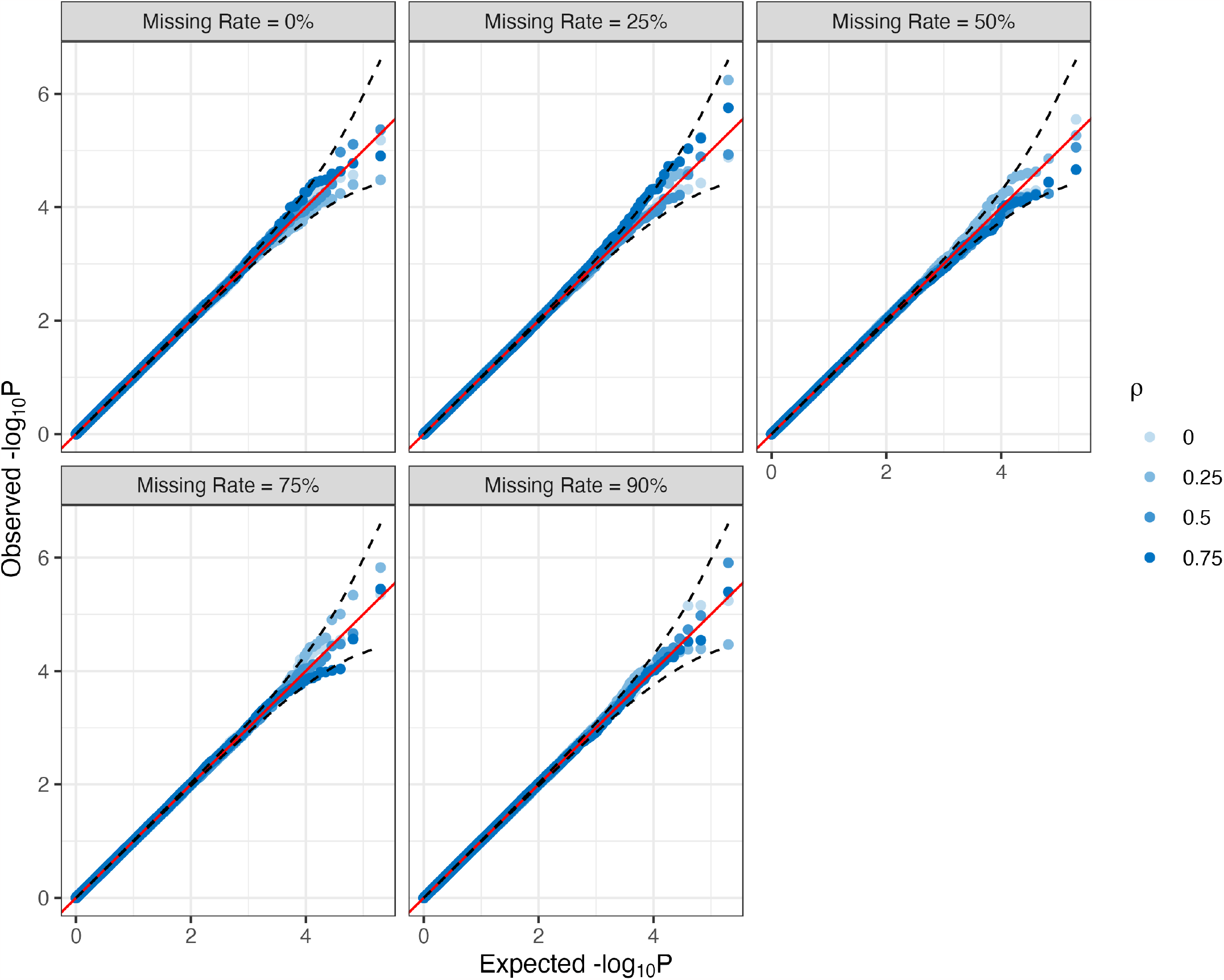
Uniform quantile-quantile plots for the SynSurr p-values under the null hypothesis of no association across various missingness rates and target-surrogate correlations. In all cases, the number of subjects with observed phenotypes was *n* = 10^3^. The number of subjects with missing phenotypes was varied to achieve the indicated level of missingness. The synthetic surrogate has correlation *ρ ∈ {*0.00, 0.25, 0.50, 0.75*}* with the target phenotype. The number of simulation replicates is 10^6^ . Adherence to the diagonal indicates that the p-values are uniformly distributed under the null.

**Figure 4:**
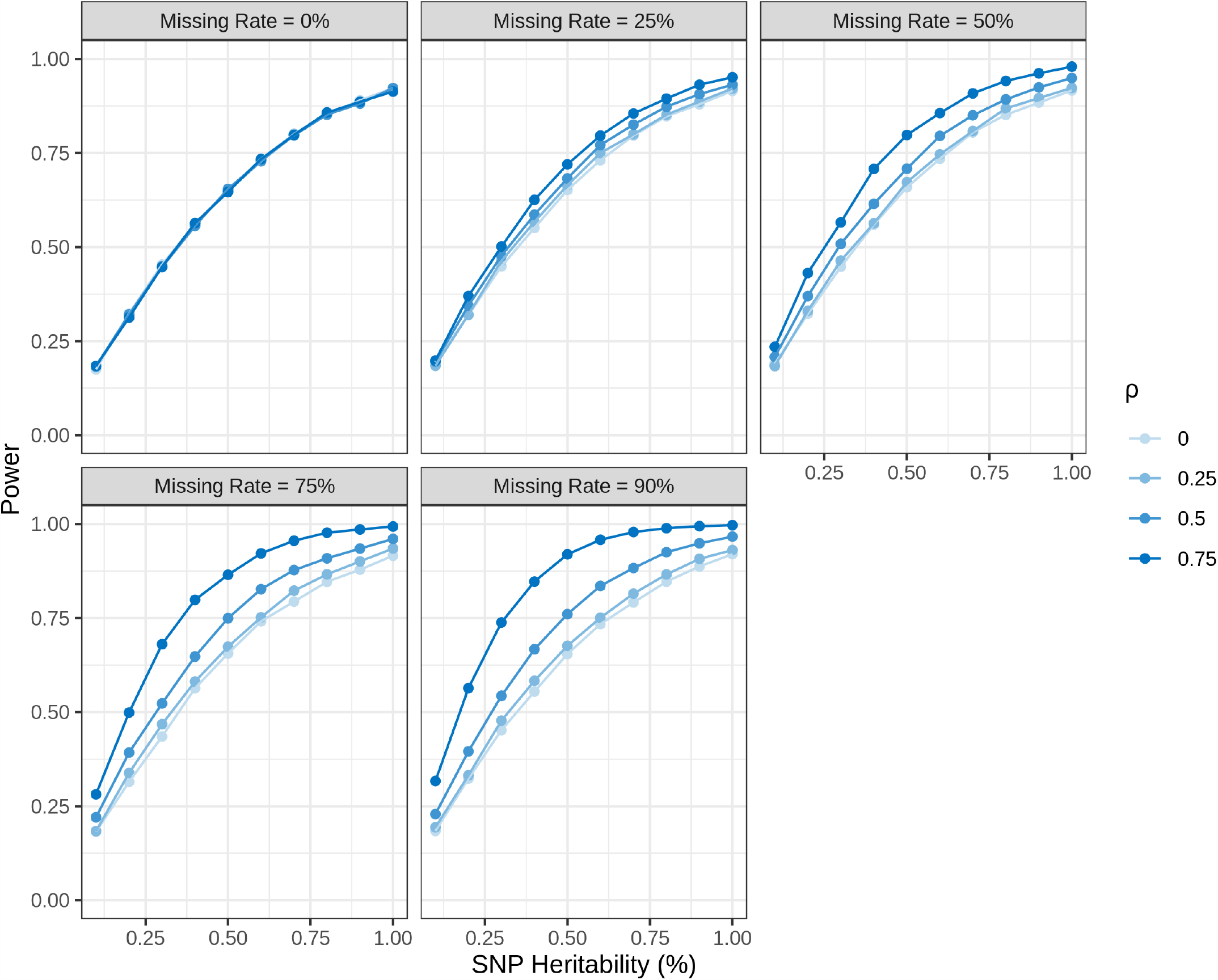
Power of SynSurr across various missing rates, target-surrogate correlations, and SNP heritabilities. In all cases, the number of subjects with observed phenotypes was *n* = 10^3^. The number of subjects with missing phenotypes was varied to achieve the indicated level of missingness. In each panel, the synthetic surrogate has correlation *ρ ∈ {*0.00, 0.25, 0.50, 0.75*}* with the target phenotype and the SNP heritability was varied from 0.1% to 1%. When there is no missingness, SynSurr is equivalent to the standard analysis and shows no variation across values of *ρ*. The power of SynSurr increases with increasing missingness and target-surrogate correlation. The number of simulation replicates is 10^4^.

### Evaluation on UK Biobank (UKB) data

We next demonstrate the advantages of SynSurr over a standard GWAS through multiple analyses in the UKBB. The UKBB is a research cohort composed of approximately 500K participants, aged 40 to 69, who agreed to provide medical histories, physical measurements, and biospecimens for research use between 2007 and 2010^36^. Upon enrollment, standard anthropometric measurements were captured, including height and weight, as were imprecise measurements of body composition ascertained by bioelectrical impedance^36,37^.

We first consider GWAS of two well-studied traits, height and FEV1, that were measured for a majority of the cohort. This analysis enables us to compare the performance of SynSurr with standard GWAS as the target phenotype is increasingly ablated. We then perform SynSurr analysis of 6 incompletely observed DEXA body composition phenotypes. In contrast to impedance, DEXA is a highly precise method of evaluating body composition^38,39^. Ascertainment of DEXA scans began in 2014 with an imaging pilot study among 5K randomly selected subjects, and imaging is expected for up to 100K subjects by the end of 2023^38^. At the time of our study, DEXA measurements of body composition were available for around 30K participants, whereas anthropometry and impedance measurements were available for nearly 350K participants. This provided a natural opportunity to deploy SynSurr for GWAS of DEXA traits, leveraging synthetic surrogates derived from baseline anthropometry and impedance.

### SynSurr recovers more oracle associations than standard or imputation-based GWAS as the target phenotype is increasingly ablated

The genotype and sample quality control steps are described in the Methods. Our analyses included 435,468 directly genotyped genetic variants. In total, 349,474 unrelated subjects were included in the height GWAS and 308,518 in the FEV1 GWAS. For each phenotype, sets of kin up to the third-degree were identified. One subject was randomly allocated to the GWAS data set and the remaining subjects to the model-building data set. Note that the subjects allocated to model-building could not otherwise participate in the GWAS as our bivariate association model (Equation 5) does not currently accommodate related individuals. The model-building data were used to construct a random forest model for predicting the target phenotype on the basis of age, sex, and various anthropomorphic measurements (see Supplementary Section 9.1 for details). Having related subjects in the model-building and inference (GWAS) data sets is not problematic because the GWAS data set is not being used to evaluate generalization performance.

For each phenotype, we performed an oracle GWAS prior to the introduction of missingness. This GWAS establishes the number of genome-wide significant (GWS) associations that would be detected if the target phenotype were fully observed. Then, 25%, 50%, 75%, and 90% of the target phenotypes were ablated at random. Both standard and SynSurr GWAS were performed on the remaining data. Prior to the introduction of missingness, the correlation between the target and synthetic surrogate phenotypes in the GWAS data set was *R*^2^ = 0.67 for height and *R*^2^ = 0.51 for FEV1. Scatter plots of predicted vs. observed height and FEV1 in the model-building and GWAS data are shown in **Supplementary Figures 7-8. Table 1** presents the numbers of oracle associations recovered by both the standard and SynSurr GWAS. SynSurr consistently recovers a higher proportion of the oracle associations than standard GWAS. Importantly, as demonstrated in **Supplementary Table 6**, SynSurr does not achieve this higher recovery by having a higher false discovery rate (FDR). Rather, SynSurr is leveraging the correlated surrogate outcome to obtain more precise SEs (**Supplementary Table 7**; also see **Supplementary Table 8**). For example, at 90% missingness, the mean square standard error of the SynSurr estimator, across oracle associations, is 7.4*×*10^−5^ (height) and 8.3*×*10^−5^ (FEV1) compared with 9.8*×*10^−5^ (height) and 9.7*×*10^−5^ (FEV1) for the standard estimator (relative improvements of 33% and 17%, respectively). **Supplementary Figure 9** verifies that SynSurr is estimating the same genetic effect as the oracle GWAS (so too is standard GWAS, see **Supplementary Figure 10**). Even with 90% of target phenotypes ablated, the *R*^2^ for the genetic effects between SynSurr and oracle is 0.90 for height and 0.87 for FEV1. With 50% of target phenotypes ablated, the *R*^2^ rises to 0.99 and 0.98 respectively, and in the absence of missingness, *R*^2^ = 1.00.

**Table 1:** Number of genome-wide significant SNPs recovered by standard and SynSurr GWAS across increasing ablation of the target phenotype. The oracle method establishes the number of genome-wide significant (GWS) variants (*p <* 5*×* 10^−8^) that would be identified in the absence of missingness. Missingness was introduced by ablating 25%, 50%, 75%, and 90% of the target phenotypes. Standard and SynSurr GWAS were performed on each of the ablated data sets, where standard GWAS refers to performing GWAS using only the observed values of the target outcome. The number and percentage of full-sample (oracle) GWS variants recovered by each method is reported. The false negative rate is 100% minus the recovery rate shown. Also see Supplementary Tables 6-8.

| Missing Rate (%) | Height |  |  |  | FEV1 |  |  |  |
| --- | --- | --- | --- | --- | --- | --- | --- | --- |
| | $n_{obs}$ | Oracle | Standard | SynSurr | $n_{obs}$ | Oracle | Standard | SynSurr |
| 0 | 349,474 | 7,177 | 7,177(100%) | 7,177(100%) | 308,518 | 974 | 974(100%) | 974(100%) |
| 25 | 262,105 | 7,177 | 4,896(68.22%) | 5,305(73.92%) | 231,388 | 974 | 546(56.06%) | 599(61.50%) |
| 50 | 174,737 | 7,177 | 2,742(38.21%) | 3,421(47.67%) | 154,259 | 974 | 278(28.54%) | 326(33.47%) |
| 75 | 87,368 | 7,177 | 834(11.62%) | 1,243(17.32%) | 77,129 | 974 | 60(6.16%) | 81(8.32%) |
| 90 | 34,947 | 7,177 | 192(2.68%) | 329(4.58%) | 30,852 | 974 | 0(0%) | 0(0%) |

Working within the ablation framework, we also compared SynSurr with imputation-based GWAS and Multi-Trait Analysis of GWAS (MTAG)^40^. Beginning with imputation, we focused on the setting of 50% missingness, comparing SynSurr with SI and MI when the surrogates and the imputations were either high quality, generated by random forest, or low quality, generated by linear regression. To showcase the fragility of imputation-based inference, we considered cases where the surrogates and imputations were either permuted or negated. The results are presented in **Supplementary Tables 9-13**. As expected, SI fails to consistently control the FDR, and although MI performed better in this regard, it was underpowered, identifying fewer than 20% of the associations identified by SynSurr. While permutation or negation compromised imputation-based inference, SynSurr was robust to permutation, performing comparably to standard GWAS, and is unaffected by negation. **Supplementary Figures 11-12** demonstrate that even when SI or MI properly control the FDR, the estimated genetic effects are generally biased, whereas those of SynSurr are always unbiased.

Next, we first performed proxy GWAS (i.e. standard GWAS of the synthetic surrogate) then combined standard GWAS of the target outcome with proxy GWAS via MTAG. The results are presented in **Supplementary Tables 16-17**. Due to imperfect prediction of the target outcome, proxy GWAS poorly controlled the FDR, identifying numerous significant associations not detected by the oracle. As a result, MTAG tended to inherit an inflated FDR. For example, in the case of FEV1, proxy GWAS had a FDR of 86%; MTAG of standard GWAS with proxy had a FDR rising from 19% in the absence of missingness to 67% at 90% missingness. In cases where MTAG did control the FDR, SynSurr typically had better power.

### SynSurr identifies more associations with partially missing body composition phenotypes than standard GWAS

We next performed SynSurr analyses of 6 incompletely measured DEXA phenotypes: android, arm, gynoid, leg, trunk, and total mass. In each case, the model-building data set included 4,584 subjects with observed target outcomes, while the GWAS data set included 347,498 subjects, 29,577 (8.5%) of which have observed target phenotypes. Within the model-building data set, a random forest was trained to predict the DEXA phenotype on the basis of age, sex, height, body weight, body mass index, and 5 measures of impedance: whole body, left/right arm, left/right leg. The fitted models were then transferred to the GWAS data set, where a synthetic surrogate outcome was generated for all subjects. Among subjects in the inference data set with an observed DEXA phenotype, the average *R*^2^ between the target phenotype and the synthetic surrogate was 0.80 (**Supplemental Figure 13**).

**Figure 5** presents the number of GWS associations (*p <* 5 *×* 10^−8^) for standard GWAS, based on the subjects with observed DEXA measurements only, and SynSurr GWAS, as well as the average *χ*^2^ statistic at the union of variants that reached significance in either GWAS. A greater expected *χ*^2^ statistic directly corresponds to greater power to detect an association. Standard GWAS identified between 8 and 10 GWS variants (8.3 on average), while SynSurr GWAS identified between 65 and 270 GWS variants, for an average of 179.5 (21.5-fold improvement; **Supplemental Table 18**). The average *χ*^2^ statistic at GWS variants was 46.2 for SynSurr GWAS, compared with 14.1 for standard GWAS, a 3.3-fold improvement. To check control of the type I error, the SynSurr analysis was repeated using permuted phenotypes. The uniform quantile-quantile plots in **Supplementary Figure 14** show no evidence of type I error inflation under the null.

**Figure 5:**
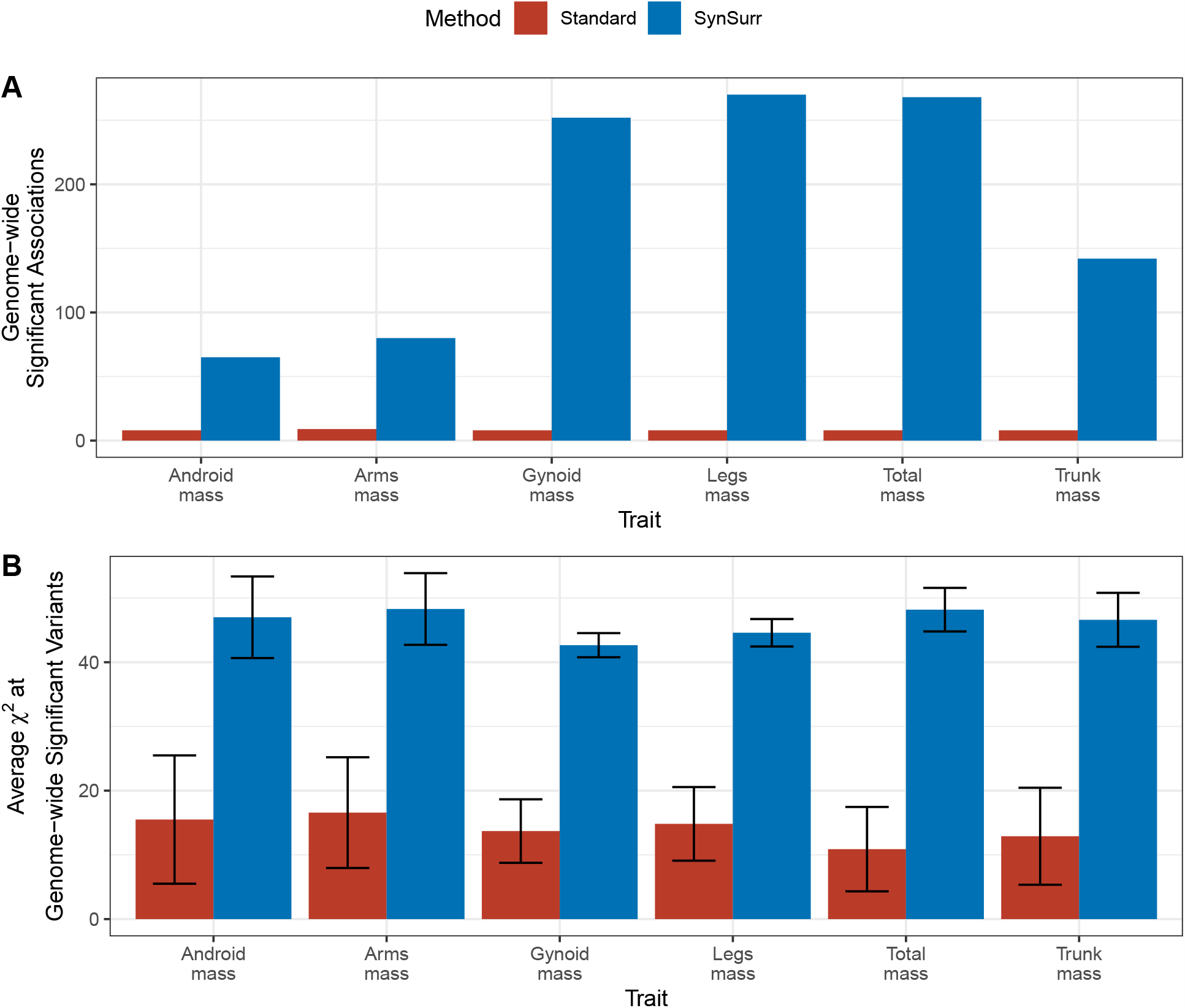
Comparing SynSurr and standard GWAS with respect to the number and significance of genome-wide significant associations for body composition traits. **A**. Number of genome-wide significant associations (*p <* 5*×* 10^−8^) with DEXA body composition traits for standard and synthetic surrogate (SynSurr) GWAS. **B**. Average *χ*^2^ statistic at the union of variants that reached genome-wide significance under either method. Error bars are 95% confidence intervals. A greater expected *χ*^2^ statistic directly corresponds to greater power to detect an association.

The Miami plots in **Supplementary Figure 15** show how SynSurr can elevate a subthreshold signal to genome-wide significance. For example, the association of rs2814993 with leg mass, which has a suggestive *P* = 1.1 *×* 10^−6^ with standard GWAS, becomes GWS with SynSurr *P* = 2.3 *×* 10^−20^ with SynSurr (in fact, this SNP is significant for all DEXA traits via SynSurr). rs2814993 was previously associated with height in a meta-analysis^41^ of European populations and an Australian twin study^42^. As another example, rs17782313 is associated with all DEXA traits by SynSurr, foremost with total mass *P* = 1.8 *×* 10^−26^, but at best reaches a *P* of 1.6 *×* 10^−5^ with standard GWAS. This SNP has a well-characterized association with obesity^43–45^.

Although we are aware of no independent GWAS of the same traits, for external validation SynSurr’s findings were overlapped with publicly available associations from the GWAS Catalog^32^ for the following traits: body fat distribution, body fat percentage, fat body mass, lean body mass (**Figure 6**). On average, 70% of the DEXA associations identified by SynSurr overlapped with a previously reported body composition association from the GWAS Catalog. As an internal validation, we performed a split-sample analysis, randomly allocating our GWAS data set 80:20 to independent discovery and validation cohorts. On average, 75.8% of GWS associations identified by SynSurr in the discovery cohort (*N* = 277, 998) replicated in the validation cohort (*N* = 69, 500)(**Supplemental Table 19**). Moreover, for all traits, the genetic correlation between the discovery and validation cohorts was high, 94.2% on average, with the 95% confidence interval always including 1.0 (**Supplemental Figure 17**), underscoring the reproducibility of SynSurr’s findings.

**Figure 6:**
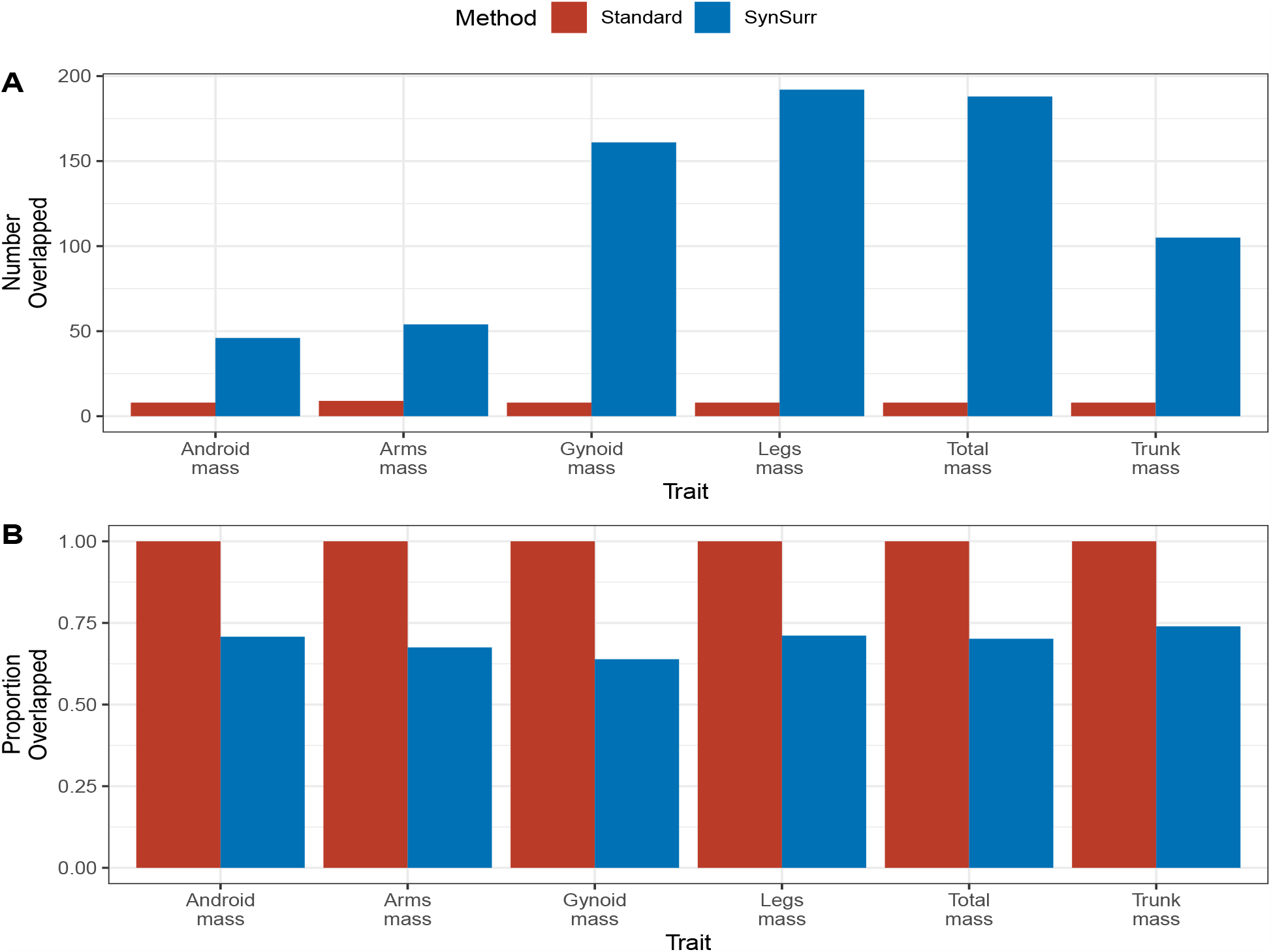
External validation via overlap of genome-wide significant variants for body composition with associations from the GWAS catalog. Variants from the GWAS catalog associated with body fat distribution, body fat percentage, fat body mass, and lean body mass were compiled. A study variant was considered overlapped with it fell within 250 kb of a GWAS catalog variant. Panels A and B show the counts and proportions of overlapped variants, respectively. Note that, with 1 exception, all variants identified by standard GWAS were also identified by SynSurr (**Supplemental Table 18**). The perfect overlap of the standard GWAS variants with known body composition associations in panel B is a direct consequence of the standard GWAS detecting very few genome-wide significant variants (8.3 on average), and indicates that none of these variants are novel.

To further investigate the biological function of the GWS variants, we performed gene set enrichment analysis using FUMA^46^. On average, 509 gene sets were enriched among the SynSurr results, while no significant enrichment were identified with the standard GWAS results. For all phenotypes, gene sets related to body fat distribution were among the most significant, and numerous enrichments related to anthropometric traits were identified (**Supplementary Data 1**).

## Discussion

We introduced SynSurr, a simple and computationally tractable procedure for leveraging a surrogate phenotype generated by a machine learning (ML)-model to improve inference on partially missing target phenotype. In the course of a typical SynSurr analysis, an ML-model is trained to generate a prediction of the target outcome (a “synthetic surrogate”) as a function of available information, then a GWAS is performed by jointly regressing the partially missing target phenotype and its synthetic surrogate on genotype and covariates. Unlike imputation, SynSurr is robust to the choice of surrogate, not requiring the often untenable assumption that the synthetic surrogate is generated from a correctly specified imputation model. We demonstrated that a SynSurr GWAS is robust, powerful, and valid: providing power at least equaling, but typically exceeding standard GWAS, while controlling the type I error and estimating the proper genetic effect. The power advantage afforded by SynSurr increases with the extent of target missingness and the target-surrogate correlation. In real data ablation analyses, SynSurr consistently recovered more of the full-sample (oracle) associations than standard GWAS or multiple-imputation, and did so without either inflating the false discovery rate or distorting the estimated genetic effects. Instead, SynSurr leverages correlation between the target and synthetic surrogate to obtain more precise estimates of the parameter of interest (i.e. to reduce the standard error). Finally, when applied to GWAS of 6 partially missing body composition phenotypes from the UKB, SynSurr identified 21.5 times more genome-wide significant associations than standard GWAS. These associations were significantly enriched for gene sets related to body composition and overlapped considerably with body composition associations from the GWAS catalog.

Although performing valid inference by means of imputation is possible, it typically requires both a correctly specified imputation model and the use of multiple imputations to correctly estimate standard errors. Here we propose jointly modeling a partially missing outcome together with its predicted value, rather than replacing missing values by their predicted values. In so doing, we avoid the often unverifiable assumption that the imputation model is correctly specified, and obtain valid statistical inference in a single shot. This robust approach to handling missing data has broad potential for applications outside of GWAS.

For all analyses reported in the main text, the model-building and GWAS data sets were independent. In our UKB analyses, the relatives of subjects in the GWAS data set formed a natural cohort for model-building. For the simple class of random forest models utilized for generating synthetic surrogates, this cohort was sufficiently sized to obtain reasonable target-surrogate correlations. For more complex models (e.g. deep neural networks), larger model-building sets are typically required. Analyses in simulated (**Supplementary Figure 4**) and real (**Supplementary Tables 14-15**) data suggest that utilizing the same subjects for both model-building and GWAS neither biases the estimated genetic effects nor inflates the type I error. While dropping the requirement for independent model-building and GWAS data sets would simplify analyses and facilitate the development of more complex surrogate models, more empirical and theoretical work is needed to understand the conditions under which this remains valid. Notably, utilizing the labeled subjects to train the model that the generates synthetic surrogates for all subjects would seem to introduce dependencies across subjects and invalidate the likelihood in **Supplemental Equations (6-7)**.

Several additional simulation studies are reported in the Supplementary Materials. Supplemental Section 6 examines the trade-off between allocating subjects to the model-building data set, to improve the quality of the synthetic surrogate, versus allocating subjects to the GWAS data set, to improve power, given that the model-building and GWAS data sets will be kept independent. We find that allocating more of the labeled subjects to GWAS, as opposed to model-building, tends to improve power (**Supplementary Figure 5**). Supplemental Section 7 examines the validity of imputing a predictor used as an input to the surrogate model. Unlike imputing the target outcome (i.e. *Y*), imputing an input to the surrogate model does not modify the relationship of interest (i.e. that between *G* and *Y*, as quantified by *β*_*G*_). Thus, as demonstrated in **Supplemental Figure 6**, imputing an input to *Ŷ* neither biases the estimated genetic effects nor inflates the type I error.

Lastly, SynSurr is not without limitations. First, the method was derived under joint normality of the target phenotype and the synthetic surrogate. As phenotypes and surrogates may follow non-normal distributions, we suggest applying the rank-based inverse normal transformation (INT) to both the target and the surrogate prior to analysis^47^, and did so for all analyses presented in this paper. While INT only guarantees marginal normality of each phenotype, it is commonly used in GWAS and mitigates the extent to which the joint distribution can depart from bivariate normality. Our current work focuses on extending SynSurr to a broader class of distributions by dropping the bivariate normality assumption. This can be achieved by replacing the score equations derived from maximum likelihood theory with a set of weighted estimating equations^48^. Moreover, we plan to extend SynSurr to generalized linear models with outcomes from the exponential family using a zero-mean augmentation approach^20,49^. Second, the validity of SynSurr assumes that the missing phenotypes are missing at random (MAR; see Supplementary Methods). While MAR is expected to hold for the DEXA phenotypes considered in this study, because the invitation to participate in imaging did not depend on a subject’s actual or expected body composition^38^, it may fail in settings where the factors affecting ascertainment are unknown^50^. It is of future interest to study the performance of SynSurr in settings where data are missing not at random. Third, SynSurr does not currently accommodate related individuals. Future work will extend SynSurr to allow for related individuals by adding a subject-specific random effect with cross-subject covariance proportional to the genetic relatedness matrix^51^. Finally, SynSurr currently requires individual-level data. Reworking the estimation procedure to enable SynSurr analysis from summary statistics would enable applications in settings where individual-level data are unavailable.

## Methods

### Standard GWAS

For the standard GWAS, the target phenotype *Y* is regressed on genotype *G* and covariates *X* among those subjects whose target phenotypes are observed:

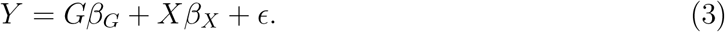

The null hypothesis *H*_0_ : *β*_*G*_ = 0 is evaluated using the standard two-sided Wald test^52^:

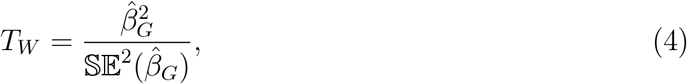

where 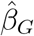 is the ordinary least squares (OLS) estimate of the genetic effect, and 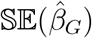 is the corresponding standard error.

### SynSurr GWAS

Within the inference data set, suppose the target phenotype *Y* is observed for *n*_obs_ subjects and missing for *n*_miss_. Let *n* = *n*_obs_ + *n*_miss_ denote the total sample size. Denote by *Ŷ* a synthetic surrogate for *Y* that is available for all *n* subjects. We recommend constructing *Ŷ* by means of a nonlinear ML-model, such as a random forest or neural network, trained on an independent model-building data set. Let *G* denote the genotype and *X* a vector of covariates, such as age, sex, and genetic principal components. To make use of both *Y* and *Ŷ* in evaluating the association between *Y* and *G*, SynSurr utilizes the joint association model:

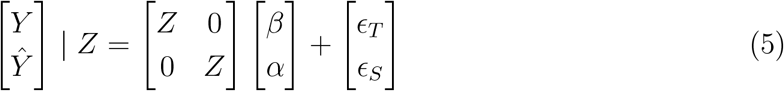

where *Z* = (*G, X*)^*T*^, *β* = (*β*_*G*_, *β*_*X*_)^*T*^, *α* = (*α*_*G*_, *α*_*X*_)^*T*^, and the residuals (*ϵ*_*T*_, *ϵ*_*S*_)^*T*^ follow a bivariate normal distribution:

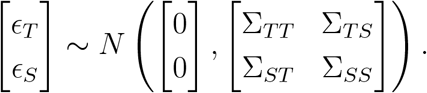

Here, the subscripts of *T* and *S* denote the *target* phenotype and the *synthetic surrogate* respectively. In model (5), *β*_*G*_ is the parameter of interest, which quantifies the association between the genotype and the target phenotype. This is the same parameter estimated by standard GWAS of *Y* on *G* among the *n*_obs_ subjects with observed target phenotypes in (3). Proxy GWAS, which regresses *Ŷ* rather than *Y* on *G*, estimates *α*_*G*_ instead of *β*_*G*_. Needless to say, *α*_*G*_ can differ from *β*_*G*_, and likely will when *Ŷ* is generated from a misspecified model. SynSurr enables inference on *β*_*G*_ while making use of all *n* subjects. Moreover, SynSurr is computationally tractable at biobank scale, requiring only two ordinary least squares regressions:

1. First, among all *n* subjects, regress *Ŷ* on *Z* to obtain an estimate of *α*.
2. Second, among the *n*_obs_ subjects with observed phenotypes, regress *Y* on (*Ŷ, Z*)^*T*^ to obtain an estimate of the associated regression coefficient, denoted (*δ, γ*)^*T*^ .

The validity of this two-step approach is demonstrated through a reparameterization of the log-likelihood function which allows the association parameter can be recovered as *β* = *γ*+*δα*. Details of this equivalence, as well as derivation of the Wald test for SynSurr, are provided in the Supplementary Methods.

### UKBB genotype and sample quality control

Our UKBB data release contains genotypes for 488,377 subjects and 784,256 directly genotyped variants. Prior to the analysis, we performed the following common quality control steps^53^:

1. Excluded individuals with *>* 10% missing genotypes.
2. Excluded SNPs with a genotyping rate *<* 90%.
3. Excluded SNPs with a Hardy-Weinberg Equilibrium *p <* 10^−5^.
4. Excluded SNPs with MAF *<* 1%.
5. Included only who self-identified as ‘White British’ and have very similar genetic ancestry based on a principal components analysis of the genotypes (Data-Field 22006).
6. Selected one member of each set of subjects with a kinship coefficient greater than 0.0625 (the threshold for third-degree relatives) for the GWAS data set, and allocated the remaining subjects to the model-building data set.

The results from our genotype QC are summarized in **Supplementary Table 4**. Note that we allowed the model-building data set to contain related individuals.

### GWAS catalog overlap analysis

Summary statistics for `body fat distribution, body fat percentage, fat body mass, lean body` mass were downloaded from the NHGRI-EBI GWAS catalog^32^. After concatenating and reducing to 1 record per unique combination of chromosome and base pair, this set contained 984 associated variants. Overlap of study variants with GWAS catalog variants was assessed using the `GenomicRanges`^54^ package in `R`^55^. A study variant was considered overlapped if fell within 250 kb of GWAS catalog variant for one of the aforementioned traits.

## Data availability

This work used genotypes and phenotypes from the UK Biobank study (https://www.ukbiobank.ac.uk) and our access was approved under application 64875.

## Code availability

SurrogateRegression is available as an R^55^ package on the Comprehensive R Archive Network: https://CRAN.R-project.org/package=SurrogateRegression.

Replication code for the analyses presented in this paper is available on GitHub at: https://github.com/jianhuig/SyntheticSurrogateAnalysis.

## Acknowledgments

This work was supported by the National Institutes of Health grants R35 CA197449 and F31 HL140822 (to ZRM); and R35-CA197449, U19-CA203654, R01-HL163560, U01-HG012064, and U01-HG009088 (to XL); and the Natural Sciences and Engineering Research Council of Canada grant RGPIN-2021-03734 and a Connaught New Researcher Award (to JeG).

## Author information

These authors jointly supervised this work: Xihong Lin and Jessica Gronsbell.

## Contributions

ZRM, XL, and JeG designed the study and the experiments. ZRM implemented the software with input from XL. ZRM and JiG performed the simulation experiments. JiG conducted analyses of the UK Biobank data. ZRM performed the overlap analysis. ZRM and JeG wrote the first draft of the manuscript and all co-authors provided intellectual revisions.

## References

1. Kurki, M., Karjalainen, J., Palta, P., et al. FinnGen provides genetic insights from a well-phenotyped isolated population. Nature 613, 508–518 (2023).

2. Gaziano, J. M. et al. Million Veteran Program: A mega-biobank to study genetic influences on health and disease. Journal of clinical epidemiology 70, 214–223 (2016).

3. Bycroft, C. et al. The UK Biobank resource with deep phenotyping and genomic data. Nature 562, 203–209 (2018).

4. Beesley, L. J. et al. The emerging landscape of health research based on biobanks linked to electronic health records: Existing resources, statistical challenges, and potential opportunities. Statistics in medicine 39, 773–800 (2020).

5. Tan, V. Y. & Timpson, N. J. The UK Biobank: A Shining Example of Genome-Wide Association Study Science with the Power to Detect the Murky Complications of Real-World Epidemiology. Annual Review of Genomics and Human Genetics 23 (2022).

6. Wei, W.-Q. & Denny, J. C. Extracting research-quality phenotypes from electronic health records to support precision medicine. Genome medicine 7, 1–14 (2015).

7. Banda, J. M., Seneviratne, M., Hernandez-Boussard, T. & Shah, N. H. Advances in electronic phenotyping: from rule-based definitions to machine learning models. Annual review of biomedical data science 1, 53 (2018).

8. Allen, N. E., Sudlow, C., Peakman, T., Collins, R., et al. UK biobank data: come and get it. 2014.

9. Littlejohns, T. J. et al. The UK Biobank imaging enhancement of 100,000 participants: rationale, data collection, management and future directions. Nature communications 11, 1–12 (2020).

10. Elliott, L. et al. Genome-wide association studies of brain imaging phenotypes in UK Biobank. Nature 562, 210–216 (2018).

11. Pirruccello, J. et al. Analysis of cardiac magnetic resonance imaging in 36,000 individuals yields genetic insights into dilated cardiomyopathy. Nat Commun 11, 2254 (2020).

12. Alipanahi, B. et al. Large-scale machine learning-based phenotyping significantly improves genomic discovery for optic nerve head morphology 2020. arXiv: 2011.13012 [q-bio.GN].

13. Li, X. & Zhao, H. Automated feature extraction from population wearable device data identified novel loci associated with sleep and circadian rhythms. PLoS Genet 16, e1009089 (2020).

14. Zhang, Y. et al. High-throughput phenotyping with electronic medical record data using a common semi-supervised approach (PheCAP). Nature protocols 14, 3426–3444 (2019).

15. Liao, K. P. et al. High-throughput multimodal automated phenotyping (MAP) with application to PheWAS. Journal of the American Medical Informatics Association 26, 1255–1262 (2019).

16. Yang, S., Varghese, P., Stephenson, E., Tu, K. & Gronsbell, J. Machine learning approaches for electronic health records phenotyping: A methodical review. medRxiv (2022).

17. Hormozdiari, F. et al. Imputing phenotypes for genome-wide association studies. The American Journal of Human Genetics 99, 89–103 (2016).

18. Wang, S., McCormick, T. H. & Leek, J. T. Methods for correcting inference based on outcomes predicted by machine learning. Proceedings of the National Academy of Sciences 117, 30266–30275 (2020).

19. Huang, J. et al. PIE: A prior knowledge guided integrated likelihood estimation method for bias reduction in association studies using electronic health records data. Journal of the American Medical Informatics Association 25, 345–352 (2018).

20. Tong, J. et al. An augmented estimation procedure for EHR-based association studies accounting for differential misclassification. Journal of the American Medical Informatics Association 27, 244–253 (2020).

21. Rubin, D. B. Inference and missing data. Biometrika 63, 581–592 (1976).

22. Hubbard, R. A., Tong, J., Duan, R. & Chen, Y. Reducing bias due to outcome misclassification for epidemiologic studies using EHR-derived probabilistic phenotypes. Epidemiology 31, 542–550 (2020).

23. Hong, C., Liao, K. P. & Cai, T. Semi-supervised validation of multiple surrogate outcomes with application to electronic medical records phenotyping. Biometrics 75, 78–89 (2019).

24. Little, R. J. & Rubin, D. B. Statistical Analysis with Missing Data 2nd (John Wiley & Sons, 2002).

25. Rubin, D. Multiple Imputation for Nonresponse in Surveys (John Wiley & Sons, 1987).

26. Rubin, D. B. Multiple imputation after 18+ years. Journal of the American statistical Association 91, 473–489 (1996).

27. Van Buuren, S. Flexible Imputation of Missing Data 2nd (Chapman and Hall/CRC, 2018).

28. Bartlett, J. W. & Hughes, R. A. Bootstrap inference for multiple imputation under uncongeniality and misspecification. Statistical methods in medical research 29, 3533–3546 (2020).

29. Austin, P. C., White, I. R., Lee, D. S. & van Buuren, S. Missing data in clinical research: a tutorial on multiple imputation. Canadian Journal of Cardiology 37, 1322–1331 (2021).

30. Murray, J. S. Multiple imputation: a review of practical and theoretical findings. Statistical Science 33, 142–159 (2018).

31. McCaw, Z. R., Gaynor, S. M., Sun, R. & Lin, X. Leveraging a surrogate outcome to improve inference on a partially missing target outcome. Biometrics Online ahead of print (2022).

32. Buniello, A., MacArthur, J., Cerezo, M., et al. The NHGRI-EBI GWAS Catalog of published genome-wide association studies, targeted arrays and summary statistics 2019. Nucleic Acids Res 47, D1005–D1012 (2019).

33. Breiman, L. Random forests. Machine learning 45, 5–32 (2001).

34. Chen, T. & Guestrin, C. XGBoost: A Scalable Tree Boosting System. CoRR abs/1603.02754. arXiv: 1603.02754. http://arxiv.org/abs/1603.02754 (2016).

35. Casella, B. & Berger, R. Statistical Inference. 2nd ed. (Duxbury/Thomson Learning, Pacific Grove, CA, 2002).

36. Allen, N. E., Sudlow, C., Peakman, T., Collins, R. & biobank, U. UK biobank data: come and get it 2014.

37. Biobank, U. UK Biobank Body Composition Measurement https://biobank.ndph.ox.ac.uk/showcase/refer.cgi?id=1421. 2011.

38. Littlejohns, T., Holliday, J., Gibson, L., et al. The UK Biobank imaging enhancement of 100,000 participants: rationale, data collection, management and future directions. Nat Commun 11, 2624 (2020).

39. Biobank, U. UK Biobank Imaging Modality DXA https://biobank.ndph.ox.ac.uk/showcase/refer.cgi?id=502. 2015.

40. Turley, P., Walters, R., Maghzian, O., et al. Multi-trait analysis of genome-wide association summary statistics using MTAG. Nature Genetics 50, 229–237 (2018).

41. Weedon, M., Lango, H., Lindgren, C., et al. Genome-wide association analysis identifies 20 loci that influence adult height. Nature Genetics 40, 575–583 (2008).

42. Liu, J., Medland, S., Wright, M., et al. Genome-wide association study of height and body mass index in Australian twin families. Twin Res Hum Genet 13, 179–193 (2010).

43. Meyre, D., Delplanque, J., Chevre, J., et al. Genome-wide association study for earlyonset and morbid adult obesity identifies three new risk loci in European populations. Nature Genetics 41, 157–159 (2009).

44. Willer, C., Speliotes, E., Loos, R., et al. Six new loci associated with body mass index highlight a neuronal influence on body weight regulation. Nature Genetics 41, 25–34 (2009).

45. Loos, R. & Yeo, G. The genetics of obesity: from discovery to biology. Nat Rev Genet 23, 120–133 (2022).

46. Watanabe, K., Taskesen, E., van Bochoven, A. & Posthuma, D. Functional mapping and annotation of genetic associations with FUMA. Nat Commun 8, 1826 (2017).

47. McCaw, Z., Lane, J., Saxena, R., Redline, S. & Lin, X. Operating characteristics of the rank-based inverse normal transformation for quantitative trait analysis in genome-wide association studies. Biometrics 76, 1262–1272 (2020).

48. Robins, J. & Rotnitzky, A. Semiparametric Efficiency in Multivariate Regression Models with Missing Data. Journal of the American Statistical Association 90, 122–129 (1995).

49. Wang, X. & Wang, Q. Semiparametric linear transformation model with differential measurement error and validation sampling. Journal of Multivariate Analysis 141, 67–80 (2015).

50. Little, R. J. & Rubin, D. B. Statistical analysis with missing data (John Wiley & Sons, 2019).

51. Po-Ru, L., Tucker, G., Bulik-Sullivan, B., Vilhjalmsson, B., Finucane, H., et al. Efficient Bayesian mixed model analysis increases association power in large cohorts. Nature Genetics 47, 284–290 (2015).

52. Seber, G. The Linear Model and Hypothesis. A General Unifying Theory 1st ed. (Springer Cham, 2015).

53. Purcell, S. et al. PLINK: a tool set for whole-genome association and population-based linkage analyses. The American Journal of Human Genetics 81, 559–575 (2007).

54. Lawrence, M., Huber, W., Pages, H., et al. Software for Computing and Annotating Genomic Ranges. PLoS Comput Biol 9, e1003118 (2013).

55. R Core Team. R: A Language and Environment for Statistical Computing R Foundation for Statistical Computing (Vienna, Austria, 2022). https://www.R-project.org/.

