## Supplemental for "Leveraging a machine learning derived surrogate phenotype to improve power for genome-wide association studies of partially missing phenotypes in population biobanks"

### 1 Estimation and Inference Procedures for the SynSurr Model

Recall that the SynSurr joint association model takes the form:

$$\begin{bmatrix} Y \\ \hat{Y} \end{bmatrix} \mid Z = \begin{bmatrix} Z & 0 \\ 0 & Z \end{bmatrix} \begin{bmatrix} \beta \\ \alpha \end{bmatrix} + \begin{bmatrix} \epsilon_T \\ \epsilon_S \end{bmatrix} \tag{1}$$

where $Z = (G, X)^T$, $\beta = (\beta_G, \beta_X)^T$, $\alpha = (\alpha_G, \alpha_X)^T$, and the residuals $(\epsilon_T, \epsilon_S)^T$ follow a bivariate normal distribution

$$\begin{bmatrix} \epsilon_T \\ \epsilon_S \end{bmatrix} \sim N\left( \begin{bmatrix} 0 \\ 0 \end{bmatrix}, \begin{bmatrix} \Sigma_{TT} & \Sigma_{TS} \\ \Sigma_{ST} & \Sigma_{SS} \end{bmatrix} \right).$$

Here $T$ stands for target phenotype, i.e. $Y$, and $S$ stands for surrogate phenotype, i.e. $\hat{Y}$.

In the presence of bilateral outcome missingness, model (1) may be estimated using an expectation conditional maximization algorithm[1,2]. Here we show that when missingness is confined to the target outcome, the maximum likelihood estimates (MLEs) of all parameters from model (1) may be obtained via two sequential ordinary least squares (OLS) regressions. Estimation and inference procedures for both bilaterial and unilaterial missingness are implemented in the `SurrogateRegression`[3] `R`[4] package.

#### 1.1 Reparameterization

From (1), the marginal distribution of the synthetic surrogate is:

$$\hat{Y} \sim N(Z\alpha, \Sigma_{SS}). \tag{2}$$

Given the synthetic surrogate $\hat{Y}$, the conditional distribution of the true phenotype is:

$$Y|(Z, \hat{Y}) \sim N\big\{Z\beta + \Sigma_{TS}\Sigma_{SS}^{-1}(\hat{Y} - Z\alpha), \Sigma_{TT} - \Sigma_{TS}\Sigma_{SS}^{-1}\Sigma_{ST}\big\}. \tag{3}$$

Consider the following reparameterization:

$$\delta = \Sigma_{TS}\Sigma_{SS}^{-1}, \qquad \gamma = \beta - \delta\alpha, \qquad \Lambda_{TT}^{-1} = \Sigma_{TT} - \Sigma_{TS}\Sigma_{SS}^{-1}\Sigma_{ST}. \tag{4}$$

Note that $\Lambda_{TT}$ is the upper left entry of the precision matrix $\Lambda = \Sigma^{-1}$:

$$\begin{bmatrix} \Lambda_{TT} & \Lambda_{TS} \\ \Lambda_{ST} & \Lambda_{SS} \end{bmatrix} = \begin{bmatrix} (\Sigma_{TT} - \Sigma_{TS}\Sigma_{SS}^{-1}\Sigma_{ST})^{-1} & -(\Sigma_{TT} - \Sigma_{TS}\Sigma_{SS}^{-1}\Sigma_{ST})^{-1}\Sigma_{TS}\Sigma_{SS}^{-1} \\ -(\Sigma_{SS} - \Sigma_{ST}\Sigma_{TT}^{-1}\Sigma_{TS})^{-1}\Sigma_{ST}\Sigma_{TT}^{-1} & (\Sigma_{SS} - \Sigma_{ST}\Sigma_{TT}^{-1}\Sigma_{TS})^{-1} \end{bmatrix}. \tag{5}$$

The inverse transformation to (4) is:

$$\Sigma_{TS} = \delta\Sigma_{SS}, \qquad \beta = \gamma + \delta\alpha, \qquad \Sigma_{TT} = \Lambda_{TT}^{-1} + \Sigma_{TS}\Sigma_{SS}^{-1}\Sigma_{ST}.$$

After reparameterizing, the conditional distribution of $Y$ given $\hat{Y}$ is:

$$Y|(Z,\hat{Y}) \sim N\big(Z\gamma + \delta\hat{Y},\ \Lambda_{TT}^{-1}\big).$$

#### 1.2 Likelihood

Collect the model parameters into $\theta = (\alpha, \gamma, \delta, \Sigma_{SS}, \Lambda_{TT}^{-1})$. Recall that:

$$R = \begin{cases} 1 & \text{if } Y \text{ is observed,} \\ 0 & \text{if } Y \text{ is missing.} \end{cases}$$

The likelihood contribution of a subject whose target outcome is missing ($R = 0$) is:

$$f(\hat{Y}|Z;\boldsymbol{\theta}) = f(\hat{Y}|Z;\alpha,\Sigma_{SS}),$$

while that of a subject with an observed target outcome ($R = 1$) is:

$$f(Y,\hat{Y}|Z;\theta) = f(Y|\hat{Y},Z;\gamma,\delta,\Lambda_{TT}^{-1}) \cdot f(\hat{Y}|Z;\alpha,\Sigma_{SS})$$

Indexing subjects by $i$, the observed data likelihood is:

$$L(\theta) = \prod_{\{i:R_i=1\}} f(Y_i|\hat{Y}_i,Z_i;\gamma,\delta,\Lambda_{TT}^{-1}) \cdot f(\hat{Y}_i|Z_i;\alpha,\Sigma_{SS}) \cdot \prod_{\{i:R_i=0\}} f(\hat{Y}_i|Z_i;\alpha,\Sigma_{SS}) \tag{6}$$

$$= \prod_{i=1}^{n} f(\hat{Y}_i|Z_i;\alpha,\Sigma_{SS}) \cdot \prod_{\{i:R_i=1\}} f(Y_i|\hat{Y}_i,Z_i;\gamma,\delta,\Lambda_{TT}^{-1}). \tag{7}$$

Because the likelihood factors as a product over variationally independent parameter spaces, the component likelihoods may be optimized separately to obtain the full MLE of $\theta$[5].

##### 1.2.1 Maximum Likelihood Estimates

The logarithm of the observed data likelihood decomposes as the sum of two terms:

$$\ell(\boldsymbol{\theta}) = \ell_S(\alpha,\Sigma_{SS}) + \ell_{T|S}(\gamma,\delta,\Lambda_{TT}^{-1}).$$

The first component:

$$\ell_S(\alpha,\Sigma_{SS}) = \sum_{i=1}^{n} \ln f(\hat{Y}_i|Z_i;\alpha,\Sigma_{SS})$$

is the marginal log likelihood of the synthetic surrogate, which receives a contribution from all $n$ subjects. The second component:

$$\ell_{T|S}(\gamma, \delta, \Lambda_{TT}^{-1}) = \sum_{\{i:R_i=1\}} \ln f(Y_i|\hat{Y}_i, Z_i; \gamma, \delta, \Lambda_{TT}^{-1})$$

is the conditional log likelihood of the true outcome, given the synthetic surrogate, and receives a contribution from subjects whose target outcomes are observed ($R_i = 1$).

In matrix notation, the marginal log likelihood is expressible as:

$$\ell_S(\alpha, \Sigma_{SS}) \propto -\frac{n}{2} \ln \Sigma_{SS} - \frac{\Sigma_{SS}^{-1}}{2} (\hat{\boldsymbol{y}} - \boldsymbol{Z}\alpha)^T (\hat{\boldsymbol{y}} - \boldsymbol{Z}\alpha), \tag{8}$$

where $\hat{\boldsymbol{y}} = (\hat{Y}_1, \cdots, \hat{Y}_n)^T$ is the $n \times 1$ vector of synthetic surrogates, and $\boldsymbol{Z}$ is the $n \times p$ matrix of subjects by genotypes and covariates. By maximizing (8), the MLEs of $\alpha$ and $\Sigma_{SS}$ are:

$$\hat{\alpha} = (\boldsymbol{Z}^T\boldsymbol{Z})^{-1}\boldsymbol{Z}^T\hat{\boldsymbol{y}}, \qquad \hat{\Sigma}_{SS} = \frac{1}{n}(\hat{\boldsymbol{y}} - \boldsymbol{Z}\hat{\alpha})^T(\hat{\boldsymbol{y}} - \boldsymbol{Z}\hat{\alpha}).$$

Observe that $\hat{\alpha}$ is the OLS estimate for regression of $\hat{\boldsymbol{y}}$ on $\boldsymbol{Z}$.

Suppose there are $n_{\text{obs}}$ subjects whose true outcomes are observed:

$$n_{\text{obs}} = \sum_{i=1}^{n} R_i. \tag{9}$$

The conditional log likelihood of $Y$ given $\hat{Y}$ is expressible as:

$$\ell_{T|S}(\gamma, \delta, \Lambda_{TT}^{-1}) = \frac{n_{\text{obs}}}{2} \ln \Lambda_{TT} - \frac{\Lambda_{TT}}{2} (\boldsymbol{y}_{\text{obs}} - \hat{\boldsymbol{y}}_{\text{obs}}\delta - \boldsymbol{Z}_{\text{obs}}\gamma)^T (\boldsymbol{y}_{\text{obs}} - \hat{\boldsymbol{y}}_{\text{obs}}\delta - \boldsymbol{Z}_{\text{obs}}\gamma),$$

where the subscript obs restricts a matrix or vector to the subset of $n_{\text{obs}}$ subjects whose true outcomes are observed. To further simplify, define the augmented design matrix $\boldsymbol{W}_{\text{obs}} = (\hat{\boldsymbol{y}}_{\text{obs}}, \boldsymbol{Z}_{\text{obs}})$, whose first column is $\hat{\boldsymbol{y}}_{\text{obs}}$ and whose remaining columns are $\boldsymbol{Z}_{\text{obs}}$, and the corresponding regression coefficient $\zeta = (\delta, \gamma)^T$. Then:

$$\ell_{T|S}(\zeta, \Lambda_{TT}^{-1}) \propto \frac{n_{\text{obs}}}{2} \ln \Lambda_{TT} - \frac{\Lambda_{TT}}{2} (\boldsymbol{y}_{\text{obs}} - \boldsymbol{W}_{\text{obs}}\zeta)^T (\boldsymbol{y}_{\text{obs}} - \boldsymbol{W}_{\text{obs}}\zeta). \tag{10}$$

(10) now has the same form as (8). Thus, the MLEs of $\zeta$ and $\Lambda_{TT}$ are:

$$\hat{\zeta} = (\boldsymbol{W}_{\text{obs}}^T\boldsymbol{W}_{\text{obs}})^{-1}\boldsymbol{W}_{\text{obs}}^T\boldsymbol{y}_{\text{obs}}, \qquad \Lambda_{TT}^{-1} = \frac{1}{n_{\text{obs}}}(\boldsymbol{y}_{\text{obs}} - \boldsymbol{W}_{\text{obs}}\hat{\zeta})^T(\boldsymbol{y}_{\text{obs}} - \boldsymbol{W}_{\text{obs}}\hat{\zeta}),$$

which are readily obtained via OLS of $\boldsymbol{y}_{\text{obs}}$ on $\boldsymbol{W}_{\text{obs}}$. Finally, by the invariance principle[6], the MLEs of all parameters from the original model (1) are:

$$\hat{\Sigma}_{TS} \leftarrow \hat{\delta}\hat{\Sigma}_{SS}, \qquad \hat{\beta} \leftarrow \hat{\gamma} + \hat{\delta}\hat{\alpha}, \qquad \hat{\Sigma}_{TT} \leftarrow \hat{\Lambda}_{TT}^{-1} + \hat{\Sigma}_{TS}\hat{\Sigma}_{SS}^{-1}\hat{\Sigma}_{ST}.$$

##### 1.2.2 Missingness

Unbiased estimation within the maximum likelihood framework requires that missingness occur at random[7]. Formally, the data are missing at random (MAR) if the observation indicator $R$ is independent of the target outcome $Y$ given the model covariates $Z$. In practice, MAR holds if the decision of whether to measure $Y$ for a particular subject does not depend on the value of $Y$, although it may depend on the value of $Z$. For example, MAR would hold if the subjects for whom $Y$ is measured were selected completely at random, meaning independently of any of the subject's characteristics. MAR also holds in cases where ascertain is differential across levels of a covariate, provided that covariate is included in $Z$. For example, suppose $Y$ is an imaging phenotype and subjects with a disease are imaged with much higher probability than subjects without the disease. This case falls within MAR provided disease status is included in $Z$. MAR would not hold if individuals with extreme values of $Y$ were systematically excluded from measurement.

#### 1.3 Inference under the SynSurr Model

Here we derive the Wald test of the null hypothesis $H_0 : \beta = 0$, from which tests regarding specific components of $\beta$ may easily be obtained.

##### 1.3.1 Score Equations

The score equation for $\beta$ is:

$$\mathcal{U}_\beta = \frac{\partial \ell}{\partial \beta} = \frac{\partial \gamma^T}{\partial \beta} \cdot \frac{\partial \ell_{T|S}}{\partial \beta} = \boldsymbol{I} \cdot \boldsymbol{Z}_{\text{obs}}^T \Lambda_{TT} (\boldsymbol{y}_{\text{obs}} - \hat{\boldsymbol{y}}_{\text{obs}} \delta - \boldsymbol{Z}_{\text{obs}} \gamma),$$

and that for $\alpha$ is:

$$\mathcal{U}_\alpha = \frac{\partial \ell}{\partial \alpha} = \frac{\partial \ell_S}{\partial \alpha} + \frac{\partial \gamma^T}{\partial \alpha} \cdot \frac{\partial \ell_{T|S}}{\partial \gamma} = \boldsymbol{Z}^T \Sigma_{SS}^{-1} (\hat{\boldsymbol{y}} - \boldsymbol{Z}\alpha) - \delta \boldsymbol{I} \cdot \boldsymbol{Z}_{\text{obs}}^T \Lambda_{TT} (\boldsymbol{y}_{\text{obs}} - \hat{\boldsymbol{y}}_{\text{obs}} \delta - \boldsymbol{Z}_{\text{obs}} \gamma)$$

Note that $\mathcal{U}_\beta$ only incorporates data from subjects with observed target outcomes, whereas $\mathcal{U}_\alpha$ incorporates data from all subjects.

##### 1.3.2 Fisher Information

The Fisher information for $\beta$ is:

$$\mathcal{I}_{\beta\beta^T} = -\mathbb{E}\left\{ \frac{\partial^2 \ell}{\partial \beta \partial \beta^T} \right\} = -\mathbb{E}\left\{ \frac{\partial \gamma^T}{\partial \beta} \frac{\partial^2 \ell_{T|S}}{\partial \gamma \partial \gamma^T} \frac{\partial \gamma}{\partial \beta^T} \right\} = \boldsymbol{Z}_{\text{obs}}^T \Lambda_{TT} \boldsymbol{Z}_{\text{obs}},$$

and that for $\alpha$ is:

$$\mathcal{I}_{\alpha\alpha^T} = -\mathbb{E}\left\{\frac{\partial^2 \ell}{\partial\alpha\partial\alpha^T}\right\} = -\mathbb{E}\left\{\frac{\partial^2 \ell_S}{\partial\alpha\partial\alpha^T} + \frac{\partial\gamma^T}{\partial\alpha}\frac{\partial^2 \ell_{T|S}}{\partial\gamma\partial\gamma^T}\frac{\partial\gamma}{\partial\alpha^T}\right\} = \boldsymbol{Z}^T\Sigma_{SS}^{-1}\boldsymbol{Z} + \delta^T\boldsymbol{Z}_{\text{obs}}^T\Lambda_{TT}\boldsymbol{Z}_{\text{obs}}\delta.$$

To further simplify, partition $\boldsymbol{Z} = \binom{\boldsymbol{Z}_{\text{obs}}}{\boldsymbol{Z}_{\text{miss}}}$, where $\boldsymbol{Z}_{\text{miss}}$ is the covariate submatrix for subjects whose target outcome is missing. Now:

$$\begin{aligned}\mathcal{I}_{\alpha\alpha^T} &= \begin{pmatrix}\boldsymbol{Z}_{\text{obs}}\\ \boldsymbol{Z}_{\text{miss}}\end{pmatrix}^T\Sigma_{SS}^{-1}\begin{pmatrix}\boldsymbol{Z}_{\text{obs}}\\ \boldsymbol{Z}_{\text{miss}}\end{pmatrix} + \delta^T\boldsymbol{Z}_{\text{obs}}^T\Lambda_{TT}\boldsymbol{Z}_{\text{obs}}\delta\\ &= \boldsymbol{Z}_{\text{obs}}^T\Sigma_{SS}^{-1}\boldsymbol{Z}_{\text{obs}} + \boldsymbol{Z}_{\text{miss}}^T\Sigma_{SS}^{-1}\boldsymbol{Z}_{\text{miss}} + \delta^T\boldsymbol{Z}_{\text{obs}}^T\Lambda_{TT}\boldsymbol{Z}_{\text{obs}}\delta\\ &= \boldsymbol{Z}_{\text{obs}}^T\big(\Sigma_{SS}^{-1} + \delta^T\Lambda_{TT}\delta\big)\boldsymbol{Z}_{\text{obs}} + \boldsymbol{Z}_{\text{miss}}^T\Sigma_{SS}^{-1}\boldsymbol{Z}_{\text{miss}}.\end{aligned}$$

Recall that $\delta = \Sigma_{TS}\Sigma_{SS}^{-1}$. Using the identity:

$$\Sigma_{SS}^{-1} + \delta^T\Lambda_{TT}\delta = \Sigma_{SS}^{-1} + \Sigma_{SS}^{-1}\Sigma_{ST}\Lambda_{TT}\Sigma_{TS}\Sigma_{SS}^{-1} = \Lambda_{SS},$$

obtained by block matrix inversion of $\boldsymbol{\Sigma}$ (see 5), the final form of the information for $\alpha$ is:

$$\mathcal{I}_{\alpha\alpha^T} = \boldsymbol{Z}_{\text{obs}}\Lambda_{SS}\boldsymbol{Z}_{\text{obs}} + \boldsymbol{Z}_{\text{miss}}^T\Sigma_{SS}^{-1}\boldsymbol{Z}_{\text{miss}}.$$

The cross information between $\alpha$ and $\beta$ is:

$$\mathcal{I}_{\beta\alpha^T} = -\mathbb{E}\left\{\frac{\partial^2 \ell}{\partial\beta\partial\alpha^T}\right\} = -\mathbb{E}\left\{\frac{\partial\gamma^T}{\partial\beta}\frac{\partial^2 \ell_{T|S}}{\partial\gamma\partial\gamma^T}\frac{\partial\gamma}{\partial\alpha^T}\right\} = -\boldsymbol{Z}_{\text{obs}}^T\Lambda_{TT}\boldsymbol{Z}_{\text{obs}}\delta.$$

Using the identity:

$$-\Lambda_{TT}\delta = -\Lambda_{TT}\Sigma_{TS}\Sigma_{SS}^{-1} = \Lambda_{TS},$$

the final form of the cross information is:

$$\mathcal{I}_{\beta\alpha^T} = \boldsymbol{Z}_{\text{obs}}\Lambda_{TS}\boldsymbol{Z}_{\text{obs}}.$$

Overall, the joint information for $(\beta, \alpha)$ is:

$$\mathcal{I} = \begin{pmatrix}\mathcal{I}_{\beta\beta^T} & \mathcal{I}_{\beta\alpha^T}\\ \mathcal{I}_{\alpha\beta^T} & \mathcal{I}_{\alpha\alpha^T}\end{pmatrix} = \begin{pmatrix}\boldsymbol{Z}_{\text{obs}}^T\Lambda_{TT}\boldsymbol{Z}_{\text{obs}} & \boldsymbol{Z}_{\text{obs}}^T\Lambda_{TS}\boldsymbol{Z}_{\text{obs}}\\ \boldsymbol{Z}_{\text{obs}}^T\Lambda_{ST}\boldsymbol{Z}_{\text{obs}} & \boldsymbol{Z}_{\text{obs}}^T\Lambda_{SS}\boldsymbol{Z}_{\text{obs}} + \boldsymbol{Z}_{\text{miss}}^T\Sigma_{SS}^{-1}\boldsymbol{Z}_{\text{miss}}\end{pmatrix}, \tag{11}$$

and the efficient information for $\beta$ is:

$$\mathcal{I}_{\beta\beta^T|\alpha} = \mathcal{I}_{\beta\beta^T} - \mathcal{I}_{\beta\alpha^T}\mathcal{I}_{\alpha\alpha^T}^{-1}\mathcal{I}_{\alpha\beta^T}.$$

This expression is consistent with that derived in section (4) of[2].

##### 1.3.3 Wald Test

The standard Wald test of the $H_0 : \beta = 0$ is:

$$T_W = \hat{\beta}^T \mathcal{I}_{\beta\beta^T|\alpha} \hat{\beta},$$

which is asymptotically $\chi^2$ with $\dim(\beta)$ degrees of freedom. To test the null hypothesis that a single component $\beta_j = 0$, such as the coefficient for genotype, the test statistic becomes:

$$T_j = (\boldsymbol{e}_j^T \hat{\beta})^T (\boldsymbol{e}_j^T \mathcal{I}_{\beta\beta^T|\alpha}^{-1} \boldsymbol{e}_j)^{-1} (\boldsymbol{e}_j^T \hat{\beta}),$$

where $\boldsymbol{e}_j$ is the $j$th standard basis vector. $T_j$ is asymptotically $\chi_1^2$.

#### 1.4 Analysis of Statistical Efficiency

We next illustrate the benefit of SynSurr relative to a standard analysis in terms of asymptotic relative efficiency (ARE), which compares the variances of these two estimators for $n_{\text{obs}} \to \infty$. First, consider the case when all subjects are complete cases. For ease of exposition, suppose that the covariates have been scaled so that $\boldsymbol{Z}_{\text{obs}}^T \boldsymbol{Z}_{\text{obs}} = n_{\text{obs}} \boldsymbol{I}$. From (11), we have that:

$$\mathcal{I}_{\beta\beta^T|\alpha} = \boldsymbol{Z}_{\text{obs}}^T (\Lambda_{TT} - \Lambda_{TS} \Lambda_{SS}^{-1} \Lambda_{ST}) \boldsymbol{Z}_{\text{obs}} = n_{\text{obs}} \Sigma_{TT}^{-1} \boldsymbol{I}.$$

This is precisely the information from the standard analysis for $\beta$, which implies that if all the subjects have information on the true outcome then SynSurr is asymptotically equivalent to performing a standard analysis.

Now consider the case when there are subjects with missing outcome information and the corresponding covariates have been scaled similarly so that $\boldsymbol{Z}_{\text{miss}}^T \boldsymbol{Z}_{\text{miss}} = n_{\text{miss}} \boldsymbol{I}$ ($n_{\text{miss}} = n - n_{\text{obs}}$). Again from (11), we have that:

$$\mathcal{I}_{\beta\beta^T|\alpha} = n_{\text{obs}} \left\{ \Lambda_{TT} - \Lambda_{TS} \frac{n_{\text{obs}}}{n_{\text{obs}} \Lambda_{SS} + n_{\text{miss}} \Sigma_{SS}^{-1}} \Lambda_{ST} \right\} \boldsymbol{I}$$

and the information from the standard analysis is:

$$\mathcal{I}_{\beta\beta^T} = n_{\text{obs}} \Sigma_{TT}^{-1} \boldsymbol{I}.$$

The ARE of SynSurr relative to the standard analysis is then given by:

$$\text{ARE} = \frac{\mathcal{I}_{\beta\beta^T|\alpha}}{\mathcal{I}_{\beta\beta^T}} = \Sigma_{TT} \left\{ \Lambda_{TT} - \Lambda_{TS} \frac{n_{\text{obs}}}{n_{\text{obs}} \Lambda_{SS} + n_{\text{miss}} \Sigma_{SS}^{-1}} \Lambda_{ST} \right\} \boldsymbol{I} \quad (12)$$

To unpack (12), suppose that $\Sigma$ is a correlation matrix with with $\Sigma_{TT} = \Sigma_{SS} = 1$ and $\Sigma_{TS} = \Sigma_{ST} = \rho \in (-1, 1)$. The ARE can be simplified as

$$\text{ARE} = \frac{1}{1 - \pi_T \rho^2} \boldsymbol{I} \tag{13}$$

where $\pi_T = n_{\text{miss}}/n$. This expression illustrates that the ARE is a function of both the missing rate ($\pi_T$) and the correlation between the synthetic surrogate and the target outcome ($\rho$). If either $\rho$ or $\pi_T$ is zero, then SynSurr maintains the asymptotic efficiency of the standard analysis. In the case of a fixed missing rate, the ARE is maximized at $(1 - \pi_T)^{-1}\boldsymbol{I} = (1+n_{\text{miss}}/n_{\text{obs}})\boldsymbol{I}$ as $\rho \to 1$. Similarly, for a fixed correlation, the ARE increases monotonically to $(1 - \rho^2)^{-1}\boldsymbol{I}$ as $\pi_T \to 1$. Together these properties illustrate the statistical efficiency of SynSurr, which measures the reduction in the variance and the gain in power, increases with increasing $\rho^2$ and an increasing $\pi_T$. These findings coincide with those in[2].

### 2 Imputation simulation

#### 2.1 Supplementary results

Table 1: **Point estimates and analytic and empirical standard errors of the imputation-based estimators compared to SynSurr.** The true value for the parameter of interest is $\beta_G = 0.1$, corresponding to a variant with $h^2 = 1\%$. The Estimate is the mean value of $\hat{\beta}_G$ across $10^3$ simulations; the Analytical SE is the root-mean-square of the estimated SE across simulations; and the Empirical SE is the standard deviation of the point estimates across simulations. The oracle estimator has access to the complete version of $Y$, before 25% of values were set to missing. The standard estimator has access to the observed values of $Y$ only. The imputation-based estimators (SI: single imputation, MI: multiple imputation) impute the missing values of $Y$ using the covariates indicated and an imputation model trained on an independent model-building data set. The imputation model based on $G$ and $X$ is correctly specified, whereas that based on $G$ alone or $X$ alone is misspecified. The SynSurr estimator (SS) jointly analyzes the partially missing $Y$ with the synthetic surrogate $\hat{Y}$; $\hat{Y}$ is generated from a prediction model fit on an independent model-building set using the covariates indicated.

| Estimator | Covariates | Estimate | Analytical SE | Empirical SE |
|---|---|---|---|---|
| Oracle | None | 0.100 | 0.014 | 0.015 |
| Standard | None | 0.100 | 0.016 | 0.016 |
| SI | $(G, X)$ | 0.100 | 0.012 | 0.017 |
| MI | $(G, X)$ | 0.100 | 0.016 | 0.017 |
| SS | $(G, X)$ | 0.100 | 0.016 | 0.016 |
| SI | $(G)$ | 0.012 | 0.012 | 0.015 |
| MI | $(G)$ | 0.012 | 0.017 | 0.016 |
| SS | $(G)$ | 0.100 | 0.016 | 0.016 |
| SI | $(X)$ | 0.075 | 0.012 | 0.012 |
| MI | $(X)$ | 0.075 | 0.016 | 0.012 |
| SS | $(X)$ | 0.100 | 0.016 | 0.016 |

### 3 Robustness simulation

#### 3.1 Simulation methods

Phenotypes were generated from the model:

$$Y = G\beta_G + X\beta_X + \epsilon,$$

with $\epsilon \sim N(0, \sigma^2)$. The coefficients and $\sigma^2$ were chosen such that $G\beta_G$ explained 1% of the variation in $Y$, $X\beta_X$ explained 10% of the variation, and the marginal variance $\mathbb{V}(Y) = 1$. A correlated surrogate outcome was simulated as:

$$\hat{Y} = G\beta_G + X\beta_X + \rho\epsilon + \sqrt{1-\rho^2}\varepsilon,$$

where $\varepsilon \sim N(0, \sigma^2)$ and $\varepsilon$ was independent of $\epsilon$. The residual correlation between $Y$ and $\hat{Y}$ is $\mathbb{C}(Y, \hat{Y}|G, X) = \rho$. Missingness in $Y$ was introduced completely at random. The number of subjects $n_{\text{obs}}$ with observed target phenotypes was fixed at $10^3$, while the number of subjects with missing target phenotypes $n_{\text{miss}}$ was varied to set the missingness:

$$\pi_T = \frac{n_{\text{miss}}}{n_{\text{obs}} + n_{\text{miss}}} \in \{0.00, 0.25, 0.50, 0.75\}.$$

Fixing $n_{\text{obs}}$ and varying $n_{\text{miss}}$ allows for the standard estimator, which is based on the $n_{\text{obs}}$ subjects with observed phenotypes, to serve as a common reference for all of the SynSurr estimators.

#### 3.2 Simulation results

Table 2: **Robustness and precision of the synthetic surrogate estimator with an uninformative and informative synthetic surrogate.** In all cases, the number of subjects with observed phenotypes was $n = 10^3$. The number of subjects with missing phenotypes was varied to achieve the indicated level of missingness $\pi_T$. The standard estimator utilizes the observed values of $Y$ only. Bias is the mean value of $(\hat{\beta}_G - \beta_G)$ across $R = 5 \times 10^3$, SE is the root mean square standard error, and RE is the efficiency relative to standard.

| Estimator | $\rho$ | $\pi_T$ | Bias | SE | RE |
|---|---|---|---|---|---|
| Standard | 0.000 | | -0.001 | 0.030 | 1.000 |
| SynSurr | 0.000 | 0.000 | -0.001 | 0.030 | 0.999 |
| SynSurr | 0.000 | 0.250 | -0.000 | 0.030 | 0.999 |
| SynSurr | 0.000 | 0.500 | -0.000 | 0.030 | 1.000 |
| SynSurr | 0.000 | 0.750 | -0.001 | 0.030 | 1.001 |
| SynSurr | 0.000 | 0.900 | -0.001 | 0.030 | 0.999 |
| Standard | 0.000 | | -0.001 | 0.030 | 1.000 |
| SynSurr | 0.750 | 0.000 | -0.001 | 0.030 | 0.999 |
| SynSurr | 0.750 | 0.250 | -0.000 | 0.028 | 1.163 |
| SynSurr | 0.750 | 0.500 | 0.000 | 0.025 | 1.393 |
| SynSurr | 0.750 | 0.750 | -0.001 | 0.023 | 1.732 |
| SynSurr | 0.750 | 0.900 | -0.000 | 0.021 | 2.024 |

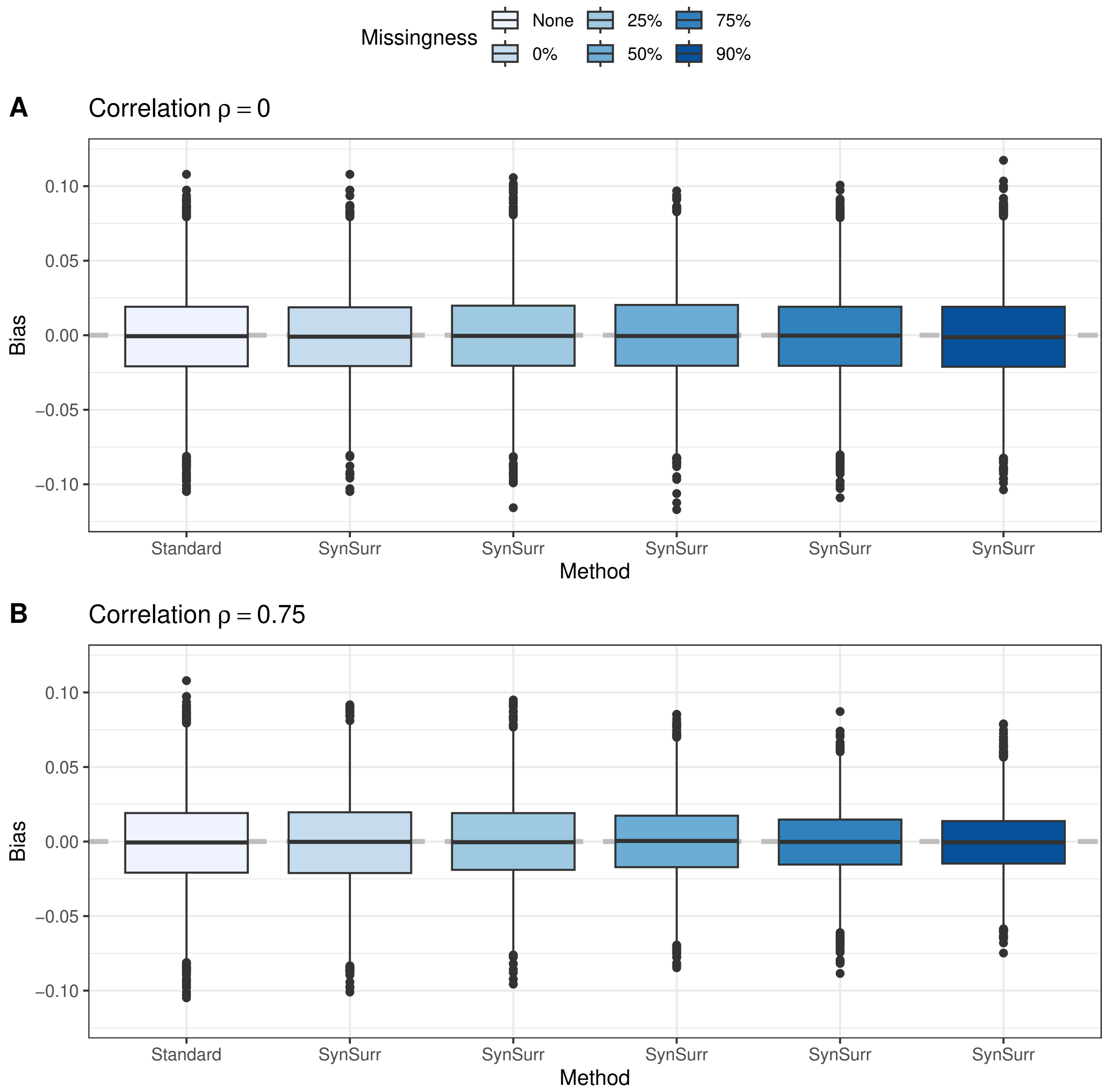

Figure 1: **Robustness and precision of SynSurr with an uninformative and informative synthetic surrogate.** In all cases, the number of subjects with observed phenotypes was $n = 10^3$. The number of subjects with missing phenotypes was varied to achieve the indicated level of missingness. The standard estimator utilizes the observed values of $Y$ only. In panel **A**, the synthetic surrogate has correlation $\rho = 0.00$ with the target phenotype, and is in fact independent of the target phenotype. Use of the SynSurr estimator with this uninformative surrogate results in no loss of efficiency relative to the standard analysis. In panel **B**, the synthetic surrogate has correlation $\rho = 0.75$ with the target phenotype. SynSurr becomes more efficient as the number of subjects with missing target outcomes increases. The number of simulation replicates is $R = 5 \times 10^3$.

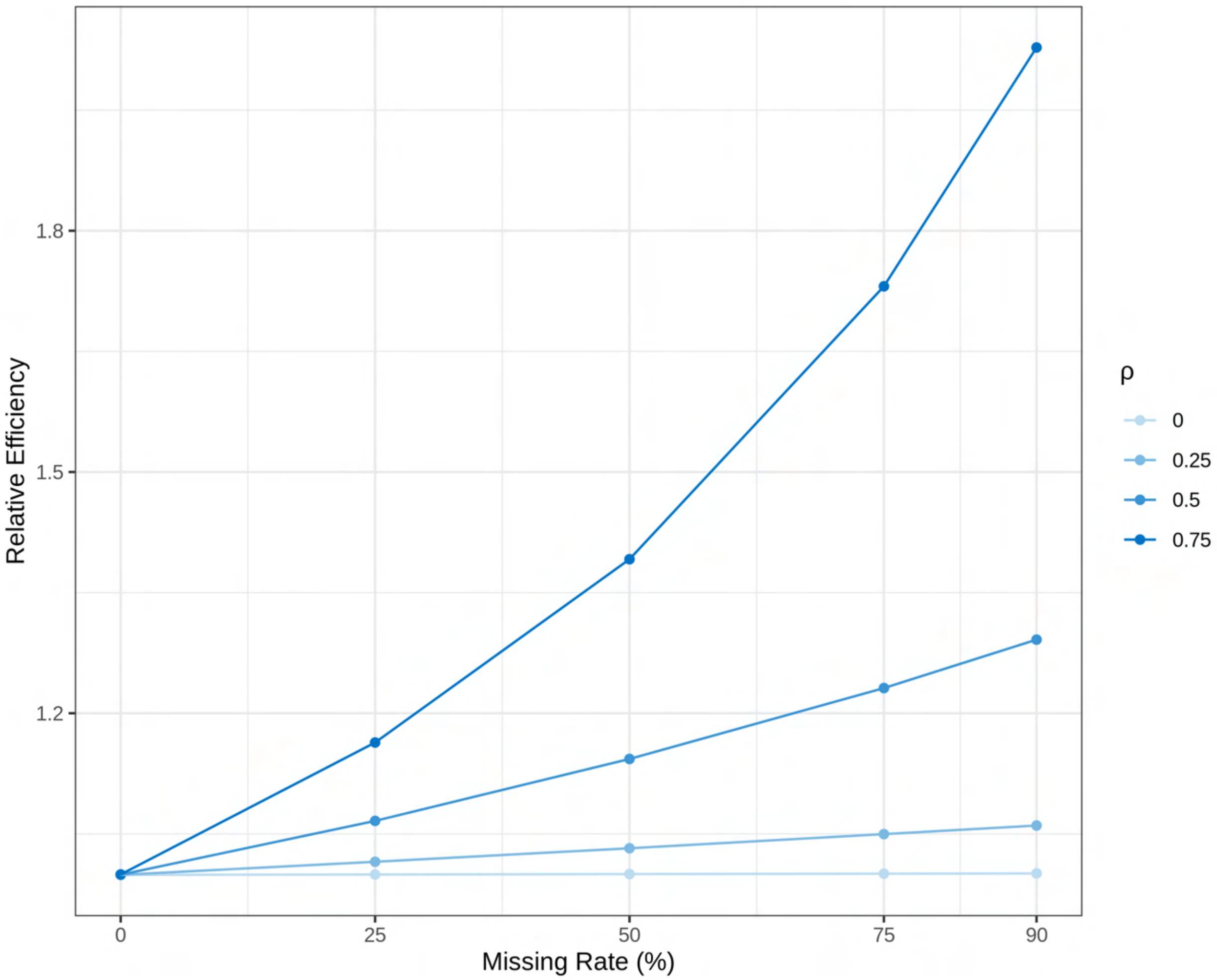

Figure 2: **Efficiency of SynSurr relative to the standard estimator across various missing rates and synthetic surrogates at a SNP heritability of 0.05%.** In all cases, the number of subjects with observed phenotypes was $n = 10^3$. The number of subjects with missing phenotypes was varied to achieve the indicated level of missingness. The synthetic surrogate has correlation $\rho \in \{0.00, 0.25, 0.50, 0.75\}$ with the target phenotype. When there is no missingness, SynSurr is equivalent to the standard estimator. The efficiency of SynSurr increases with increasing missingness rate and correlation. The number of simulation replicates is $10^3$.

### 4 Type I error and power simulations

Here and throughout the manuscript, the null hypothesis of interest is that genotype has no effect on the target outcome $H_0 : \beta_G = 0$. Note that, whether genotype has an effect on the synthetic surrogate (i.e. $H_0 : \alpha_G = 0$) is not of scientific interest. The type I error is defined as the probability of incorrectly rejecting the null hypothesis when the null hypothesis is true, and power is defined as the probability of correctly rejecting the null hypothesis when the null hypothesis is false.

#### 4.1 Supplementary results

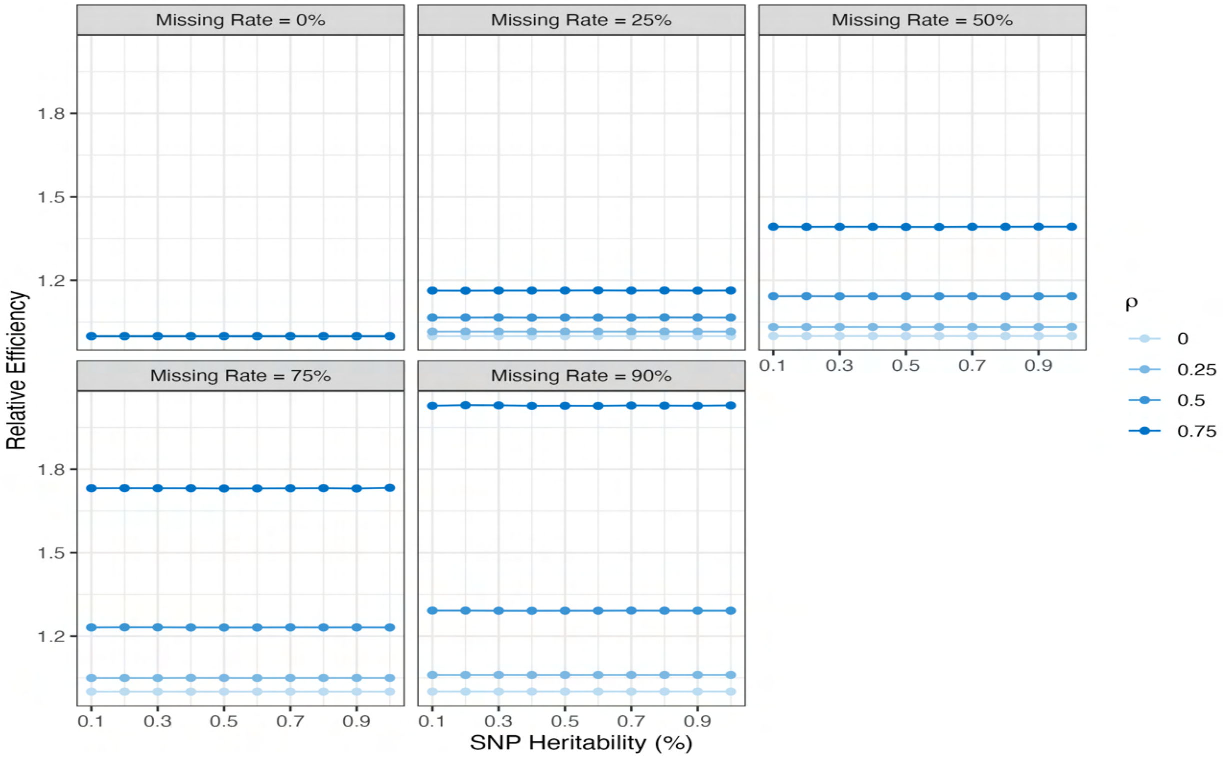

Figure 3: **Efficiency of SynSurr relative to the standard analysis against across various missing rates, synthetic surrogates, and values of SNP heritability.** In all cases, the number of subjects with observed phenotypes was $n = 10^3$. The number of subjects with missing phenotypes was varied to achieve the indicated level of missingness. In each panel, the synthetic surrogate has correlation $\rho = 0.00, 0.25, 0.50, 0.75$ with the target phenotype and the SNP heritability was varied from 0.1% to 1%. When there is no missingness, SynSurr is asymptotically equivalent to the standard analysis. The efficiency of SynSurr increases with increasing missing rate and correlation, but is maintained across different values of SNP heritability. The number of simulation replicates is $R = 10^3$.

Table 3: **Type I error and power of SynSurr across various missing rates and synthetic surrogates.** In all cases, the number of subjects with observed phenotypes was $n = 10^3$, while the number with missing phenotypes was varied to achieve the indicated level of missingness. The synthetic surrogate has correlation $\rho = 0.00, 0.25, 0.50, 0.75$ with the target phenotype. The power is reported for the setting with SNP heritability of 0.5% ($\beta_G = 0.07$). Type I error is controlled across all settings. The power is stable across values of $\rho$ when there is no missing phenotype information, which is asymptotically equivalent to the standard analysis. The power of SynSurr increases with increasing missing rate and correlation. The number of simulation replicates is $R = 10^5$.

| Missing Rate (%) | $\rho$ | Type I Error | $\chi^2$ | Power | $\chi^2$ |
|---|---|---|---|---|---|
| 0 | 0.00 | 0.05 | 1.01 | 0.65 | 6.58 |
| 0 | 0.25 | 0.05 | 1.00 | 0.65 | 6.59 |
| 0 | 0.50 | 0.05 | 0.99 | 0.65 | 6.56 |
| 0 | 0.75 | 0.05 | 1.00 | 0.65 | 6.59 |
| 25 | 0.00 | 0.05 | 0.99 | 0.66 | 6.59 |
| 25 | 0.25 | 0.05 | 1.00 | 0.66 | 6.69 |
| 25 | 0.50 | 0.05 | 1.00 | 0.68 | 6.96 |
| 25 | 0.75 | 0.05 | 1.00 | 0.72 | 7.52 |
| 50 | 0.00 | 0.05 | 1.01 | 0.65 | 6.56 |
| 50 | 0.25 | 0.05 | 1.00 | 0.67 | 6.76 |
| 50 | 0.50 | 0.05 | 0.99 | 0.71 | 7.37 |
| 50 | 0.75 | 0.05 | 1.00 | 0.79 | 8.77 |
| 75 | 0.00 | 0.05 | 1.00 | 0.65 | 6.56 |
| 75 | 0.25 | 0.05 | 1.00 | 0.68 | 6.89 |
| 75 | 0.50 | 0.05 | 1.00 | 0.75 | 7.90 |
| 75 | 0.75 | 0.05 | 1.00 | 0.87 | 10.68 |
| 90 | 0.00 | 0.05 | 1.00 | 0.66 | 6.58 |
| 90 | 0.25 | 0.05 | 1.00 | 0.68 | 6.84 |
| 90 | 0.50 | 0.05 | 1.01 | 0.76 | 8.29 |
| 90 | 0.75 | 0.05 | 0.99 | 0.92 | 12.2 |

Table 4: **Number of significant SNPs recovered by standard and SynSurr across various missing rates and target-surrogate correlations.** In all cases, the SNP heritability was 0.5% ($\beta_G = 0.07$), the number of subjects with observed phenotypes was $n = 10^3$, while the number with missing phenotypes was varied to achieve the indicated level of missingness. The rate shown in parentheses is the percentage of oracle variants recovered.

| Missing Rate (%) | $\rho$ | Oracle | Standard | SynSurr |
|---|---|---|---|---|
| 0 | 0.00 | 6533 | 6533(100%) | 6532(99.98%) |
| 0 | 0.25 | 6623 | 6623(100%) | 6618(99.92%) |
| 0 | 0.50 | 6534 | 6534(100%) | 6530(99.94%) |
| 0 | 0.75 | 6509 | 6509(100%) | 6506(99.95%) |
| 25 | 0.00 | 7814 | 6306(80.7%) | 6311(80.77%) |
| 25 | 0.25 | 7788 | 6363(81.7%) | 6401(82.19%) |
| 25 | 0.50 | 7833 | 6283(80.21%) | 6590(84.13%) |
| 25 | 0.75 | 7756 | 6290(81.1%) | 6928(89.32%) |
| 50 | 0.00 | 9199 | 6516(70.83%) | 6518(70.86%) |
| 50 | 0.25 | 9182 | 6474(70.51%) | 6589(71.76%) |
| 50 | 0.50 | 9176 | 6441(70.19%) | 6989(76.17%) |
| 50 | 0.75 | 9172 | 6449(70.31%) | 7901(86.14%) |
| 75 | 0.00 | 9970 | 6508(65.28%) | 6509(65.29%) |
| 75 | 0.25 | 9976 | 6604(66.2%) | 6812(68.28%) |
| 75 | 0.50 | 9973 | 6611(66.29%) | 7511(75.31%) |
| 75 | 0.75 | 9976 | 6584(66%) | 8759(87.8%) |
| 90 | 0.00 | 10000 | 6606(66.06%) | 6613(66.13%) |
| 90 | 0.25 | 10000 | 6564(65.64%) | 6798(67.98%) |
| 90 | 0.50 | 10000 | 6550(65.5%) | 7674(76.74%) |
| 90 | 0.75 | 10000 | 6588(65.88%) | 9221(92.21%) |

### 5 Sample reuse simulation

The objective of this simulation is to evaluate whether reuse of the same subjects for model-building and for GWAS invalidates statistical inference, either by biasing the estimated genetic effect or by inflating the type I error.

#### 5.1 Simulation methods

Phenotypes were generated from the model:

$$Y = G\beta_G + X\beta_1 + X^2\beta_2 + X^3\beta_3 + \epsilon, \tag{14}$$

with $\epsilon \sim N(0, \sigma^2)$. The coefficients and $\sigma^2$ were selected such that each power of $X$ explained 10% of the variation in $Y$, and the marginal variance was $\mathbb{V}(Y) = 1$. The correlation between genotype $G$ and the covariate $X$ was 0.5, such that $X$ confounds the association between $G$ and $Y$. For type I error simulations, $G$ explained 0% of the variation in $Y$, while for bias simulations $G$ explained 1%. As described previously, the number of subjects with observed target phenotypes was fixed at $10^3$, while the number of subjects with missing target phenotypes was varied to set the missingness. The synthetic surrogate $\hat{Y}$ was generated from a linear regression model of the form $\hat{Y}(X) = B_k(X)^T\gamma$, for $k \in \{2, 3\}$, where $B_k(X)$ is a B-spline basis expansion of order $k$. The linear regression was fit either to an independent data set of size $10^3$, or to the labeled part of the GWAS data set. In either case, SynSurr GWAS was performed using the association model:

$$\begin{bmatrix} Y \\ \hat{Y} \end{bmatrix} \big| (G, X) = \begin{bmatrix} \beta_0 + G\beta_G + X\beta_X \\ \alpha_0 + G\alpha_G + X\alpha_X \end{bmatrix} + \begin{bmatrix} \epsilon_T \\ \epsilon_S \end{bmatrix}. \tag{15}$$

Note that the true dependence of $Y$ on $X$ in (14) is cubic while the association model (15) only adjusts for a linear term.

#### 5.2 Simulation results

**Supplemental Figure (4)** demonstrates that reusing the same data for training the synthetic surrogate model and performing GWAS neither biases the estimated effect size $\hat{\beta}_G$ nor inflates the expected $\chi^2$ under the null (i.e. the type I error). Panel A considers a "mis-specified" surrogate model ($k = 2$) that can only capture quadratic dependence of $Y$ on $X$, while Panel B considers a "correctly specified" surrogate model ($k = 3$) that can capture cubic dependence. As seen here and in the main text, correct specification of the surrogate model is not required for the validity of SynSurr. Overall, these results suggest that statistical inference remains valid even when the same data are utilized for model-building and for GWAS. Also see the real-data subject reuse experiment reported in **Supplemental Tables 14-15**.

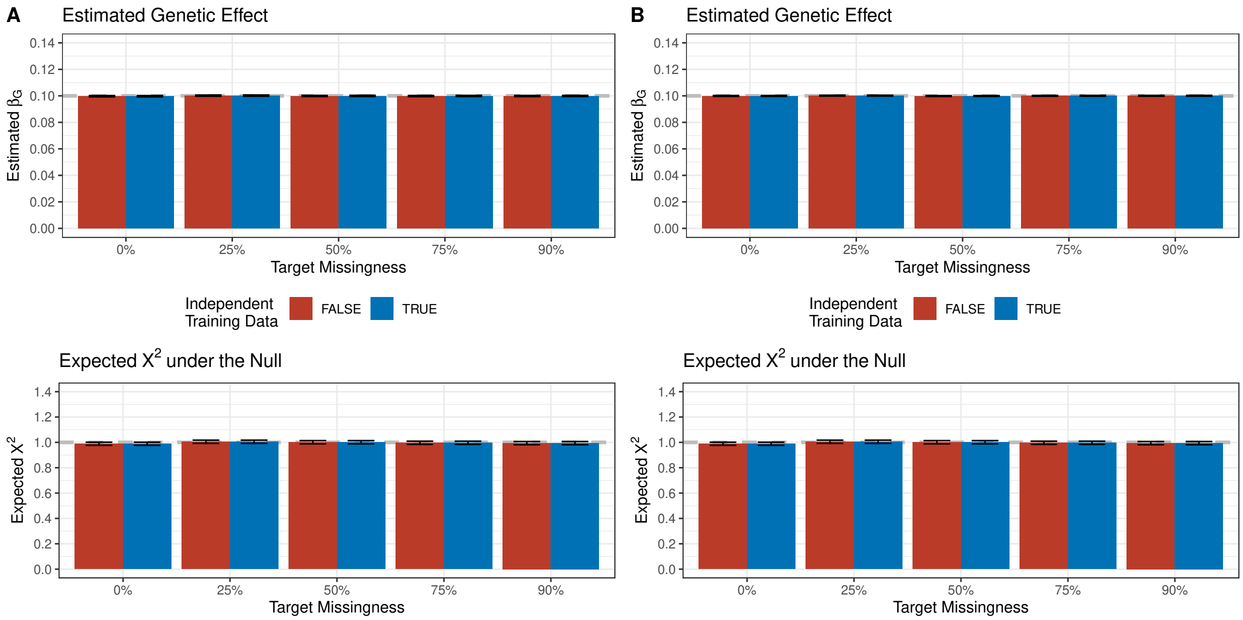

Figure 4: **SynSurr remains unbiased and properly controls the type I error when the same data are utilized for model training and for GWAS.** The number of subjects with observed phenotypes was $n = 10^3$, while the number with missing phenotypes was varied to achieve the indicated level of missingness. The model that generated the synthetic surrogate was either trained in the GWAS data set or in an independent data set of size $n = 10^3$. Shown are the estimated effect size across $20 \times 10^3$ simulation replicates, with a true genetic effect of $\beta_G = 0.1$, and the average $\chi^2$ statistic under $H_0 : \beta_G = 0$ across $50 \times 10^3$ simulation replicates, for which the expected value is 1.0. Panel A considers a "misspecified" ($k = 2$) model that can only capture quadratic dependence of $Y$ on $X$, while Panel B considers a "correctly specified" model ($k = 3$) that can capture the cubic dependence. As seen, the validity of SynSurr is not contingent on correct specification of the surrogate model.

### 6 Sample splitting simulation

The objective of this simulation is to examine the trade-off between allocating subjects to the model-building data set, to improve the quality of the synthetic surrogate, versus allocating subjects to the GWAS data set, to improve power, given that the model-building and GWAS data sets will be kept independent.

#### 6.1 Simulation methods

Phenotypes were generated from the model:

$$Y = G\beta_G + \sin(X)\beta_S + \cos(X)\beta_C + \sin(X)\cos(X)\beta_{SC} + \epsilon, \tag{16}$$

with $\epsilon \sim N(0, \sigma^2)$. The coefficients and $\sigma^2$ were selected such that genotype $G$ explained 1% of the variation in $Y$, each transformation of $X$ explained 10% of the variation, and $\mathbb{V}(Y) = 1$. The correlation between the covariate $X$ and the predictor $G$ was 0.5, such that $X$ confounds the association between $G$ and $Y$. The data set was simulated to contain $2 \times 10^2$ independent subjects in total, of which $n_{\text{obs}} = 10^3$ had observed target outcomes and $n_{\text{miss}} = 10^3$ had missing target outcomes. The subjects with missing target outcomes were all allocated to the GWAS data set. A proportion $p_{\text{train}} \in \{0.05, 0.10, 0.15, 0.20, 0.25\}$ of subjects with observed target outcomes were randomly allocated to a model-building data set, and the remainder to the GWAS data set. The synthetic surrogate $\hat{Y}$ was generated from a linear regression model of the form $\hat{Y}(X) = B_3(X)^T\gamma$, where $B_3(X)$ is a B-spline basis expansion of order 3. The coefficient $\gamma$ of the synthetic surrogate model was estimated using the $n_{\text{obs}} \cdot p_{\text{train}}$ subjects in the model-building data set. Finally, SynSurr GWAS was performed using the association model:

$$\begin{bmatrix} Y \\ \hat{Y} \end{bmatrix} \big| (G, X) = \begin{bmatrix} \beta_0 + G\beta_G + X\beta_X \\ \alpha_0 + G\alpha_G + X\alpha_X \end{bmatrix} + \begin{bmatrix} \epsilon_T \\ \epsilon_S \end{bmatrix}.$$

Note that neither the model which generates the synthetic surrogate nor the association model can fully capture the complex dependence of $Y$ on $X$ in the generative model (16).

#### 6.2 Simulation results

Notwithstanding the results of the sample reuse simulation, in some settings having independent model-building and GWAS data sets may be desirable. **Supplemnetal Figure (5)** evaluates the trade-off between allocating more subjects to the model-building data set versus retaining those subjects for use in the GWAS when a fixed number of subjects with observed target outcomes are available. Panel A underscores that as more subjects are allocated to model-building, fewer are available for GWAS. Panel B verifies that, as expected, the unbiasedness of SynSurr is not contingent on the allocation of subjects. Panel C demonstrates that as more subjects are allocated to model-building, the correlation between the target outcome $Y$ and the synthetic surrogate $\hat{Y}$ in the GWAS data set improves. However, Panel D indicates that the increase in target-surrogate correlation achieved by allocating more subjects to model-building does not compensate for the decrease in size of the GWAS data set: power ultimately decreases as more subjects are allocated to model-building.

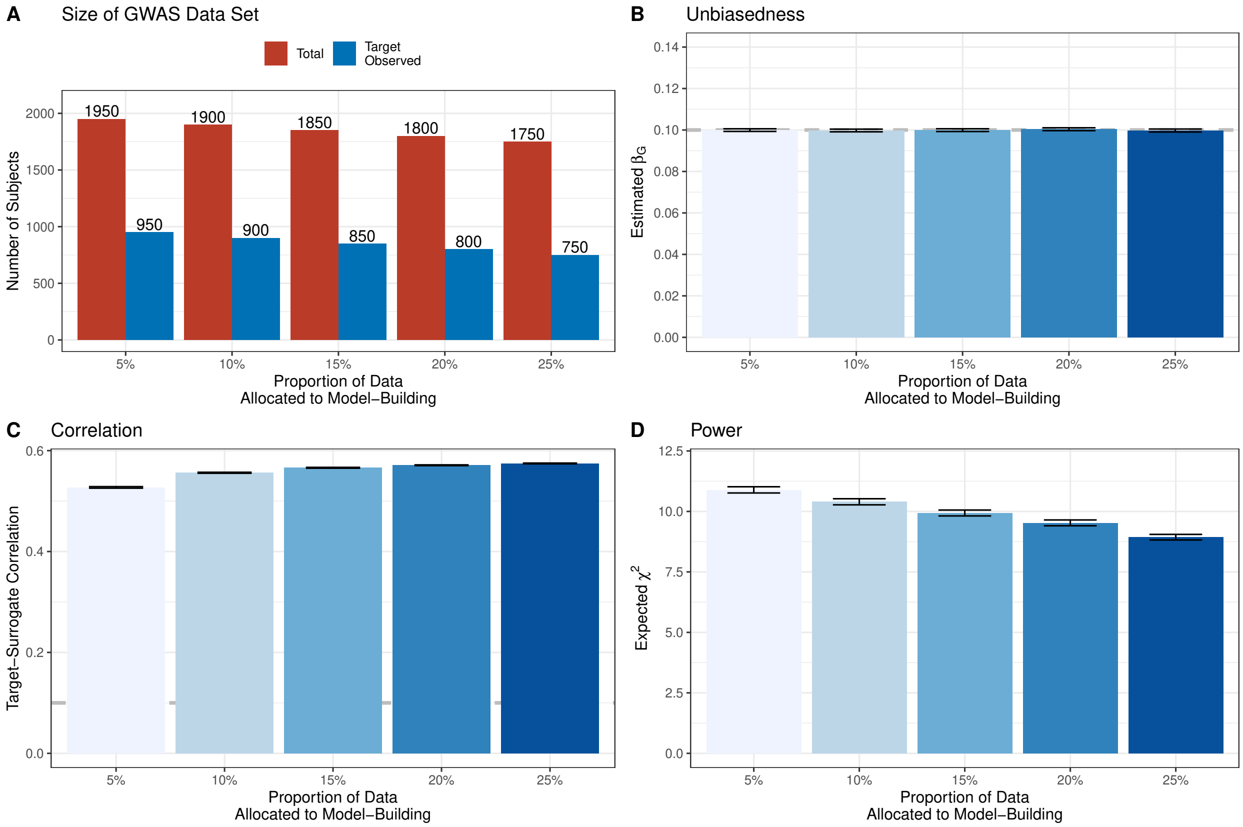

Figure 5: **Trade-off between allocating subjects to the model-building data set versus retaining subjects for use in GWAS.** In total, $10^3$ subjects with observed target outcomes and $10^3$ subjects with missing target outcomes were available. Among subjects with observed target outcomes, a proportion was randomly allocated to the model-building data set, and the remainder to the GWAS data set. A. Number of subjects in the GWAS data set. B. Estimated effect size across $10^4$ simulation replicates. The true effect size was $\beta_G = 0.1$, corresponding to a variant with $h^2 = 1\%$. C. Correlation between $Y$ and $\hat{Y}$ among subjects with observed target outcomes in the GWAS data set. D. Average $\chi^2$ statistic at associated variants across $10^4$ simulation replicates. A larger value indicates greater power to detect an association.

### 7 Predictor imputation simulation

The main text considers imputation of the target outcome $Y$. The objective of this simulation is to examine the validity of imputing a predictor $Z$ that serves as an input to the model that generates the synthetic surrogate $\hat{Y}$.

#### 7.1 Simulation methods

Phenotypes were generated from the model:

$$Y = G\beta_G + X\beta_X + Z\beta_1 + Z^2\beta_2 + Z^3\beta_3 + \epsilon, \tag{17}$$

with $\epsilon \sim N(0, \sigma^2)$. The coefficients and $\sigma^2$ were selected such that $X$ and each power of $Z$ explained 10% of the variation in $Y$, with $\mathbb{V}(Y) = 1$. For type I error simulations, $G$ explained 0% of the variation in $Y$, while for bias simulations $G$ explained 1%. The correlation between the covariate $X$ and $G$ was 0.5, as was the correlation between $X$ and $Z$. As described previously, the number of subjects with observed target phenotypes was fixed at $10^3$, while the number of subjects with missing target phenotypes was varied. Within the GWAS data set, $Z$ was randomly set to missing for a proportion $\pi_Z \in \{0.00, 0.125, 0.250, 0.375, 0.500\}$ of subjects. Cases with both $Y$ and $Z$ missing were admitted. In an independent training data set of size $10^3$, two models were trained. The first was a linear model of the form $\hat{Z}(X) = X\alpha$ to impute $Z$ from $X$. The second was a linear model of the form $\hat{Y}(Z) = B_3(Z)^T\gamma$ to generate the synthetic surrogate from $Z$. In the GWAS data set, define the imputed predictor as:

$$Z^* = \begin{cases} Z, & \text{if } Z \text{ is observed,} \\ \hat{Z}(X), & \text{if } Z \text{ is missing.} \end{cases}$$

With $Z^*$, the synthetic surrogate was generated as $\hat{Y} = \hat{Y}(Z^*)$. Finally, SynSurr GWAS was performed using the association model:

$$\begin{bmatrix} Y \\ \hat{Y} \end{bmatrix} \Big| (G, X) = \begin{bmatrix} \beta_0 + G\beta_G + X\beta_X \\ \alpha_0 + G\alpha_G + X\alpha_X \end{bmatrix} + \begin{bmatrix} \epsilon_T \\ \epsilon_S \end{bmatrix}. \tag{18}$$

#### 7.2 Simulation results

**Supplemental Figure (6)** demonstrates that generating the synthetic surrogate $\hat{Y}$ from a covariate $Z^*$ that was imputed (via $X$) neither biases the estimated effect size nor inflates the type I error. While generating $\hat{Y}$ from imputed data may distort the relationship between genotype $G$ and $\hat{Y}$ (which is not of scientific interest), it is not expected to affect the relationship between $G$ and $Y$, as seen empirically.

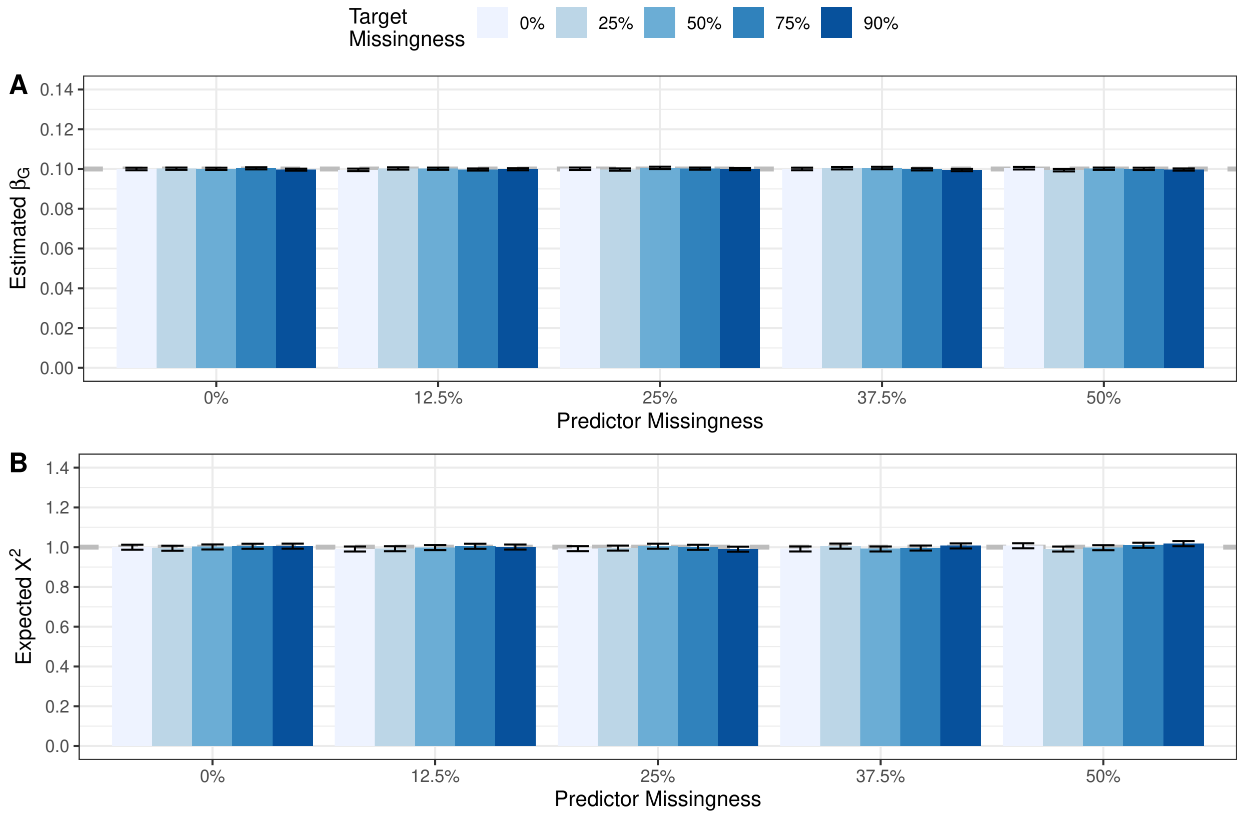

Figure 6: **SynSurr remains unbiased and properly controls the type I error when the synthetic surrogate is generated using imputed predictors.** The number of subjects with observed phenotypes was $n = 10^3$, while the number with missing phenotypes was varied to achieve the indicated level of target missingness. The synthetic surrogate depended on a partially missing predictor $Z$ that was imputed using $X$ prior to generation of $\hat{Y}$. A. Estimated effect size across $10 \times 10^3$ simulation replicates. The true effect size was $\beta_G = 0.1$, corresponding to a variant with $h^2 = 1\%$. B. Average $\chi^2$ statistic under the null $\beta_G = 0$ across $20 \times 10^3$ simulation replicates. In the absence of inflation, the expected value is 1.0.

### 8 UK Biobank (UKBB) genotype quality control (QC)

Table 5: **Quality control of the UK biobank genotypes.** The initial data set contained 488,377 subjects and 784,256 directly genotyped genetic markers. Prior to the analysis, we removed SNPs(i) with < 90% genotyping rate (GR) (ii) that failed the Hardy-Weinberg equilibrium (HWE) test at $p < 10^{-5}$, and (iii) with minor allele frequency (MAF) <1%. CHR: chromosome.

| CHR | Initial count | GR < 90% | HWE < $10^{-5}$ | MAF <1% | Final count |
|---|---|---|---|---|---|
| 1 | 63487 | 4089 | 16010 | 8722 | 34666 |
| 2 | 61966 | 4598 | 16067 | 6381 | 34920 |
| 3 | 52300 | 3640 | 13016 | 5757 | 29887 |
| 4 | 47443 | 3018 | 12594 | 3932 | 27899 |
| 5 | 46314 | 3699 | 11456 | 4056 | 27103 |
| 6 | 53695 | 4166 | 14867 | 4974 | 29688 |
| 7 | 42722 | 2658 | 11081 | 4845 | 24138 |
| 8 | 38591 | 2205 | 10173 | 3416 | 22797 |
| 9 | 34310 | 2429 | 8861 | 3811 | 19209 |
| 10 | 38308 | 2772 | 9887 | 3730 | 21919 |
| 11 | 40824 | 3113 | 9782 | 6139 | 21790 |
| 12 | 37302 | 2432 | 9437 | 4570 | 20863 |
| 13 | 26806 | 3062 | 6682 | 1827 | 15235 |
| 14 | 25509 | 1681 | 6600 | 3165 | 14063 |
| 15 | 24467 | 1544 | 6855 | 2878 | 13190 |
| 16 | 28960 | 1985 | 7679 | 4379 | 14917 |
| 17 | 28835 | 3231 | 6933 | 4828 | 13843 |
| 18 | 21962 | 1432 | 5794 | 1546 | 13190 |
| 19 | 26186 | 2703 | 6233 | 5958 | 11292 |
| 20 | 19959 | 1171 | 5228 | 2126 | 11434 |
| 21 | 11342 | 727 | 3079 | 998 | 6538 |
| 22 | 12968 | 771 | 3542 | 1768 | 6887 |
| Total | 784256 | 57126 | 201856 | 89806 | 435468 |

### 9 UKBB Ablation Analysis

#### 9.1 Random forest modeling

Sets of first, second, and third-degree relatives were identified based on the kinship coefficient. One subject was allocated to the GWAS data set, and the remaining subjects to the model-building data set. For height (Data-Field 50), 55,618 subjects were allocated to the model-building data set. A random forest model was trained to predict height based on age (Data-Field 21022), sex (Data-Field 22001), body weight (Data-Field 21002), and waist circumference (Data-Field 48). The predictions from the random forest model were used as the synthetic surrogates in the association analyses. For FEV1 (Data-Field 20150), 42,031 subjects were allocated to the model-building data set. The random forest model included, age, sex, height, waist circumference, hip circumference (Data-Field 49), smoking status (Data-Field 1239), body mass index (Data-Field 23104), basal metabolic rate (Data-Field 23105), and whole body impedance (Data-Field 23106).

Random forest models were fit using the `ranger` (Version 0.15.1) package in `R` with default parameter settings. Notably, the forest includes 500 trees, the number of variables examined at each split is the square root of the number of input variables, the minimum node size is 5, and the splitting criteria is the decrease in node variance.

Figure 7: **Predicted vs. observed height within the model-building and GWAS data sets.** A random forest was trained to predict height using 55,618 subjects allocated to the model-building data set. Model inputs included age, sex, body weight, and waist circumference.

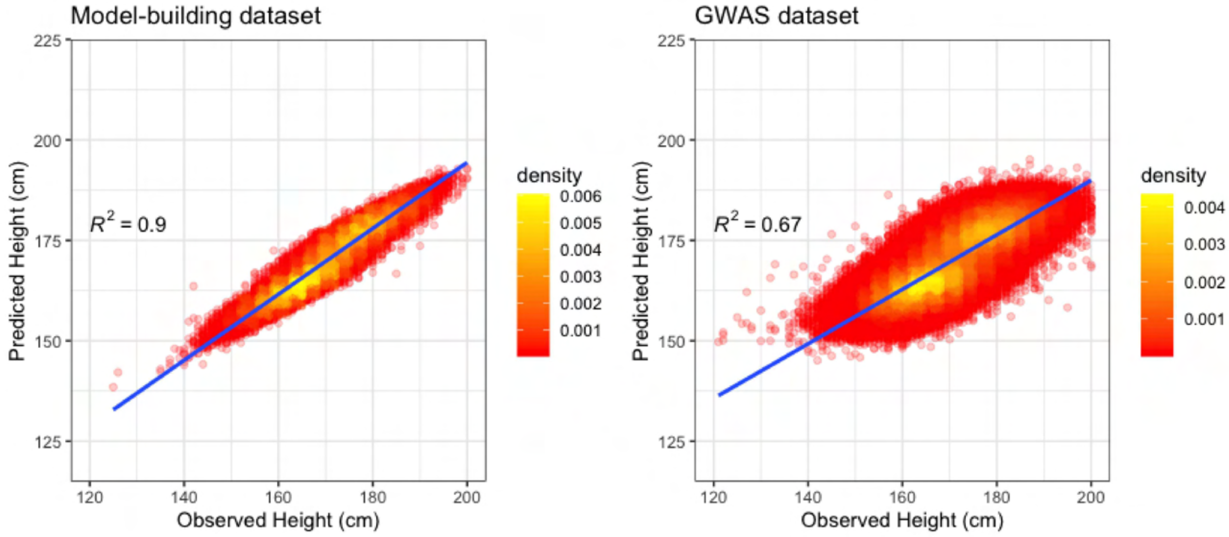

Figure 8: **Predicted vs. observed FEV1 within the model-building and GWAS data sets.** A random forest was trained to predict FEV1 using 42,031 subjects allocated to the model-building data set. Model inputs included age, sex, height, waist circumference, hip circumference, smoking status, body mass index, basal metabolic rate, and whole body impedance.

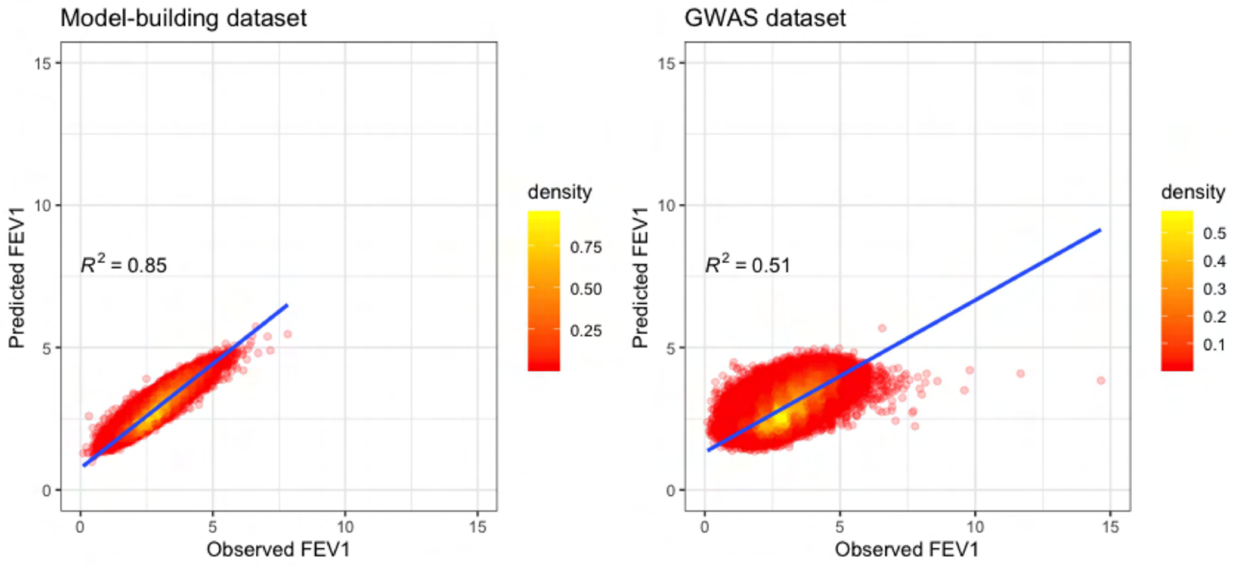

##### 9.1.1 Height and FEV1

Table 6: **False positive discoveries of the standard GWAS and the SynSurr GWAS across various missing rates.** An association was considered a false positive if it was not detected by the oracle GWAS (standard GWAS in the absence of missingness).

| | Height | | FEV1 | |
|---|---|---|---|---|
| Missing Rate(%) | Standard | SynSurr | Standard | SynSurr |
| 0 | 0(0%) | 1(0%) | 0(0%) | 0(0%) |
| 25 | 75(1.51%) | 111(2.23%) | 19(3.36%) | 17(3.01%) |
| 50 | 18(0.65%) | 32(1.15%) | 6(2.11%) | 15(5.28%) |
| 75 | 1(0.12%) | 7(0.83%) | 0(0%) | 0(0%) |
| 90 | 0(0%) | 0(0%) | 0(0%) | 0(0%) |

Table 7: **Average mean squared standard error (MSE) across the set of genome-wide significant associations identified by the oracle analysis.** The oracle method has access to the target phenotype before the introduction of missingness. The average MSE is computed across the set of variants declared significant ($p < 5 \times 10^{-8}$) by the oracle analysis. The relative efficiency is defined as the ratio of the SynSurr MSE to standard MSE.

| | Height | | | | FEV1 | | | |
|---|---|---|---|---|---|---|---|---|
| | MSE | | | | MSE | | | |
| Missing Rate(%) | Oracle | Standard | SynSurr | Relative Efficiency | Oracle | Standard | SynSurr | Relative Efficiency |
| 0 | 9.7e-06 | 9.7e-06 | 9.7e-06 | 1.00 | 9.5e-06 | 9.5e-06 | 9.5e-06 | 1.00 |
| 25 | 9.7e-06 | 1.3e-05 | 1.2e-05 | 1.08 | 9.5e-06 | 1.3e-05 | 1.2e-05 | 1.04 |
| 50 | 9.7e-06 | 1.9e-05 | 1.7e-05 | 1.16 | 9.5e-06 | 1.9e-05 | 1.8e-05 | 1.09 |
| 75 | 9.7e-06 | 3.9e-05 | 3.1e-05 | 1.26 | 9.5e-06 | 3.8e-05 | 3.4e-05 | 1.14 |
| 90 | 9.7e-06 | 9.8e-05 | 7.4e-05 | 1.33 | 9.5e-06 | 9.7e-05 | 8.3e-05 | 1.17 |

Table 8: **Average $\chi^2$ across the set of genome-wide significant associations identified by the oracle analysis.** The oracle method has access to the target phenotype before the introduction of missingness. The average $\chi^2$ is computed across the set of variants declared significant ($p < 5 \times 10^{-8}$) by the oracle analysis. The relative efficiency is defined as the ratio of the SynSurr $\chi^2$ to the standard $\chi^2$.

| | Height | | | | FEV1 | | | |
|---|---|---|---|---|---|---|---|---|
| | Avergae $\chi^2$ | | | | Avergae $\chi^2$ | | | |
| Missing Rate(%) | Oracle | Standard | SynSurr | Relative Efficiency | Oracle | Standard | SynSurr | Relative Efficiency |
| 0 | 67.59 | 67.59 | 67.59 | 1.00 | 51.27 | 51.27 | 51.27 | 1.00 |
| 25 | 67.59 | 50.95 | 54.51 | 1.07 | 51.27 | 38.58 | 40.27 | 1.04 |
| 50 | 67.59 | 34.37 | 39.44 | 1.15 | 51.27 | 26.14 | 28.48 | 1.09 |
| 75 | 67.59 | 17.55 | 21.78 | 1.24 | 51.27 | 13.18 | 14.84 | 1.13 |
| 90 | 67.59 | 7.92 | 10.09 | 1.27 | 51.27 | 5.51 | 6.00 | 1.09 |

Figure 9: **Signal recovery of SynSurr relative to the oracle GWAS for height and FEV1.** A slope of 1.0 indicates that the estimated effect sizes are consistent with the oracle effect sizes. Note that although the slope deviates from 1.0 at 90% missingness, the slope approaches 1.0 as missingness declines. The following figure, which assesses signal recovery for standard GWAS, provides a point of comparison for the $R^2$ values.

(a) Height

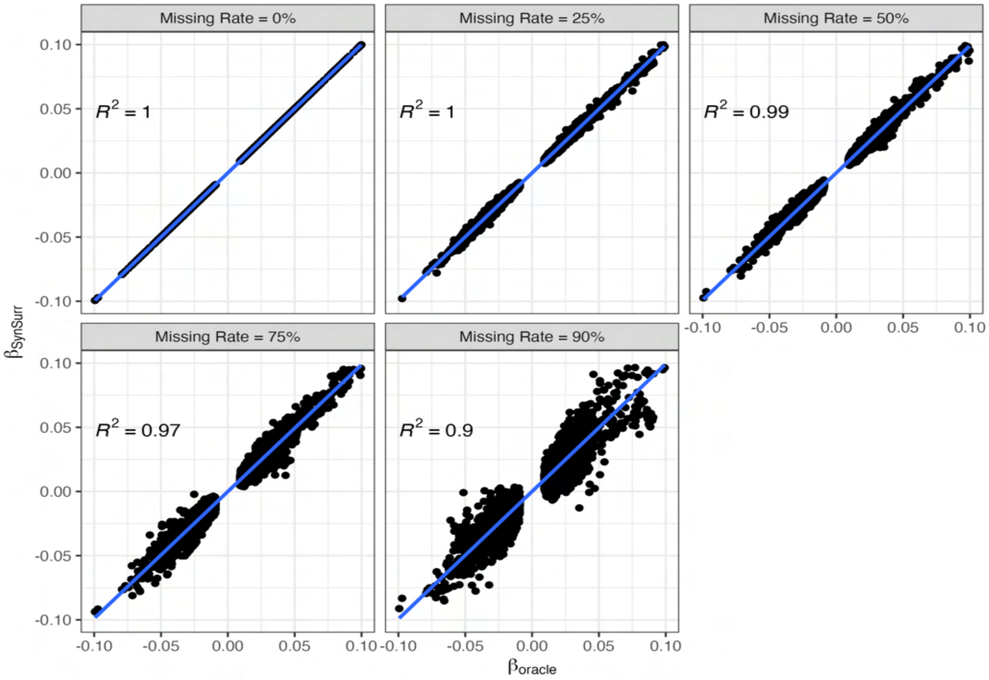

(b) FEV1

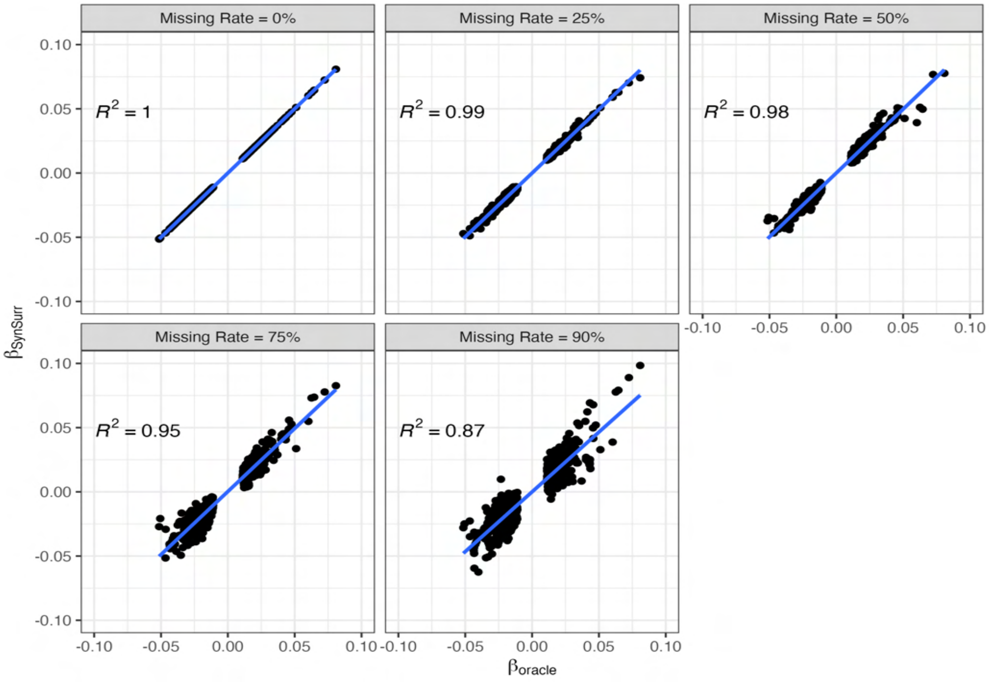

Figure 10: **Signal recovery of standard GWAS relative to the oracle GWAS for height and FEV1.** A slope of 1.0 indicates that the estimated effect sizes are consistent with the oracle effect sizes. Note that although the slope deviates from 1.0 at 90% missigness, the slope approaches 1.0 as missingness declines.

(a) Height

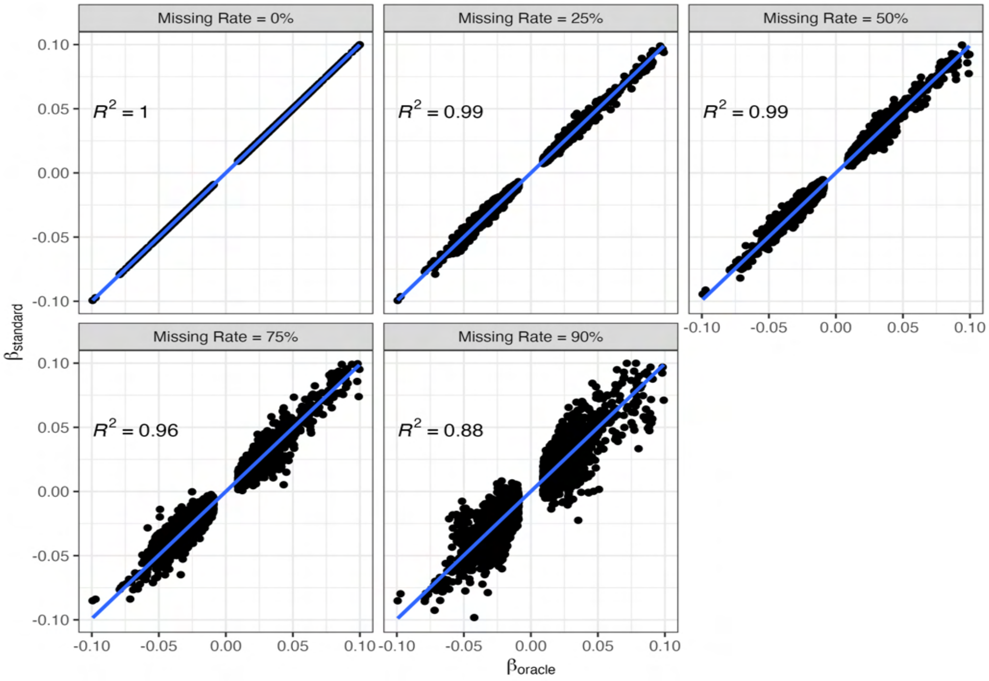

(b) FEV1

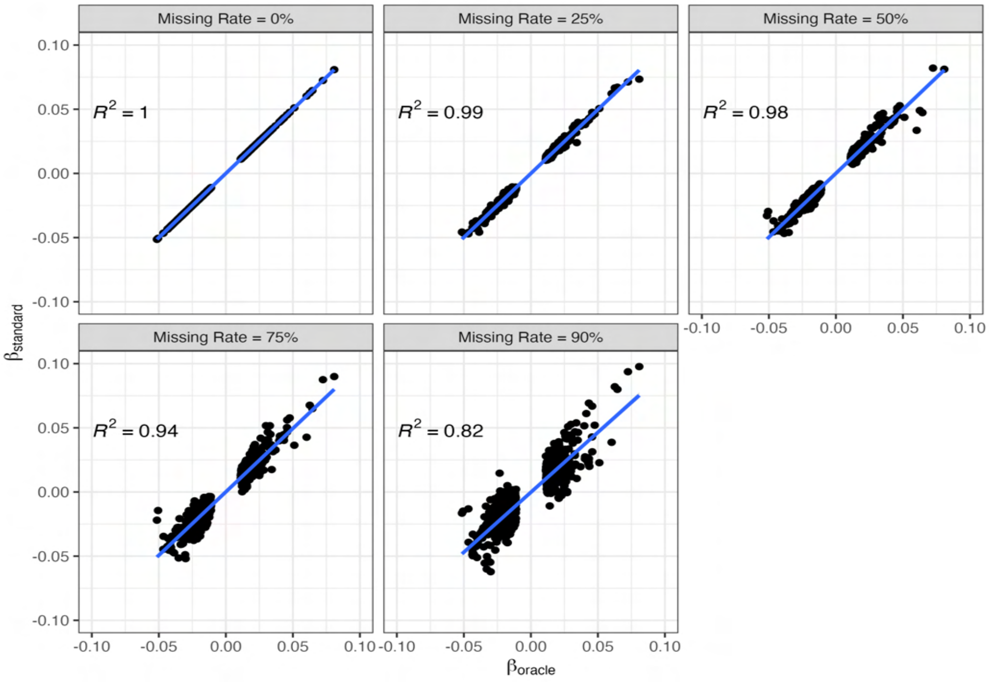

#### 9.2 Comparison of SynSurr with imputation-based GWAS

Table 9: **Target-surrogate correlation in the GWAS data set.** Surrogates/imputations were generated either from a random forest (RF) or from linear regression. Permuted surrogates/imputations were generated by shuffling the RF values, while negated surrogate/imputations were generated by multiplying the RF values by $-1$.

| Phenotype | Random Forest | Linear Regression | Permuted | Negated |
|---|---|---|---|---|
| Height | 0.79 | 0.37 | 0.00 | -0.79 |
| FEV1 | 0.75 | 0.19 | 0.00 | -0.75 |

##### 9.2.1 Height

Table 10: **Number of genome-wide significant SNPs recovered by the standard GWAS, imputation-based GWAS and the SynSurr GWAS for height with 50% missingness.** The oracle method establishes the number of genome-wide significant (GWS) variants ($p < 5 \times 10^{-8}$) that would be identified in the absence of missingness. Surrogates/imputations were generated either from a random forest (higher-quality) or a linear regression (lower-quality). In addition, cases where the random forest surrogate was either permuted (i.e. uninformative) or negated (i.e. adversarial) are considered. The recovery rate, shown in parentheses, is the percentage of oracle variants recovered. The false negative rate is 100% minus the oracle recovery rate.

| Surrogate Model | Oracle | Standard GWAS | Single Imputation | Multiple Imputation | SynSurr |
|---|---|---|---|---|---|
| Random Forest | 7177 | 2742(38.21%) | 3678(51.24%) | 685(9.54%) | 3421(47.67%) |
| Linear Regression | 7177 | 2742(38.21%) | 2424(33.77%) | 143(1.99%) | 2728(38.01%) |
| Permuted | 7177 | 2742(38.21%) | 519(7.23%) | 38(0.53%) | 2717(37.86%) |
| Negated | 7177 | 2742(38.21%) | 0(0.00%) | 0(0.00%) | 3421(47.67%) |

Table 11: **False positive discoveries of the standard GWAS, imputation-based GWAS, and the SynSurr GWAS for height with 50% missingness.** An association was considered a false positive if it was not detected by the oracle GWAS (standard GWAS in the absence of missingness). The denominator for calculating the false discovery rate is the total number of genome-wide significant associations.

| Surrogate Model | Standard GWAS | Single Imputation | Multiple Imputation | SynSurr |
|---|---|---|---|---|
| Random Forest | 18 (0.65%) | 103 (2.72%) | 0 (0%) | 32 (0.92%) |
| Linear Regression | 18 (0.65%) | 15 (0.62%) | 0 (0%) | 18 (0.66%) |
| Permuted | 18 (0.65%) | 5 (0.95%) | 0 (0%) | 16 (0.59%) |
| Negated | 18 (0.65%) | 0 (0%) | 0 (0%) | 32 (0.92%) |

Figure 11: **Signal recovery of imputation-based approaches and SynSurr relative to the oracle GWAS for height with 50% missingness.** A slope of 1.0 indicates that the estimated effect sizes are consistent with the oracle effect sizes, whereas a slope deviating from 1.0 suggests the presence of bias.

(a) Surrogates/imputations generated by random forest

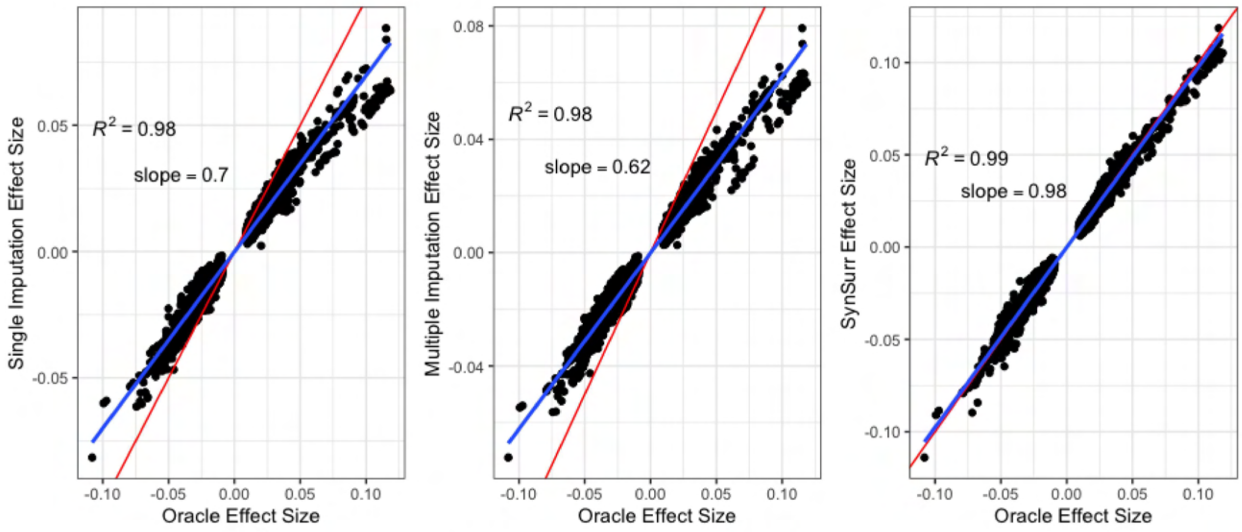

(b) Surrogates/imputations generated by linear regression

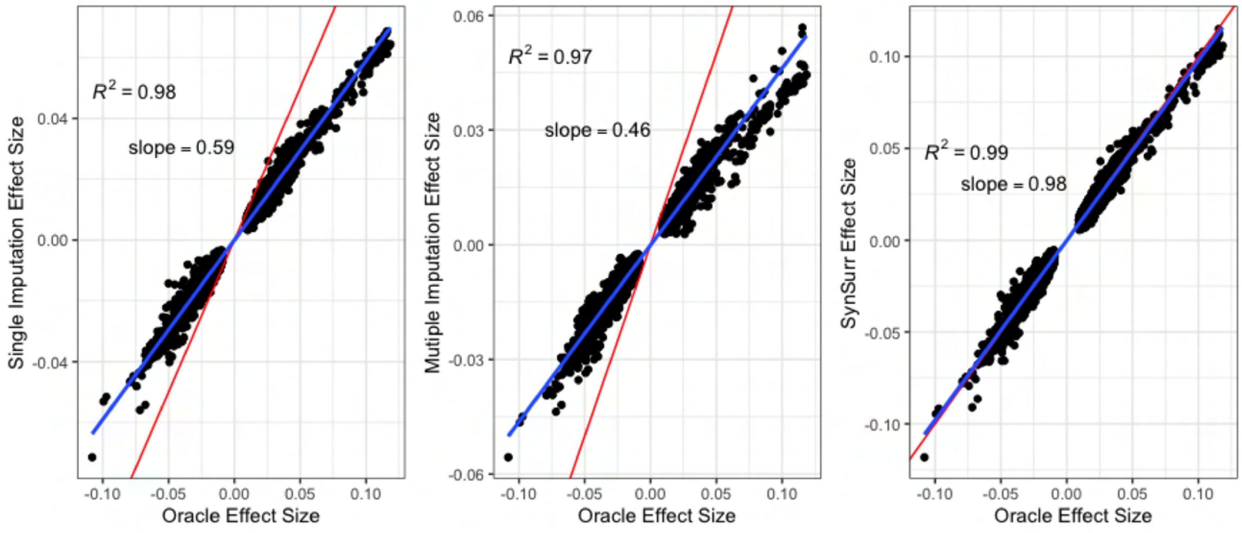

(c) Surrogates/imputations generated by random forest then permuted

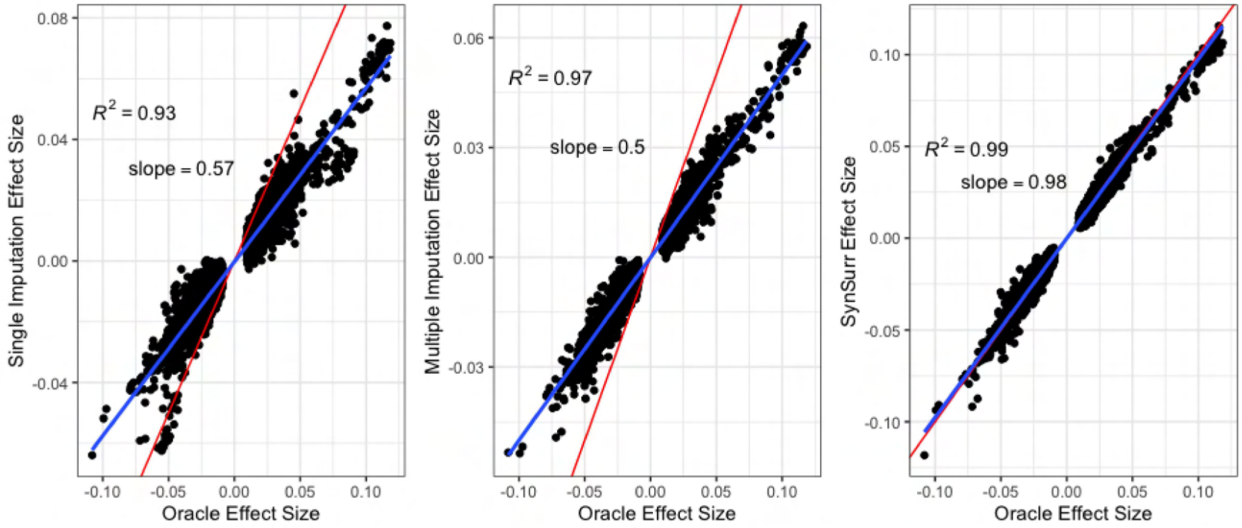

(d) Surrogates/imputations generated by random forest then negated

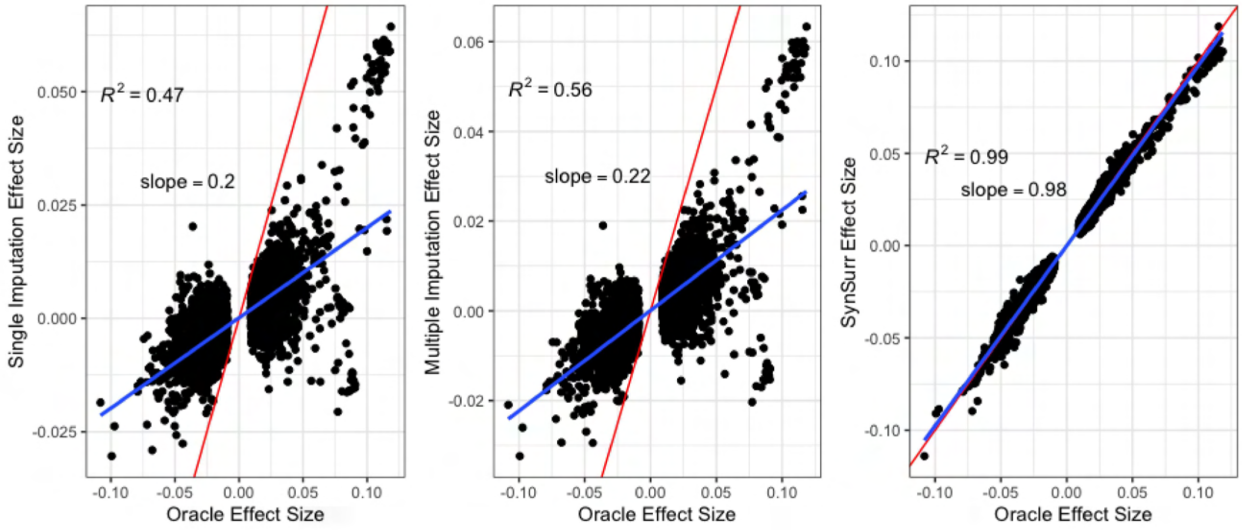

##### 9.2.2 FEV1

Table 12: **Number of genome-wide significant SNPs recovered by the standard GWAS, imputation-based GWAS and the SynSurr GWAS for FEV1 with 50% missingness.** The oracle method establishes the number of genome-wide significant (GWS) variants ($p < 5 \times 10^{-8}$) that would be identified in the absence of missingness. Surrogates/imputations were generated either from a random forest (higher-quality) or a linear regression (lower-quality). In addition, cases where the random forest surrogate was either permuted (i.e. uninformative) or negated (i.e. adversarial) are considered. The recovery rate, shown in parentheses, is the percentage of oracle variants recovered. The false negative rate is 100% minus the oracle recovery rate.

| Surrogate Model | Oracle | Standard GWAS | Single Imputation | Multiple Imputation | SynSurr |
|---|---|---|---|---|---|
| Random Forest | 974 | 278(28.54%) | 612(62.83%) | 56(5.78%) | 326(33.47%) |
| Linear Regression | 974 | 278(28.54%) | 246(25.26%) | 32(3.29%) | 261(26.80%) |
| Permuted | 974 | 278(28.54%) | 16(1.64%) | 0(0.00%) | 256(26.28%) |
| Negated | 974 | 278(28.54%) | 0(0.00%) | 0(0.00%) | 326(33.47%) |

Table 13: **False positive discoveries of the standard GWAS, imputation-based GWAS, and the SynSurr GWAS FEV1 with 50% missingness.** An association was considered a false positive if it was not detected by the oracle GWAS (standard GWAS in the absence of missingness). The denominator for calculating the false discovery rate is the total number of genome-wide significant associations.

| Surrogate Model | Standard GWAS | Single Imputation | Multiple Imputation | SynSurr |
|---|---|---|---|---|
| Random Forest | 6 (2.12%) | 412 (40.24%) | 8 (12.50%) | 15 (4.39%) |
| Linear Regression | 6 (2.12%) | 3 (1.20%) | 0 (0%) | 2 (0.79%) |
| Permuted | 6 (2.12%) | 0 (0%) | 0 (0%) | 2 (0.78%) |
| Negated | 6 (2.12%) | 0 (0%) | 0 (0%) | 15 (4.39%) |

Figure 12: **Signal recovery of imputation-based approach and SynSurr relative to the oracle GWAS for FEV1 with 50% missingness.** A slope of 1.0 indicates that the estimated effect sizes are consistent with the oracle effect sizes, whereas a slope deviating from 1.0 suggests the presence of bias.

(a) Surrogates/imputations generated by random forest

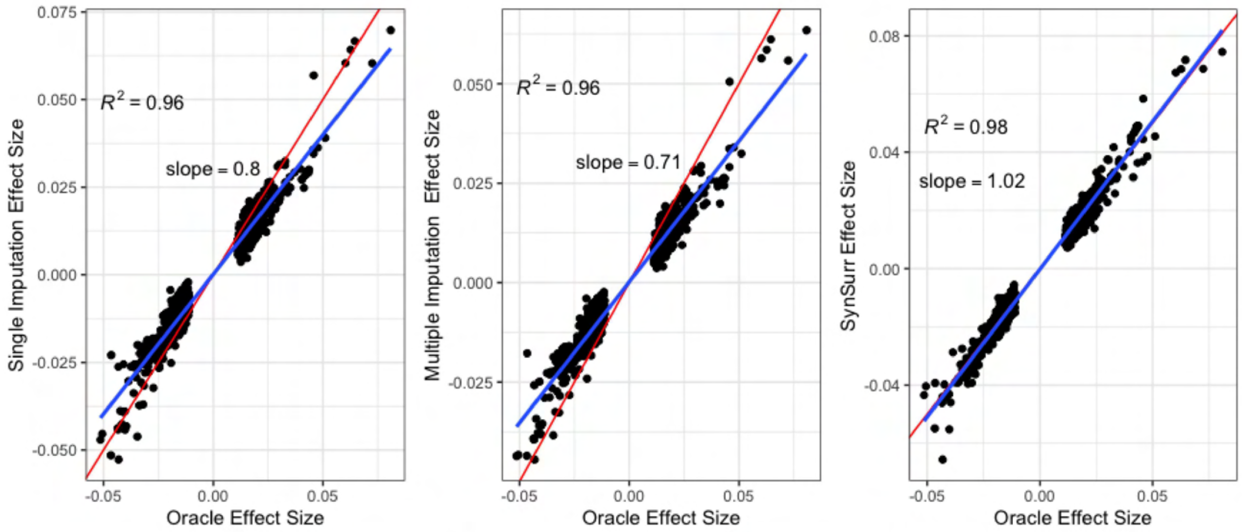

(b) Surrogates/imputations generated by linear regression

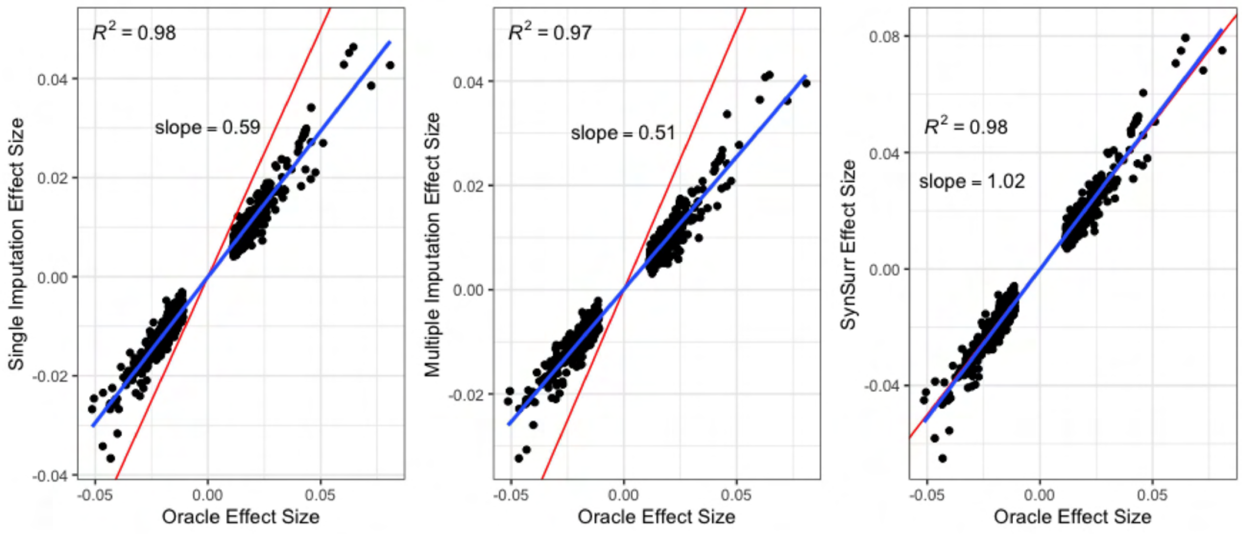

(c) Surrogates/imputations generated by random forest then permuted

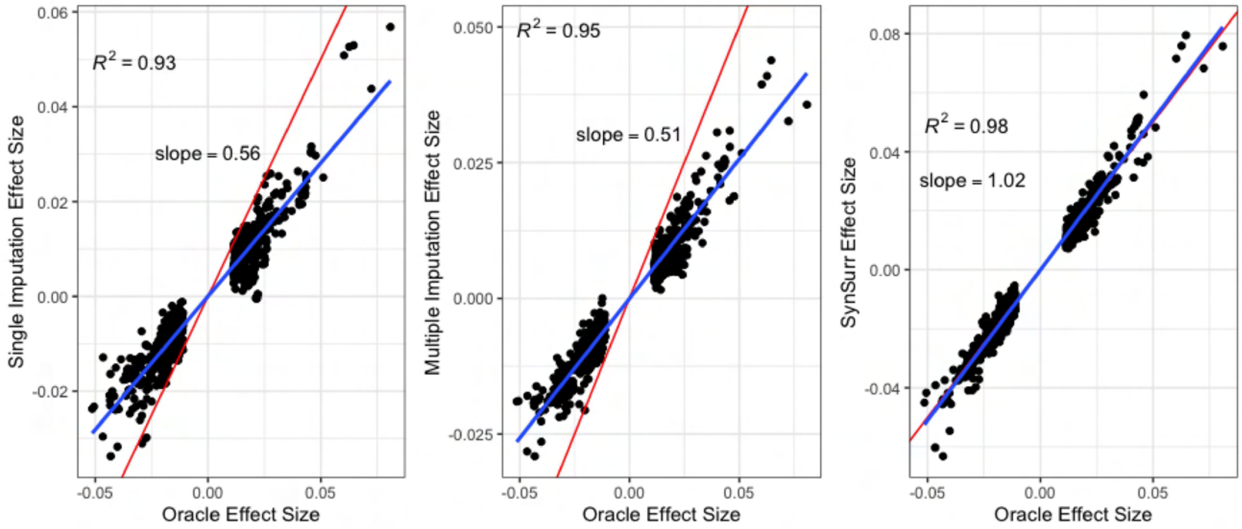

(d) Surrogates/imputations generated by random forest then negated

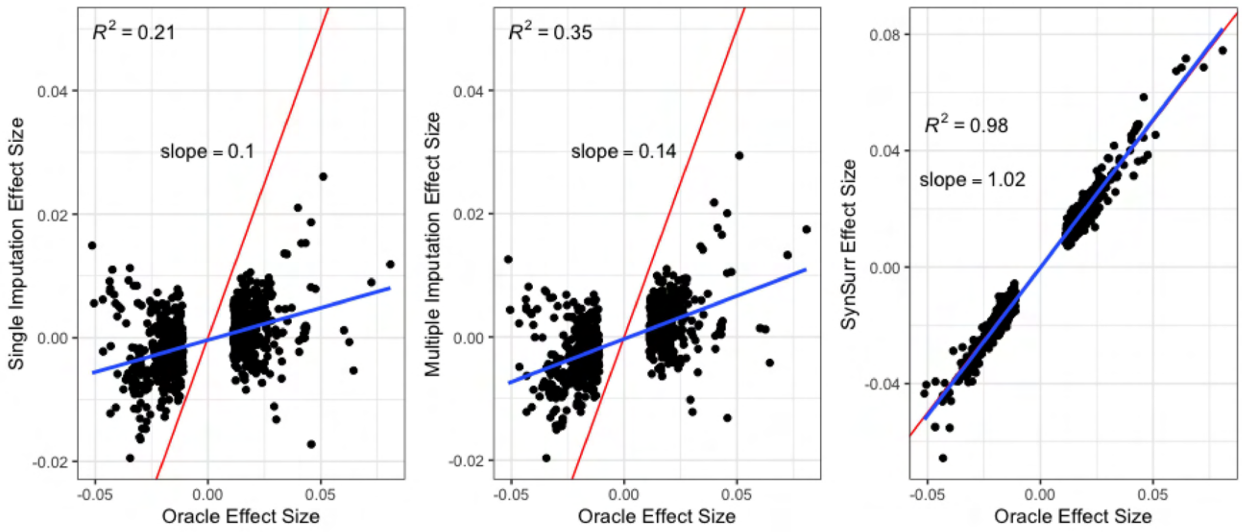

#### 9.3 SynSurr sample reuse

This section provides an empirical counterpart to the sample reuse simulation study in supplemental section 5. For these experiments, the same data were utilized for both model-building and for GWAS. SynSurr maintains a false positive rate directly comparable to that of standard GWAS while providing improved power. Importantly, the increase is recovery of oracle variants with SynSurr (i.e power) is substantially greater than the negligible increase in the false positive rate.

Table 14: **Number of genome-wide significant SNPs recovered by standard and SynSurr GWAS across increasing ablation of the target phenotype.** The same data were used for training the model that generates the synthetic surrogate and for running GWAS. The oracle method establishes the number of genome-wide significant (GWS) variants ($p < 5 \times 10^{-8}$) that would be identified in the absence of missingness. Missingness was introduced by ablating 25%, 50%, 75%, and 90% of the target phenotypes.

| | Height | | | | FEV1 | | | |
|---|---|---|---|---|---|---|---|---|
| Missing Rate (%) | $n_{obs}$ | Oracle | Standard | SynSurr | $n_{obs}$ | Oracle | Standard | SynSurr |
| 25 | 262,105 | 7,177 | 4,896(68.22%) | 5,169(72.02%) | 231,388 | 974 | 546(56.06%) | 613(62.94%) |
| 50 | 174,737 | 7,177 | 2,742(38.21%) | 3,319(46.24%) | 154,259 | 974 | 278(28.54%) | 305(31.31%) |
| 75 | 87,368 | 7,177 | 834(11.62%) | 1,325(18.46%) | 77,129 | 974 | 60(6.16%) | 75(7.70%) |
| 90 | 34,947 | 7,177 | 192(2.68%) | 320(4.46%) | 30,852 | 974 | 0(0%) | 2(0.21%) |

Table 15: **False positive discoveries of the standard GWAS and the SynSurr GWAS across various missing rates.** The same data were used for training the model that generates the synthetic surrogate and for running GWAS. An association was considered a false positive if it was not detected by the oracle GWAS (standard GWAS in the absence of missingness). The denominator for calculating the false discovery rate is the total number of genome-wide significant associations. Crucially, although the same data have been utilized for model-building and for GWAS, the false positive rate of SynSurr is directly comparable to that of standard GWAS.

| | Height | | FEV1 | |
|---|---|---|---|---|
| Missing Rate(%) | Standard | SynSurr | Standard | SynSurr |
| 25 | 75(1.51%) | 82(1.56%) | 19(3.03%) | 22(3.46%) |
| 50 | 18(0.65%) | 27(0.81%) | 6(2.12%) | 7(2.24%) |
| 75 | 1(0.12%) | 4(0.30%) | 0(0%) | 2(2.60%) |
| 90 | 0(0%) | 1(0.31%) | 0(0%) | 0(0%) |

#### 9.4 Comparison of SynSurr with Proxy and MTAG GWAS

Table 16: **Comparison of SynSurr with Proxy and MTAG GWAS for height.** Proxy GWAS analyzes the synthetic surrogate $\hat{Y}$ in place of the true target outcome $Y$. MTAG augments the results from standard GWAS, for a given level of missingness, with those from proxy GWAS. An association was considered a true positive if it was identified by the oracle GWAS (i.e. standard GWAS in the absence of missingness), and a false positive if it was not identified by the oracle GWAS. Below, "Total" is the total number of genome-wide significant associations, "True Positives" gives the number and percentage of oracle variants recovered, and "False Positives" gives the number and percentage of the total associations that were not detected by the oracle.

| Method | Missing (%) | Oracle | Total | True Positives | False Positives |
|---|---|---|---|---|---|
| Proxy | 0 | 7,177 | 1,654 | 1,363(18.99%) | 291(17.59%) |
| MTAG | 0 | 7,177 | 2,981 | 2,981(41.54%) | 0(0%) |
| SynSurr | 0 | 7,177 | 7,177 | 7,177(100%) | 0(0%) |
| MTAG | 25 | 7,177 | 2,384 | 2,384(33.22%) | 0(0%) |
| SynSurr | 25 | 7,177 | 5,416 | 5,305(73.92%) | 111(2.05%) |
| MTAG | 50 | 7,177 | 1,703 | 1,698(23.66%) | 5(0.29%) |
| SynSurr | 50 | 7,177 | 3,453 | 3,421(47.67%) | 32(0.93%) |
| MTAG | 75 | 7,177 | 1,075 | 1,057(14.73%) | 18(1.67%) |
| SynSurr | 75 | 7,177 | 1,250 | 1,243(17.32%) | 7(0.56%) |
| MTAG | 90 | 7,177 | 832 | 788(10.98%) | 44(5.29%) |
| SynSurr | 90 | 7,177 | 329 | 329(4.58%) | 0(0%) |

Table 17: **Comparison of SynSurr with Proxy and MTAG GWAS for FEV1.** Proxy GWAS analyzes the synthetic surrogate $\hat{Y}$ in place of the true target outcome $Y$. MTAG augments the results from standard GWAS, for a given level of missingness, with those from proxy GWAS. An association was considered a true positive if it was identified by the oracle GWAS (i.e. standard GWAS in the absence of missingness), and a false positive if it was not identified by the oracle GWAS. Below, "Total" is the total number of genome-wide significant associations, "True Positives" gives the number and percentage of oracle variants recovered, and "False Positives" gives the number and percentage of the total associations that were not detected by the oracle.

| Method | Missing (%) | Oracle | Total | True Positives | False Positives |
|---|---|---|---|---|---|
| Proxy | 0 | 974 | 3,185 | 429(44.05%) | 2,756(86.53%) |
| MTAG | 0 | 974 | 834 | 674(69.2%) | 160(19.18%) |
| SynSurr | 0 | 974 | 974 | 974(100%) | 0(0%) |
| MTAG | 25 | 974 | 666 | 483(49.59%) | 183(27.48%) |
| SynSurr | 25 | 974 | 616 | 599(61.5%) | 17(2.76%) |
| MTAG | 50 | 974 | 595 | 345(35.42%) | 250(42.02%) |
| SynSurr | 50 | 974 | 341 | 326(33.47%) | 15(4.4%) |
| MTAG | 75 | 974 | 597 | 250(25.67%) | 347(58.12%) |
| SynSurr | 75 | 974 | 82 | 81(8.32%) | 1(1.22%) |
| MTAG | 90 | 974 | 650 | 213(21.87%) | 437(67.23%) |
| SynSurr | 90 | 974 | 0 | 0(0%) | 0(0%) |

### 10 UKBB DEXA Phenotype Analysis

#### 10.1 Random forest modeling

For the 6 body composition phenotypes measured by DEXA scan (total android mass [Data-Field 23248], total arm mass [Data-Field 23260], total gynoid mass [Data-Field 23265], total leg mass [Data-Field 23277], total body mass [Data-Field 23283], and total trunk mass [Data-Field 23287]), 4,584 subjects were allocated to the model-building data set. A random forest model was trained to predict each body composition phenotype on the basis of age, sex, height, body weight, body mass index, and 5 measurements of impedance: whole body (Data-Field 23106), left arm (Data-Field 23110), right arm (Data-Field 23109), left leg (Data-Field 23108), right leg (Data-Field 23107). Random forest models were fit in `ranger` using default parameter settings as described in the above section on ablation analyses.

Figure 13: **Predicted vs. observed values of body composition phenotypes within the model-building and GWAS data sets.** A random forest was trained to predict each of the 6 body composition phenotypes, obtained via DEXA scan, using 4,584 subjects allocated to the model-building data set. Model inputs included age, sex, height, body weight, body mass index, and 5 impedance measures (whole body, left arm, right arm, left leg and right leg).

(a) Android Total Mass

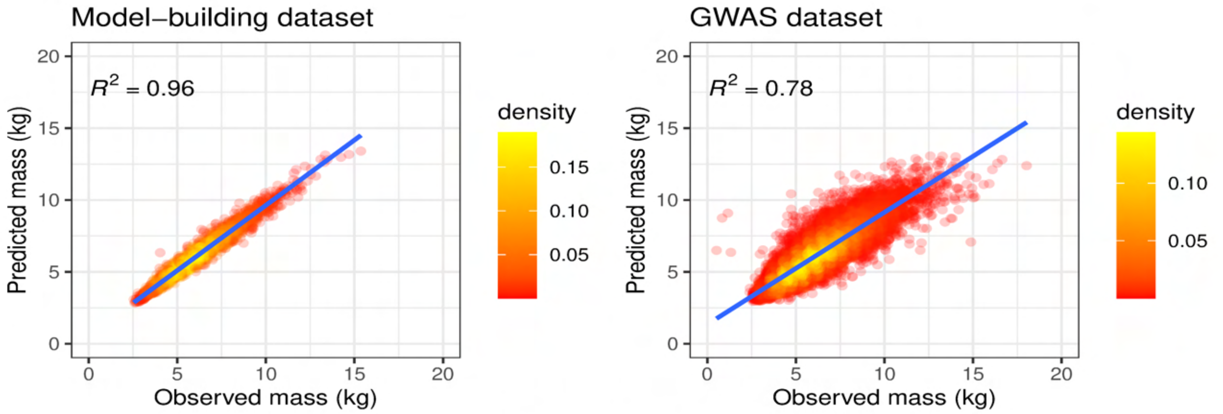

(b) Arms Total Mass

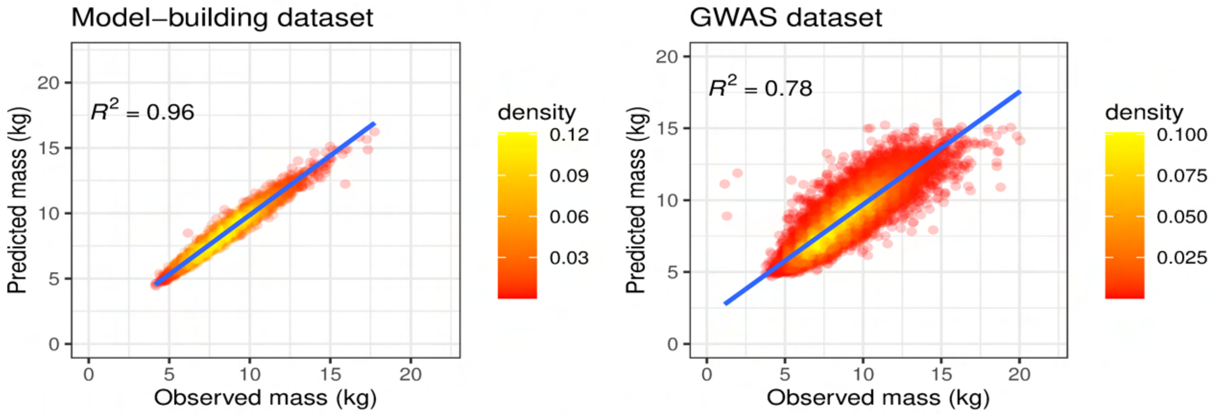

(c) Gynoid Total Mass

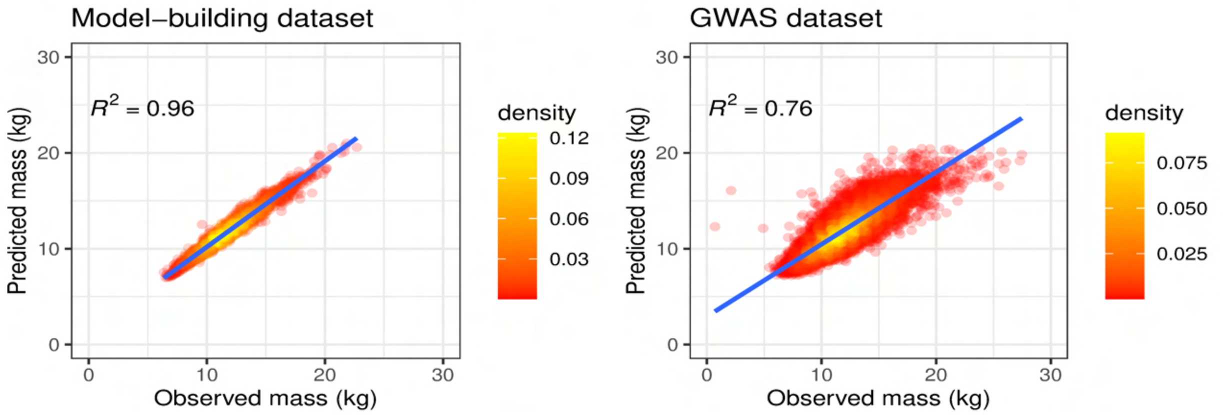

(d) Total Mass

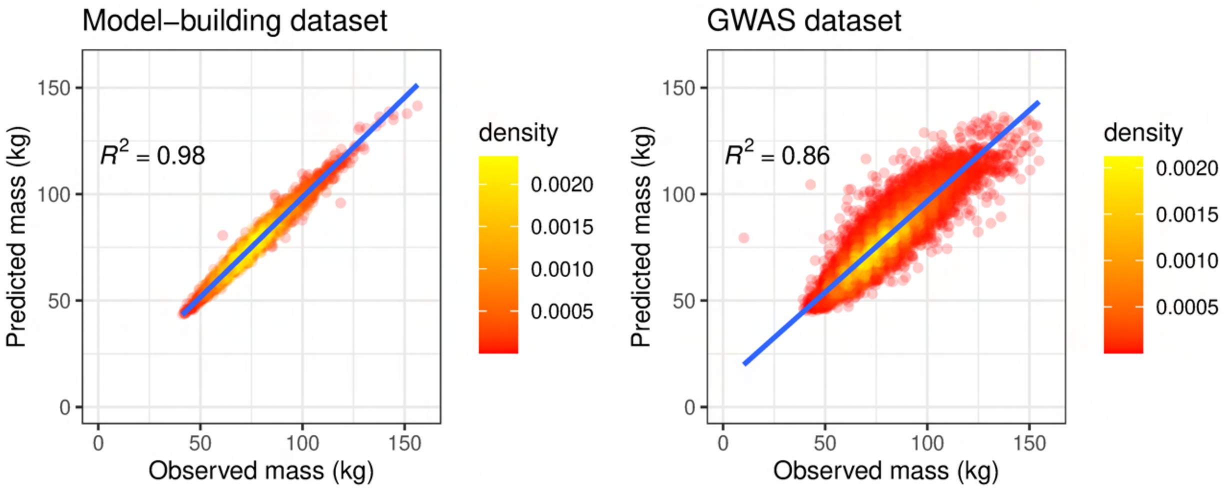

(e) Legs Total Mass

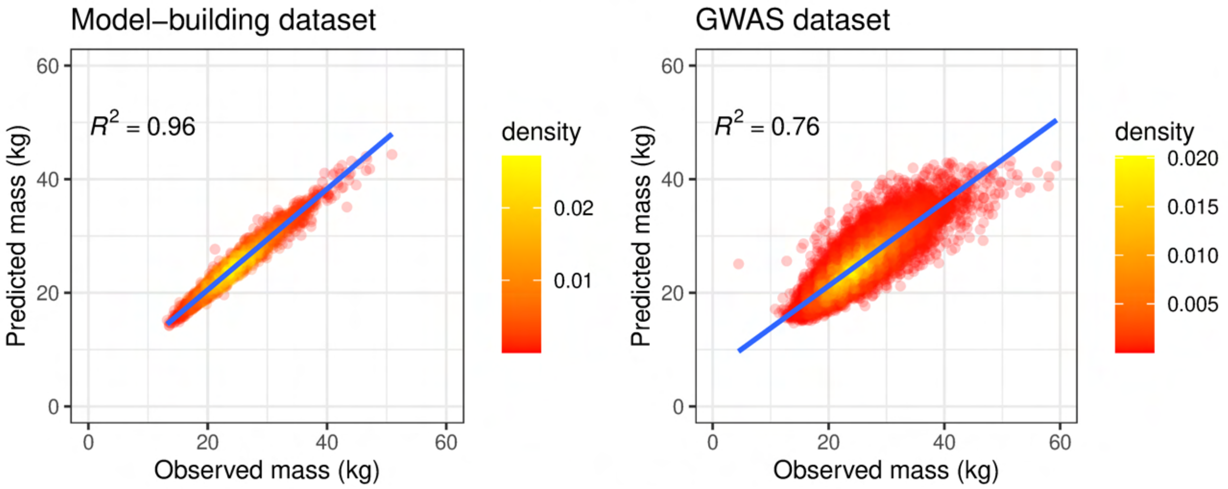

(f) Trunk Total Mass

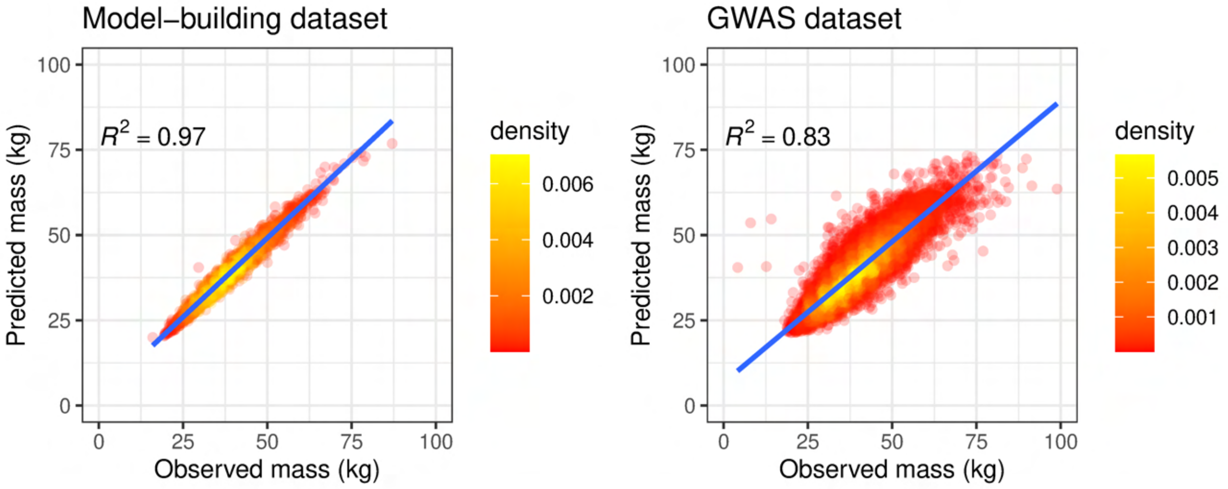

#### 10.2 Uniform QQ plots

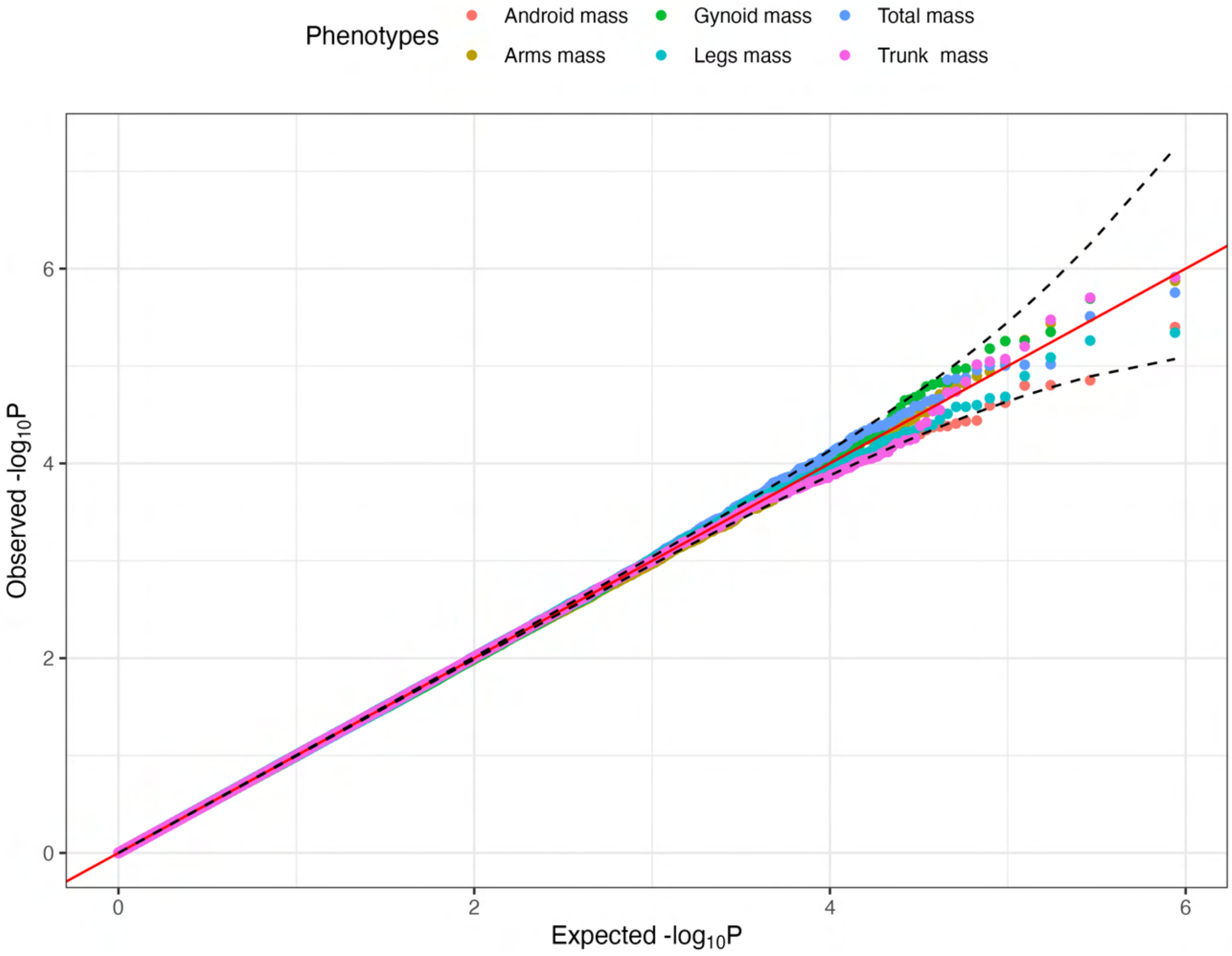

Figure 14: **Uniform quantile-quantile (QQ) plots for SynSurr GWAS of permuted UKBB DEXA phenotypes.**

#### 10.3 Miami plots

Figure 15: **Miami plots for SynSurr and Standard GWAS for the UKBB DEXA phenotypes.** SynSurr GWAS results are shown in the upper panel. Standard GWAS results based on only the observed data are shown in the lower panel. The red solid genome-wide significant lines are drawn at $-\log_{10}(5 \times 10^{-8})$.

(a) Android Total Mass

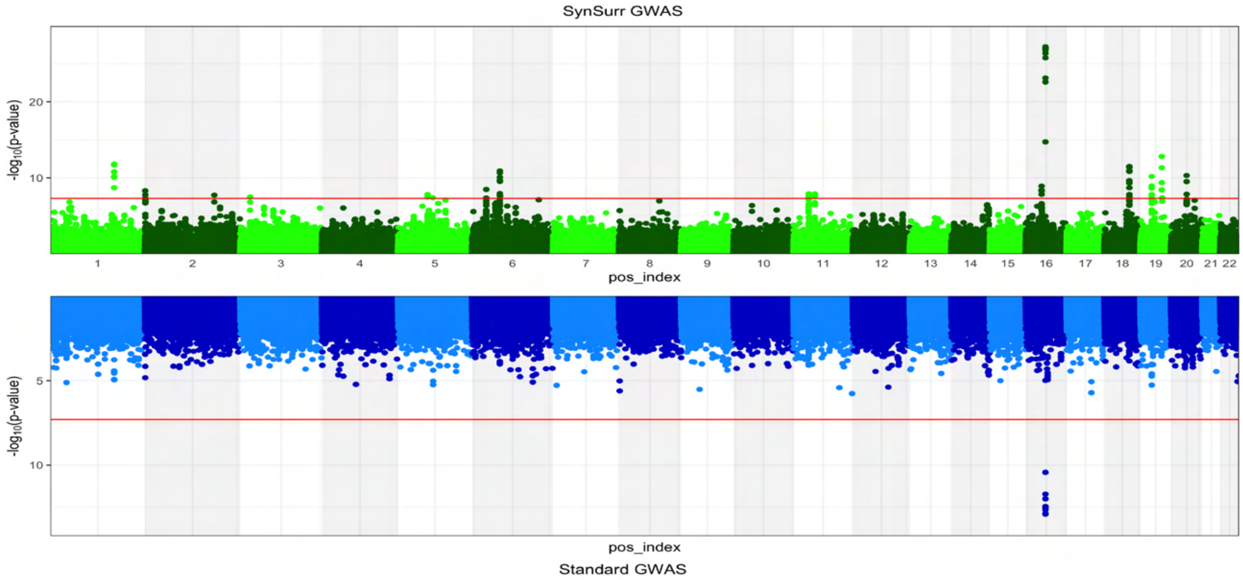

(b) Arms Total Mass

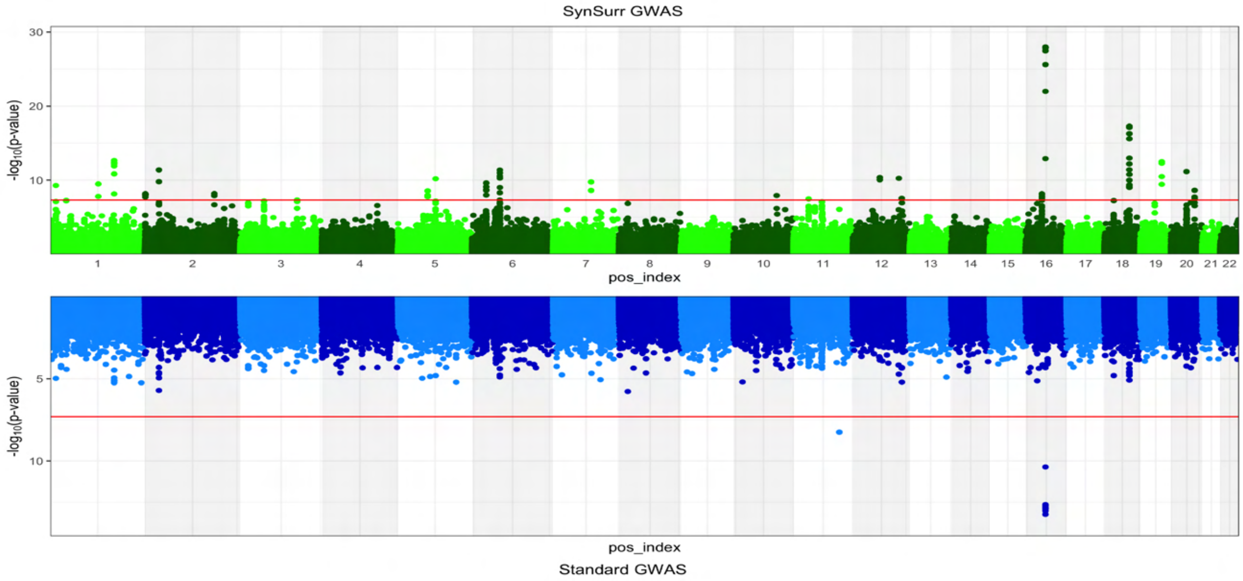

(c) Gynoid Total Mass

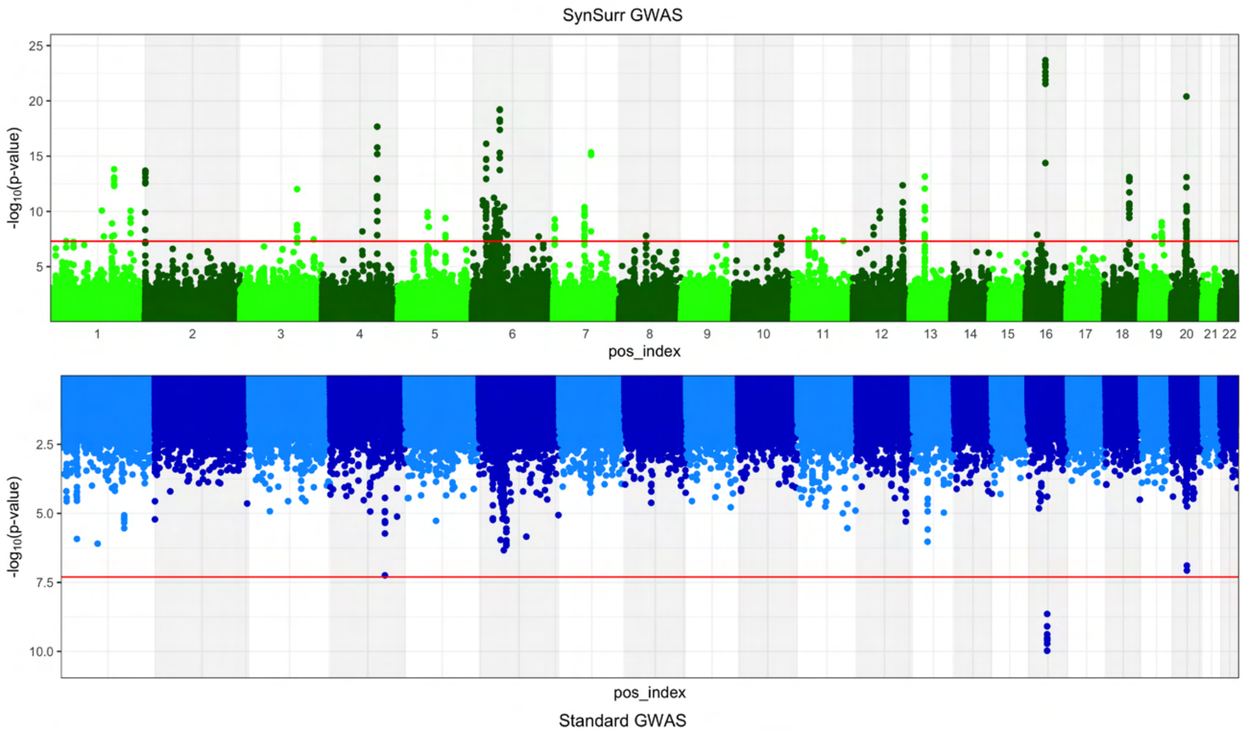

(d) Total Mass

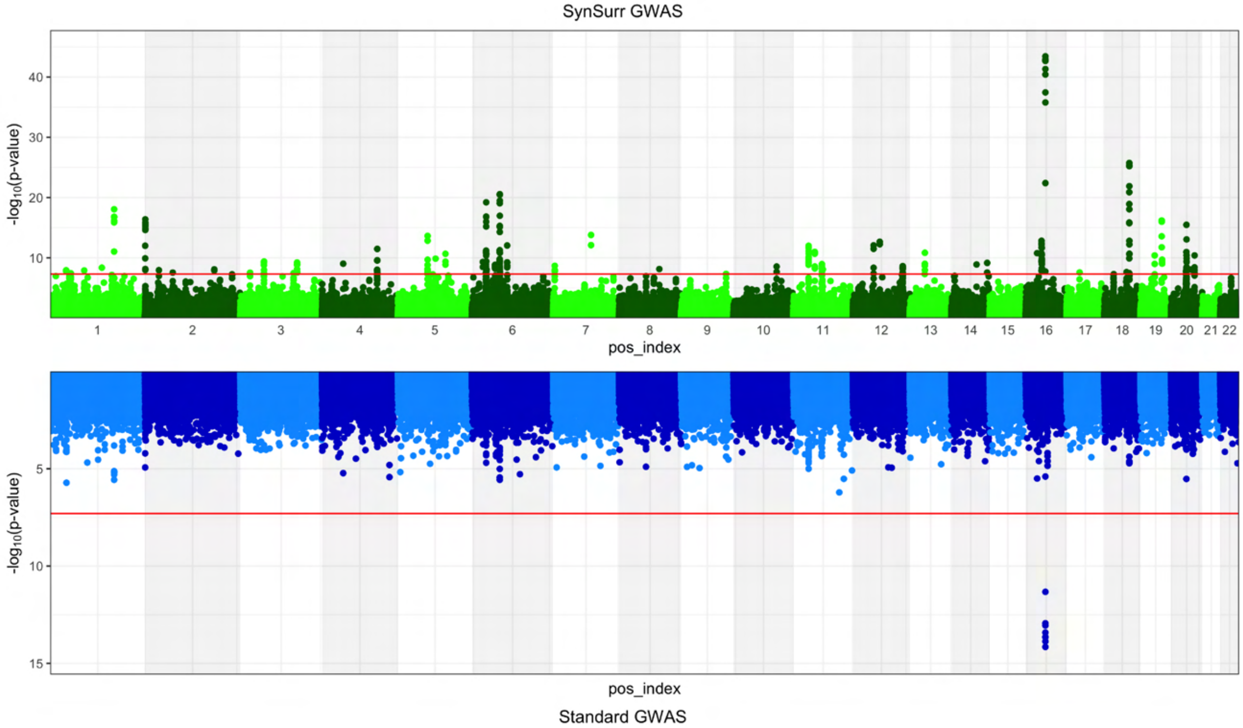

(e) Legs Total Mass

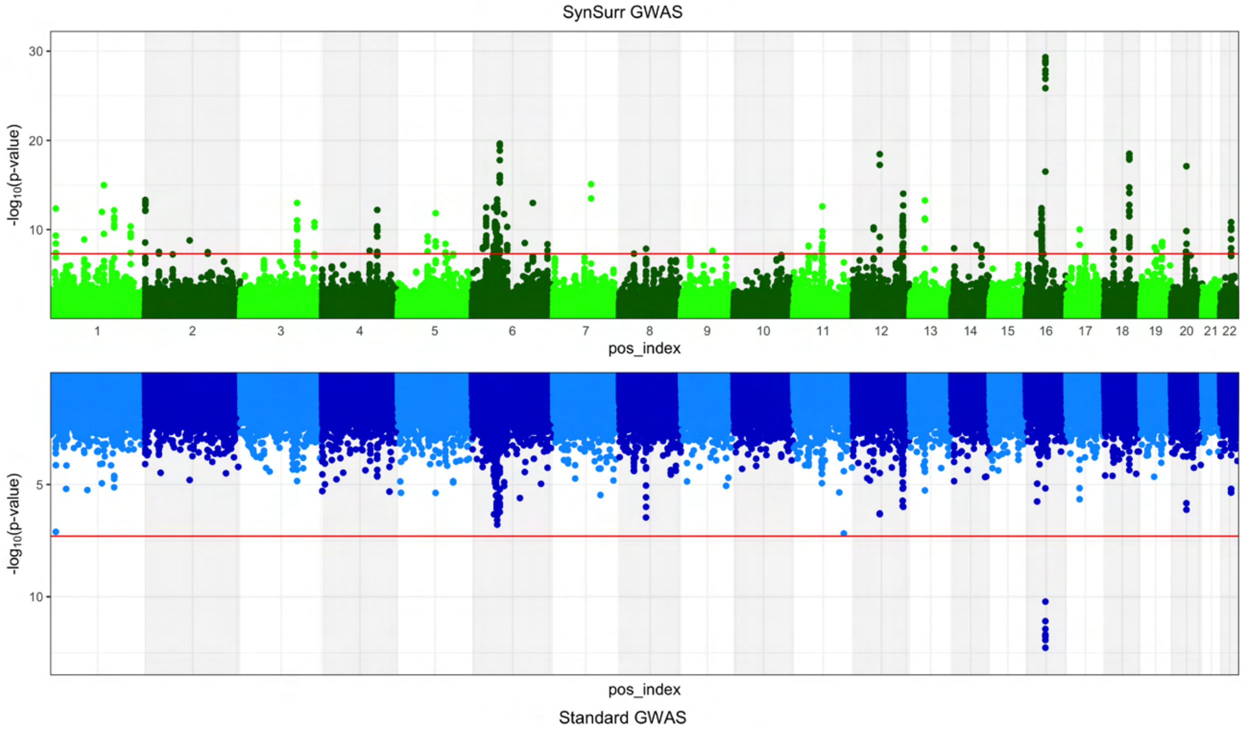

(f) Trunk Total Mass

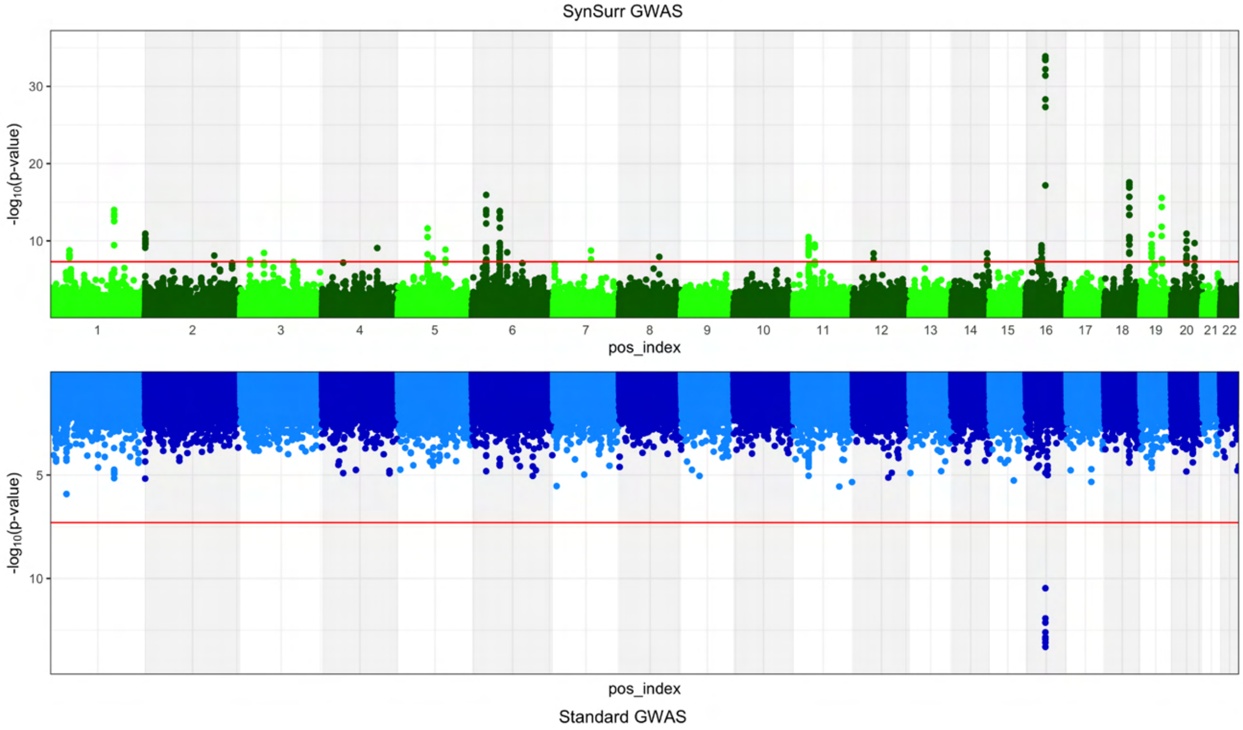

#### 10.4 Comparison of Standard and SynSurr associations

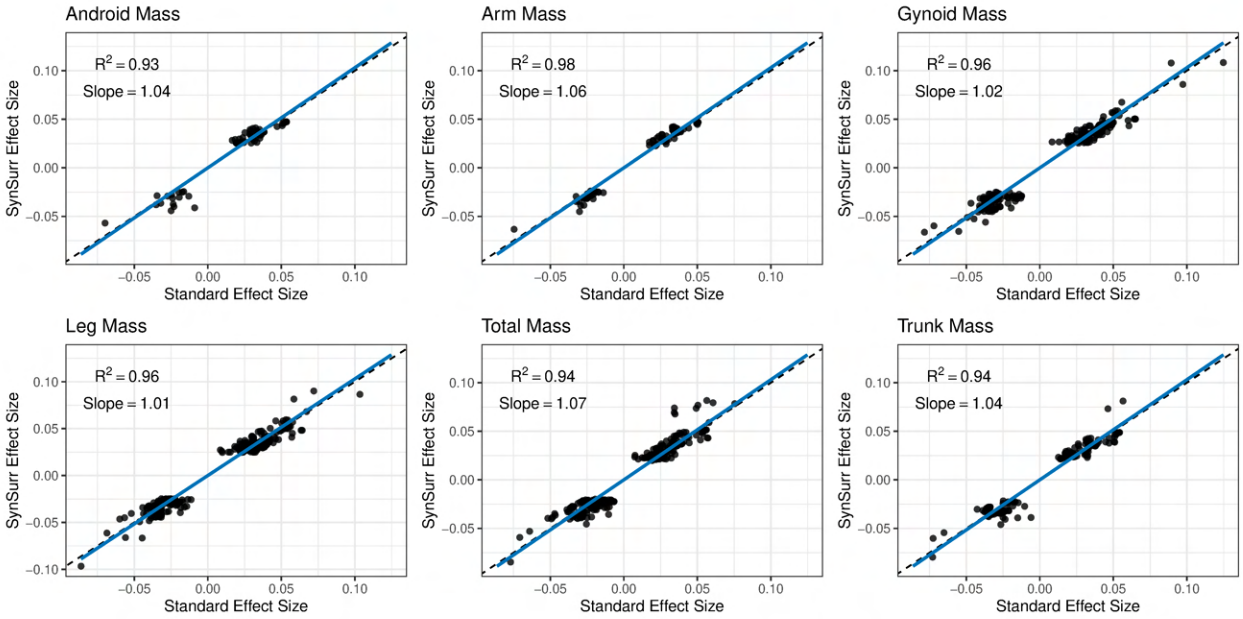

Figure 16: **Comparison of Standard and SynSurr effect sizes at genome-wide significant associations for the UKBB DEXA phenotypes.**

Table 18: **Comparison of genome-wide significant SNPs discovered by SynSurr with Standard GWAS for the UKBB DEXA phenotypes.**

| DEXA Phenotype | SynSurr Only | SynSurr and Standard | Standard Only |
|---|---|---|---|
| Android | 65 | 8 | 0 |
| Arms | 80 | 9 | 1 |
| Gynoid | 252 | 8 | 0 |
| Legs | 270 | 8 | 0 |
| Total | 268 | 8 | 0 |
| Trunk | 142 | 8 | 0 |

#### 10.5 Split sample study

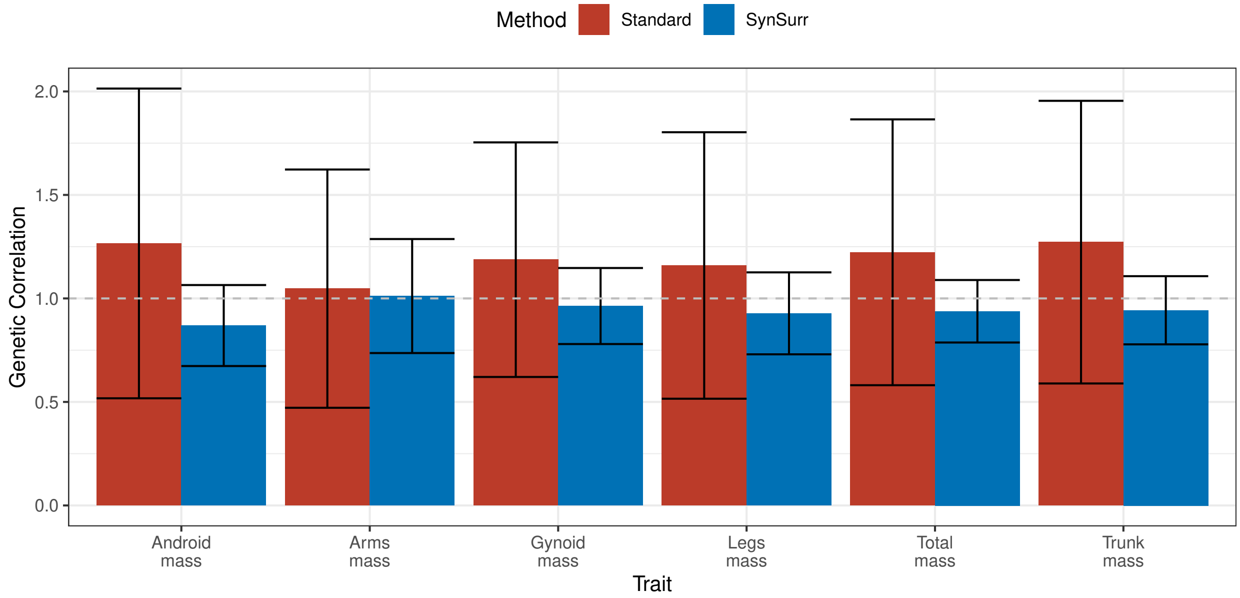

Figure 17: **Genetic correlation between the effect sizes estimated in the discovery and replication cohorts.** The discovery cohort consisted of 80% of the GWAS data, stratifying by observation of the target outcome, and the replication cohort consisted of the remaining 20% of the GWAS data. Genetic correlation was estimated from summary statistics by linkage-disequilibrium score regression (LDSC). Error bars are 95% confidence intervals.

Table 19: **Number of genome-wide significant SNPs in the discovery and replication cohorts by Standard GWAS and SynSurr.** The discovery cohort consisted of 80% of the GWAS data, stratifying by observation of the target outcome, and the replication cohort consisted of the remaining 20% of the GWAS data.

| Phenotype | Standard Discovery | Standard Replication | SynSurr Discovery | SynSurr Replication |
|---|---|---|---|---|
| Android | 8 | 8 (100%) | 22 | 17 (77%) |
| Arms | 8 | 8 (100%) | 38 | 34 (89%) |
| Gynoid | 8 | 6 (75%) | 129 | 101 (78%) |
| Legs | 9 | 7 (78%) | 181 | 109 (60%) |
| Total | 8 | 8 (100%) | 155 | 126 (81%) |
| Trunk | 8 | 8 (100%) | 76 | 56 (73%) |

### References

1. Meng, X.-L. & Rubin, D. B. Maximum likelihood estimation via the ECM algorithm: A general framework. *Biometrika* **80,** 267–278 (1993).
2. McCaw, Z. R., Gaynor, S. M., Sun, R. & Lin, X. Leveraging a surrogate outcome to improve inference on a partially missing target outcome. *Biometrics* **Online ahead of print** (2022).
3. McCaw, Z. R. *SurrogateRegression: Surrogate Outcome Regression Analysis* Comprehensive R Archive Network (2020). `https://CRAN.R-project.org/package=SurrogateRegression`.
4. R Core Team. *R: A Language and Environment for Statistical Computing* R Foundation for Statistical Computing (Vienna, Austria, 2022). `https://www.R-project.org/`.
5. Cox, D. Some Remarks on Likelihood Factorization. *Lecture Notes-Monograph Series* **36,** 165–172 (2001).
6. Casella, B. & Berger, R. *Statistical Inference.* 2nd ed. (Duxbury/Thomson Learning, Pacific Grove, CA, 2002).
7. Little, R. J. & Rubin, D. B. *Statistical Analysis with Missing Data* 2nd (John Wiley & Sons, 2002).
